# Synergistic Glycerol Coutilization Promotes Metabolic Fitness of *Saccharomyces cerevisiae* on Lactate

**DOI:** 10.1101/2025.03.27.645806

**Authors:** Wasti Nurani, Atit Kanti, Obie Farobie, Widya Fatriasari

**Affiliations:** Research Center for Biomass and Bioproducts, National Research and Innovation Agency (BRIN), Cibinong, West Java, Indonesia; Research Center for Biosystematics and Evolution, National Research and Innovation Agency (BRIN), Cibinong, West Java, Indonesia; Department of Mechanical and Biosystem Engineering, Faculty of Agricultural Engineering and Technology, IPB University (Bogor Agricultural University), Bogor, West Java, Indonesia; Surfactant and Bioenergy Research Center (SBRC), IPB University (Bogor Agricultural University), Bogor, West Java, Indonesia

## Abstract

Wet waste-derived lactic acid is a promising alternative carbon source for yeast-based biorefineries, but the knowledge and demonstration of its utilization are lacking. Using 22 natural isolates of *Saccharomyces cerevisiae* and two non-conventional yeasts as the experimental model, here we show that there is huge inter- and intra-species variability among yeasts in terms of their lactate utilization capacity. Interestingly, in agreement with the underlying theoretical framework, glycerol cofeeding at a molar ratio of 0.1–0.5 is often necessary and sufficient to substantially improve the utilization efficiency in *S. cerevisiae*, i.e., up to 94% of its maximum theoretical yield relative to glucose. The synergistic effect of glycerol cofeeding is modulated by the precultivation medium—i.e., whether the cells were previously grown on glucose or acetate—and the glycerol/lactate molar ratio applied. Specifically, a higher molar ratio (up to 2.5) often leads to saturation or even antagonism. Using the knowledge of *S. cerevisiae* metabolism, potential mechanistic underpinnings of the observed phenomena are given to guide future metabolic engineering endeavors.

## Introduction

The management of high-strength organic wet waste, which includes food waste, wastewater sludge, livestock manure, and inedible fats, oils, and grease (FOG) ^1^, is problematic and costly ^2,3^. Methane-arrested anaerobic digestion (MAAD) may provide financial incentives by transforming them into volatile fatty acids (VFAs or carboxylates), which, among others, can be catalytically upgraded into jet-range aromatics and paraffins ^4,5^. Its main drawback is the substantial generation of lactic acid, which is too short for the catalytic platform ^6^.

The potential amount of wet waste-derived lactic acid is comparable to that of agriculture-based sugars (Note S1). Although *pure* lactic acid has commercial value, the dilute-, impure-, and potentially racemic-status of the *wet waste-derived* lactic acid stream will lead to an expensive recovery/purification process. Arguably, it is more strategic to use the acid as a carbon source for microbial cultures. Cell biomass is relatively easy to recover from the broth (i.e., through solid/liquid separation) and can be converted into a highly valued single-cell protein (SCP, e.g., as animal feed) ^7,8^. Furthermore, depending on the microbe, the acid can also be transformed into secondary metabolites such as drop-in- and performance-advantaged-transportation fuel additives (Figure S1).

The baker’s yeast *Saccharomyces cerevisiae* has been commonly used to produce SCP ^9^, yeast extract ^10^, and non-native secondary metabolites ^11^. Unfortunately, *S. cerevisiae* generally grows poorly on lactate ^12–14^ (Note S2, Table S3). Interestingly, glycerol cofeeding at a 3.08 molar ratio reportedly improved the total growth of *S. cerevisiae* CEN.PK113-7D on lactate by ∼25% and the total growth rate by ∼2.6 times ^14^ (Figure S2). As such, we wondered whether the same benefits could also be experienced by other yeasts, preferably at a much lower glycerol/lactate molar ratio.

To answer this question, we extended the previous study (Tables S4-S8, Figures S3-S9). We first characterized the growth of 22 natural *S. cerevisiae* isolates on lactate alone (i.e., in a 2%-lactate/urea minimal medium termed **YM-25**) using an equivalent 2%-glucose/urea minimal medium termed **YM-21** as the reference. We then characterized their growth in three different **YM-25 + GLY** media, each containing either 0.2, 1.0, or 5.0 % w/v glycerol (resulting in a glycerol/lactate molar ratio of 0.1, ∼0.5, or ∼2.5, respectively). In parallel, we also characterized their growth in the three corresponding **YM-GLY** media (which solely contained glycerol at 0.2, 1.0, or 5.0 % w/v). To provide a positive control for glycerol growth and a negative control for lactate growth, two non-*S. cerevisiae* strains, *Clavispora lusitaniae* (Y1728) and *Wickerhamiella versatilis* (Y1125), were characterized alongside. To obtain biological replicates, and more importantly, to investigate whether the synergism was a genuine “state function” of glycerol/lactate coutilization or a “path function” (*viz*., precultivation-dependent phenotype), we precultivated each strain on two different agar plates containing either **YM-10** (i.e., 2%-acetate/urea minimal medium) or commercial potato infusion/dextrose (**PDA**). Potato infusion served as a vegan and cheaper alternative to the commonly used yeast extract/peptone basal medium. The use of glucose as a carbon source and the richness of nutrients provided by potato infusion were expected to ensure optimal inoculum growth. In contrast, the acetate growth provided by YM-10 was expected to provide a less repressive metabolic landscape that may be more accommodating to growth on lactate and/or glycerol. Both PDA and YM-10 agar contained neither lactate nor glycerol; thus, they were nutritionally distinct and theoretically unbiased from the experimental media, as intended. The agar format was chosen to ensure the maximum exposure of the cells to atmospheric oxygen—something not granted by the liquid format. The resulting growth yield was then mathematically analyzed to assess if the glycerol cofeeding effect was synergistic or antagonistic. Finally, the residual lactate abundance of some representative cultures was quantified to assess for any perturbed lactate uptake.

## Results

### Growth on lactate

The ability of each strain to grow on lactate was assessed based on the percentage of its final OD_600_ in YM-25 relative to that in YM-21 (Figure 1A). Six out of the 21 *S. cerevisiae* strains studied (29%) were able to grow reasonably well on lactate after 5-6 days of incubation (Figures 1AB, hereafter “Lactate +” strains), with a final OD_600_ percentage ranging between 14-73% depending on the strain and precultivation medium. (Y93 was excluded from the statistical analysis since the strain grew atypically poorly on glucose: Tables S5-S9, Figure S9). The growth of *C. lusitaniae* (Y1728), however, excelled even the best *S. cerevisiae* strain with a final OD_600_ percentage of approximately 82% (Figure 1A). Similar to the majority of *S. cerevisiae* strains, no meaningful growth was observed in *W. versatilis* (Y1125) (Figure 1A).

**Figure 1.**
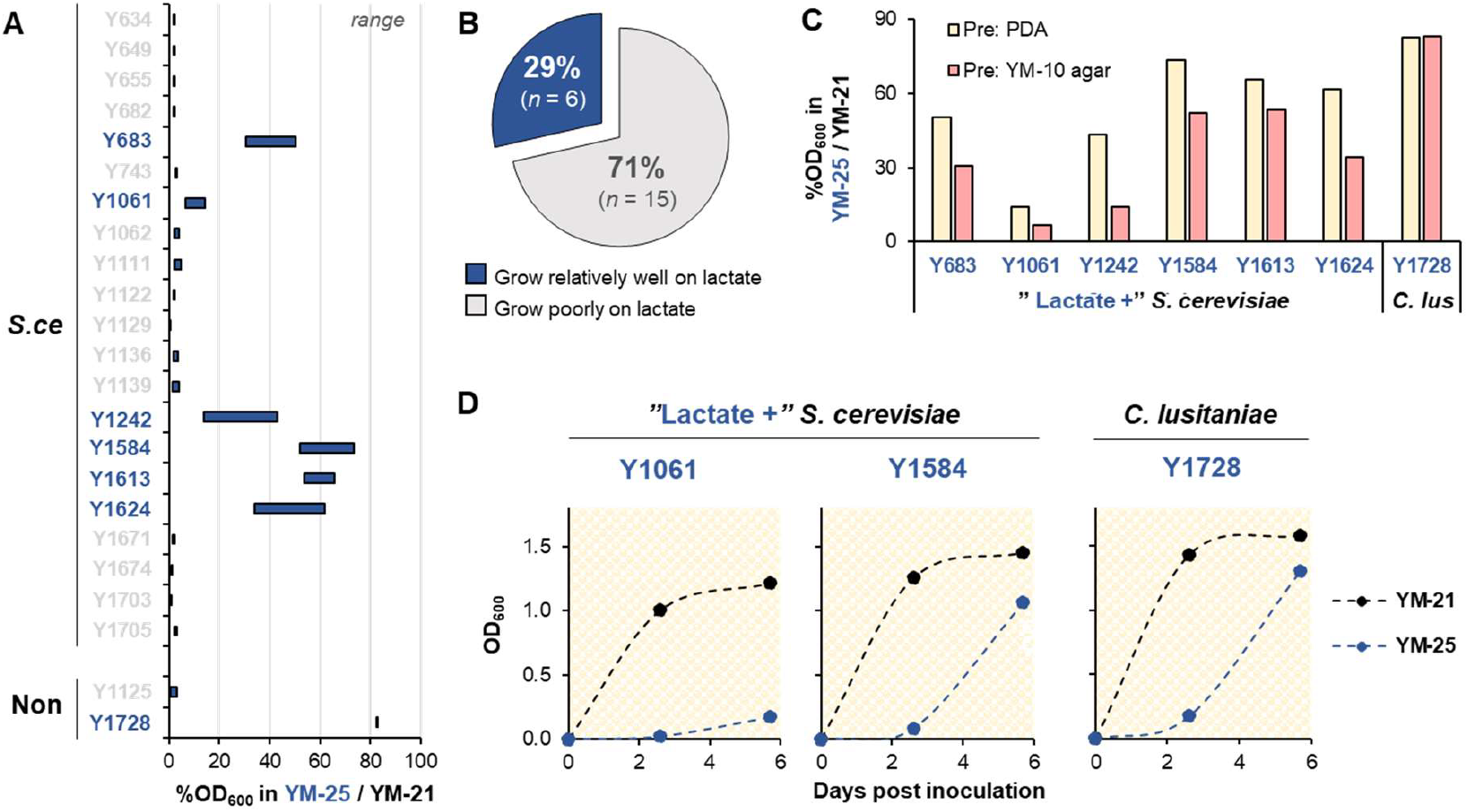
Growth profiles of 21 *S. cerevisiae* and 2 non-*S. cerevisiae* strains in YM-25 relative to their counterparts in YM-21. **A**, OD_600_ percentage of each strain in YM-25 relative to that in YM-21. *S*.*ce*: *S. cerevisiae*. Non: non-*S. cerevisiae*. The “range” on the top right side indicates that each bar encompasses the data of cultures precultivated on PDA and YM-10 agar, ranging from the smallest value to the highest value. **B**, Percentage of *S. cerevisiae* strains that either grew relatively well or poorly on lactate. *n*: number of strains. **C**, OD_600_ percentage of the seven strains considered able to grow relatively well on lactate as a function of precultivation medium (“Pre”). *C. lus*: *Clavispora lusitaniae*. **A**-**C**, Source data: Table S9. **D**, Time-course OD_600_ evolution of the three relatively-well lactate grower representatives in YM-21 and YM-25. The yellow background indicates precultivation on PDA. The profiles of the other four strains are presented in Figure S10. Source data: Table S10. **A**-**D**, Blue font indicates strains that grow relatively well on lactate based on the bar height.

In contrast to the lactate growth ability of *C. lusitaniae*, that of *S. cerevisiae* was sensitive to the precultivation medium. For all the six “Lactate +” strains, precultivation on YM-10 agar led to substantially poorer growth on lactate than the precultivation on PDA did (Figure 1C). The statistical analysis in Figure 3C (left) indicated that the differences were significant.

Time-course OD_600_ measurement of cultures precultivated on PDA suggested that both “Lactate +” *S. cerevisiae* and *C. lusitaniae* experienced an apparent lag period when transferred to YM-25 (Figure 1D). This lag period was absent in their counterparts in YM-21.

### Growth on glycerol

The ability of each strain to grow solely on glycerol in a minimal medium context (i.e., in each of the YM-GLY media series) was also assessed relative to their growth on glucose in YM-21 (Figure 2A). Irrespective of the precultivation medium and the specific glycerol concentration where growth was displayed, around 38% of the 21 *S. cerevisiae* strains tested (i.e., 8 strains) at least once displayed a final OD_600_ percentage between 3-33% of that on glucose, whereas 57% (12 strains) between 33-63% (Figure 2B). In other words, almost all the *S. cerevisiae* strains tested at least once demonstrated an ability to grow on glycerol. However, the growth of *S. cerevisiae* on glycerol was rather sporadic and did not exhibit the logical dose-dependent response. In fact, the final OD_600_ percentage of most strains in the glycerol minimal media typically started at a near-zero value (Figure 2A), indicating that at least once no noticeable growth was detected even after 5-6 days of cultivation. Furthermore, the highest values did not always correspond to the highest (5.0% w/v) glycerol concentration (Figure 2C). Instead, 18% of the 22 *S. cerevisiae* strains tested displayed the highest OD_600_ percentage when cultivated in the lowest (0.2% w/v) glycerol concentration, and 59% of the strains grew best at two different glycerol concentrations depending on the precultivation medium. Such aberrant growth was in contrast with that of the two non-*S. cerevisiae* strains, in which the lowest percentage OD_600_ was at least 42% and corresponded to the lowest glycerol concentration in the respective YM-GLY media series (Figures 2AD, S11). Additionally, regardless of the precultivation medium, both strains grew better in 5% w/v glycerol than in 2% w/v glucose, with a final OD_600_ percentage of approximately 120% (Figure 2A).

**Figure 2.**
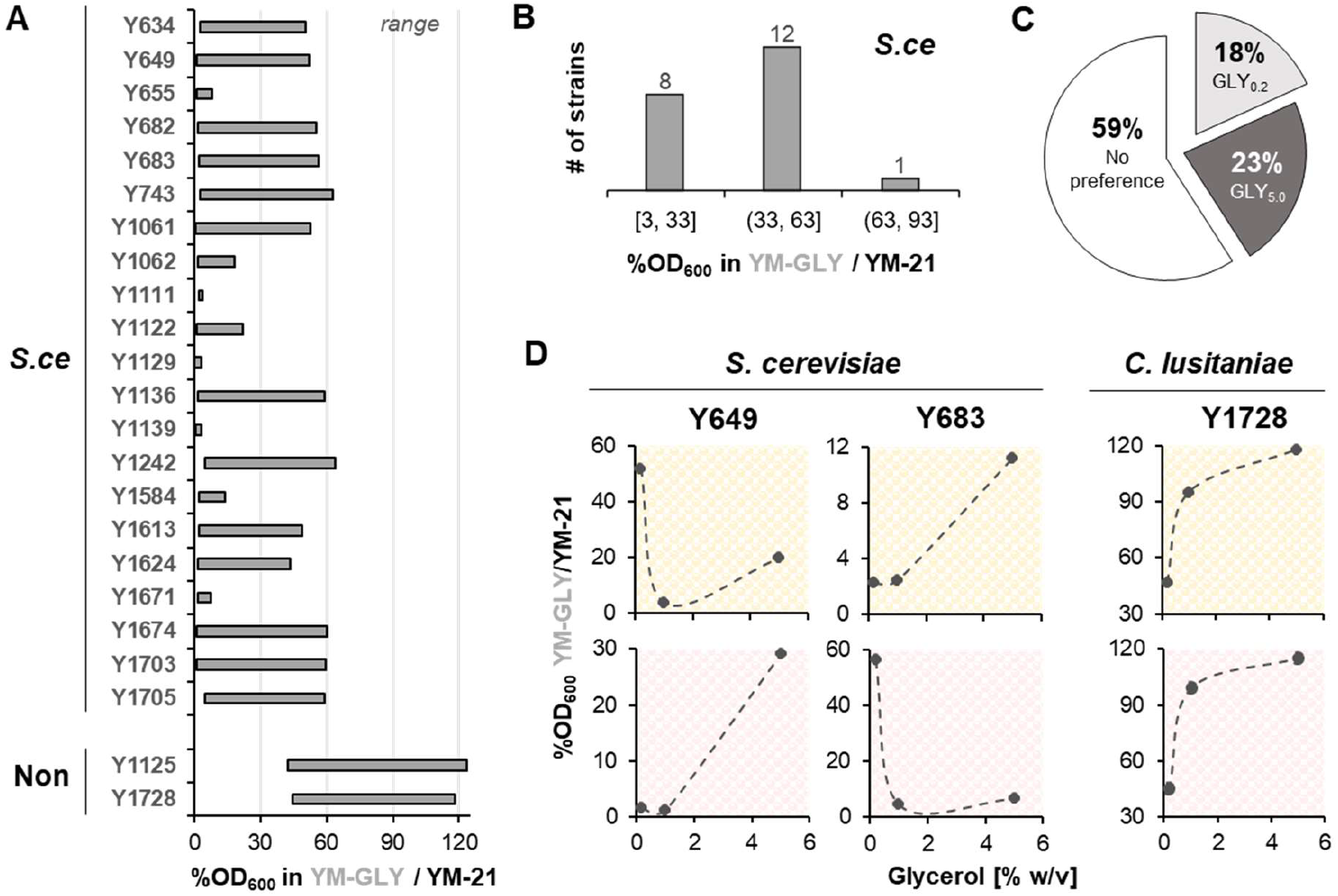
Growth profiles of 21 *S. cerevisiae* and 2 non-*S. cerevisiae* strains in the three variants of YM-GLY relative to their counterpart in YM-21. **A**, OD_600_ percentage of each strain in each YM-GLY variant relative to that in YM-21. *S*.*ce*, Non: see Figure 1. The “range” indicates that each bar encompasses the data of cultures precultivated on PDA and YM-10 agar, in each of the three glycerol concentration variants, ranging from the smallest value to the highest value. **B**, Number of *S. cerevisiae* (“*S*.*ce*”) strains described in (**A**) exhibiting OD_600_ percentage within the indicated range. **C**, Percentage of *S. cerevisiae* strains that reached their maximum OD_600_ at a glycerol concentration of 0.2% w/v (GLY_0.2_) or 5.0% w/v (GLY_5.0_) versus those that did not prefer any specific concentration for reaching their maximum OD_600_. Analysis and source data: Tables S12 and S13. The graphical rendering of each strain’s preference: Figure S11. **D**, OD_600_ percentage of three representative strains in the three YM-GLY variants relative to that in YM-21 as a function of initial glycerol concentration. The yellow background indicates precultivation on PDA and pink on YM-10 agar. The profiles of other strains: Figure S12. **A, B, D**: Source data: Table S11.

### Growth on glycerol/lactate mixtures

To determine how glycerol supplementation affected the growth of all strains on lactate, the OD_600_ percentage in each of the YM-25 + GLY media relative to their counterparts in YM-21 was calculated. Upon sorting the maximum OD_600_ percentage of all PDA-precultivated strains in increasing order, two distinct groups of “Lactate –” *S. cerevisiae* strains emerged: the one whose value was similar to that of their YM-10-agar precultivated counterpart and centered around 15% (hereafter, “Lactate – (15)” strains) and the other whose value was substantially higher than their YM-10 agar-precultivated counterpart’s and centered around 70% (hereafter, “Lactate – (70)” strains) (Figure 3A).

**Figure 3.**
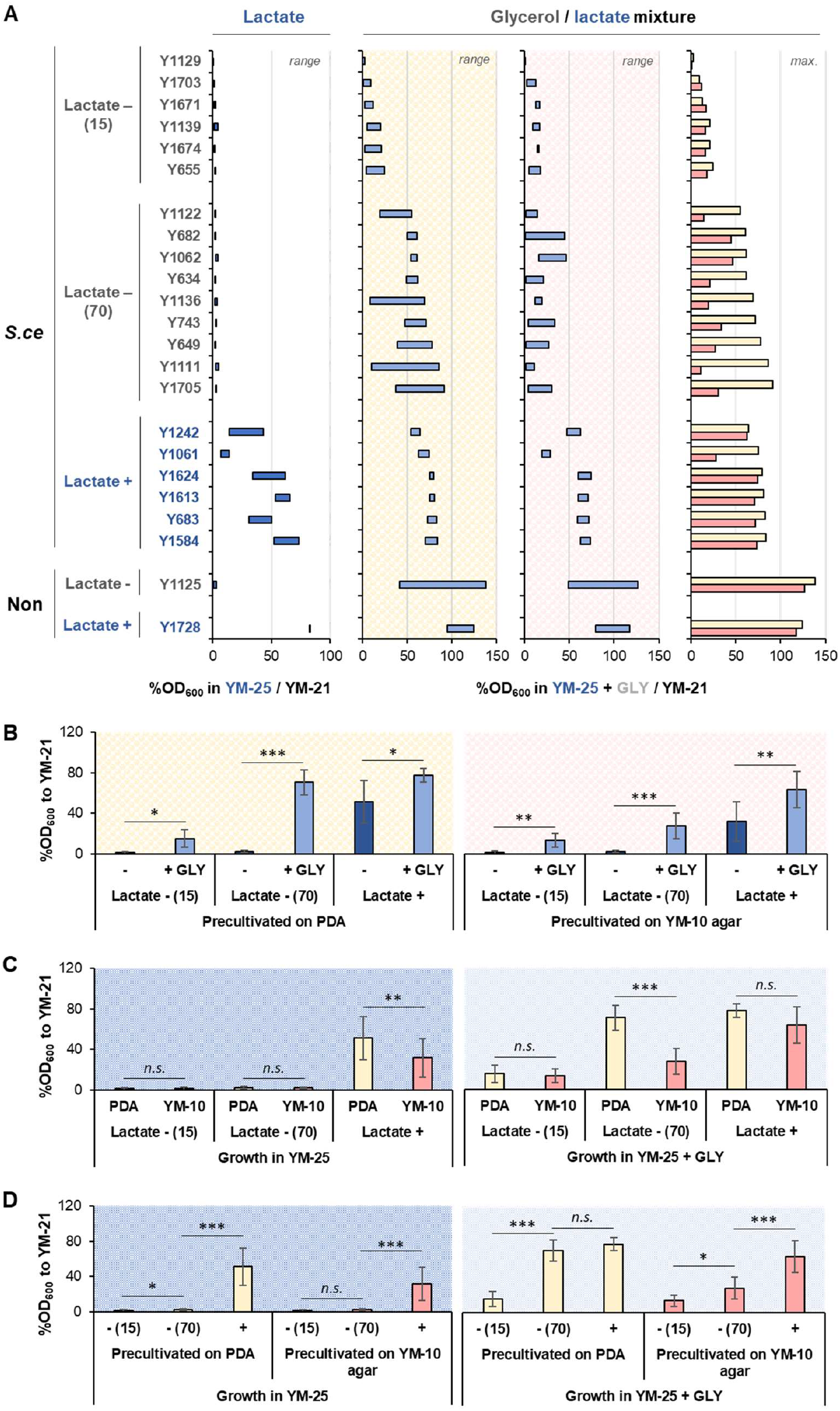
Growth profiles of 21 *S. cerevisiae* and 2 non-*S. cerevisiae* strains in three variants of YM-25 + GLY relative to their counterparts in YM-21 and YM-25. **A**, OD_600_ percentage of each strain in each YM-25 + GLY variant relative to that in YM-21. *S*.*ce*, Non: see Figure 1. The first bar chart on the left-hand side is the rearranged version of Figure 1A. The “range” indicates that each bar encompasses the data of cultures in each of the three glycerol concentration variants, ranging from the smallest value to the highest value. The “max” indicates that each bar displays the maximum value of the respective dataset. Lactate – (15) and Lactate – (70): see text. Source data: Table S14. **B**-**D**, The statistical analysis of the differences in OD_600_ percentage to YM-21 as a function of glycerol supplementation (**B**), precultivation medium (**C**), and genetic background (**D**). Each bar represents the mean ± standard deviation. *n*.*s*.: not statistically significant (*p*-value ≥ 0.05). *: *p*-value < 0.05. **: *p*-value < 0.005. ***: *p*-value < 0.001. Source data and statistical analysis: Tables S15-S18. **A**-**D**, Yellow and pink background: see Figure 2. The dark blue background indicates growth in YM-25 and light blue in YM-25 + GLY.

Given the observations, it is evident that there were at least three variables affecting the growth of *S. cerevisiae* on lactate: in addition to glycerol supplementation (variable #1) , the precultivation medium (variable #2) and genetic background (i.e., whether the strain was “Lactate +”, “Lactate – (70)”, or “Lactate – (15)”; variable #3) also play a determining role.

Statistical analysis indicated that (*i*) glycerol supplementation significantly improved the growth of all *S. cerevisiae* strains tested, regardless of the precultivation medium and genetic background (Figure 3B); (*ii*) precultivation on PDA led to a significantly higher biomass yield relative to the precultivation on YM-10 agar, but only to “Lactate +” strains cultivated in YM-25 and “Lactate – (70)” strains cultivated in YM-25 + GLY (Figure 3C); and (*iii*) the “Lactate +” genotype resulted in a growth advantage relative to the two “Lactate –” genotypes in YM-25 (regardless of the precultivation medium) and also in YM-25 + GLY, but only when they were precultivated on YM-10 agar (Figure 3D). When the Lactate – (70) strains were precultivated on PDA and cultivated in YM-25 + GLY, the biomass yield was not statistically different from that of the “Lactate +” (Figure 3D).

To mathematically assess the nature of interactions between glycerol- and lactate-utilizations, the difference between the observed OD_600_ value and the theoretical additive OD_600_ value (i.e., the sum of OD_600_ values on pure substrates at the matching concentration) was calculated. A positive-value difference was interpreted as synergism, whereas a negative-value difference as antagonism (Equations 1-4, Figure 4A).

**Figure 4.**
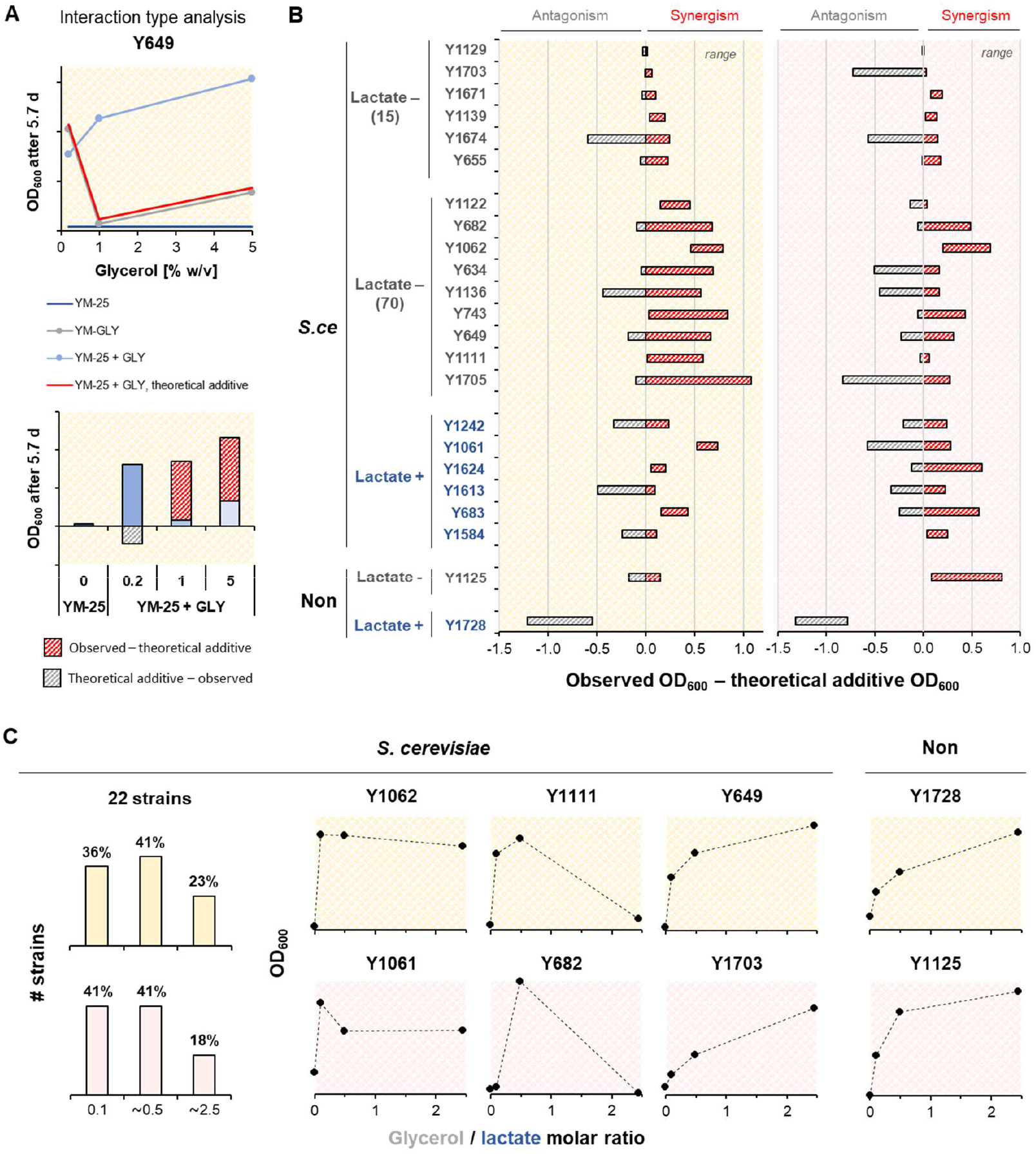
Analysis of interactions between glycerol- and lactate-utilizations based on the growth profiles of each strain in YM-25, YM-GLY, and YM-25 + GLY. **A**, Illustration of the analysis process (see text for details). The graphs of other strains: Figure S13. **B**, Range of differences between the observed and the theoretical additive OD_600_ exhibited by the respective strains when cultivated in each of the three YM-25 + GLY variants. range: see Figure 3. Source data: Table S19. **C**, OD_600_ as the function of the initial glycerol/lactate molar ratio of the medium. Bar charts on the left-hand side indicate the percentage of strains that reached their maximum OD_600_ value in the respective medium. “22 strains”: the total number of strains considered in the calculation. Source data for the bar chart: Table S20. The scatter plots of other strains: Figure S14. **A**-**C**, Yellow and pink background: see Figure 2.

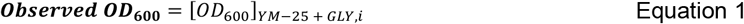

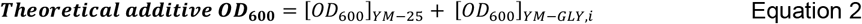

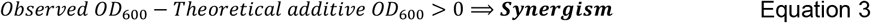

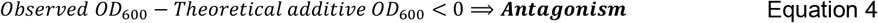

The quantitative analysis indicated that all *S. cerevisiae* strains tested (including Y93), irrespective of their natural lactate growth capacity, displayed at least one instance of synergism across the six events tested (i.e., two precultivation media × three glycerol/lactate molar ratios) (Figure 4B). Synergism was also displayed by the “Lactate -” *W. versatilis* (Y1125), and the effect was more pronounced upon precultivation on YM-10 agar (Figure 4B). In contrast, no synergism was displayed by the “Lactate +” *C. lusitaniae* (Y1728) regardless of the precultivation medium; instead, the cofeeding resulted in striking antagonism (Figure 4B).

The plotting of the final OD_600_ to the glycerol/lactate molar ratio indicated that in nearly all (77-82%) of *S. cerevisiae* tested, regardless of the precultivation medium, the glycerol/lactate synergistic effect was saturated, meaning that after the culture reached a certain OD_600_ value, the supplementation of more glycerol did not increase the OD_600_ further (Figure 4C). Often, the maximum OD_600_ value was already achieved at the lowest (0.1) molar ratio. In fact, the three highest growth yield improvements, reaching between 86% to 94% of the maximum theoretical yield relative to glucose, were achieved by the PDA-precultivated strains at 0.1 and ∼0.5 molar ratios (Table S15 red, Note S3). The saturation was often followed by a drop in OD_600_ at a higher glycerol/lactate molar ratio irrespective of the strain being “Lactate +” or “Lactate – (70)” (Figure 4C). Interestingly, the decrease in biomass level occurred although lactate uptake was unperturbed (Figures 5AB). In contrast, the two non-*S. cerevisiae* strains did not display any saturation effect, let alone a drop in OD_600_ at higher glycerol loading (Figures 4C, S14).

**Figure 5.**
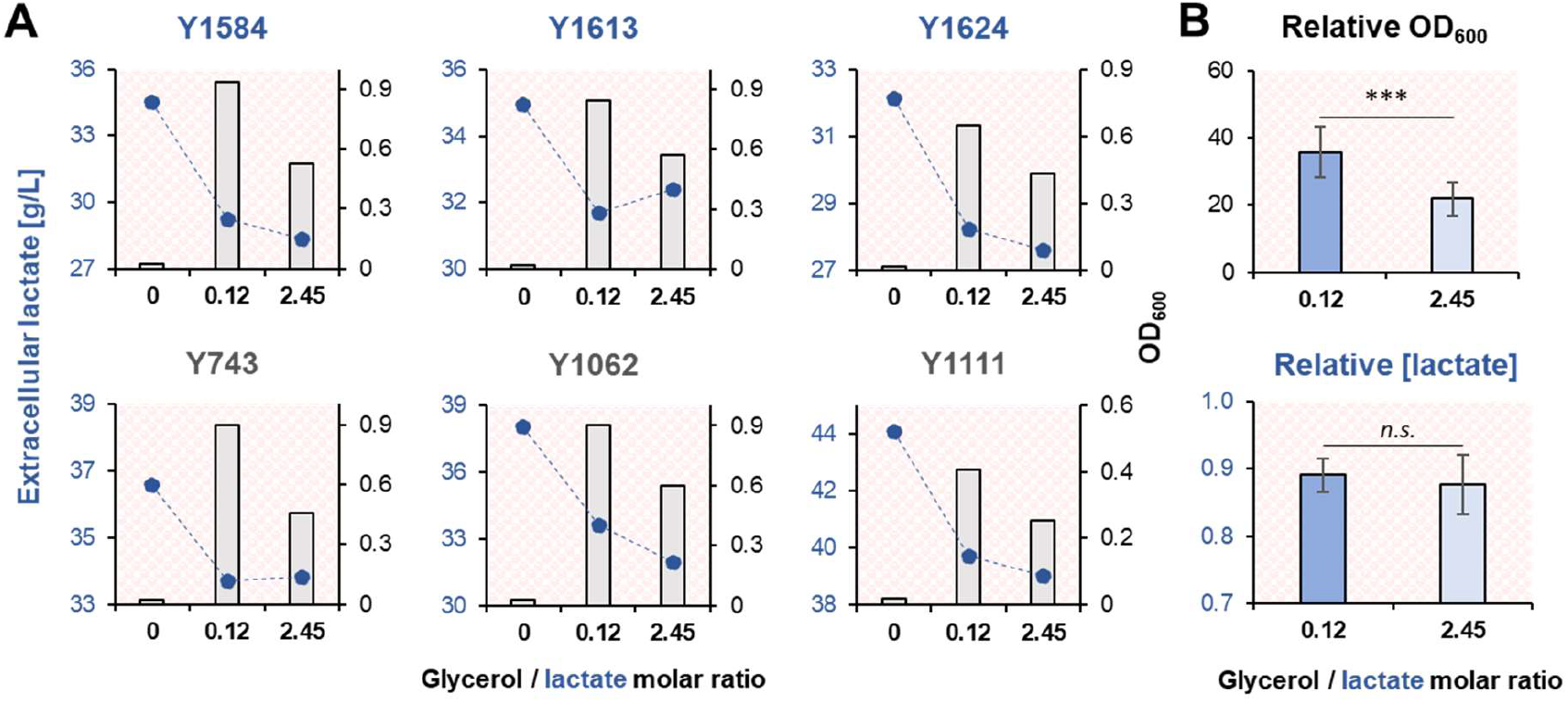
OD_600_ and residual lactate concentration in the YM-25 and YM-25 + GLY cultures of six representative *S. cerevisiae* strains. **A**, OD_600_ and residual (extracellular) lactate concentration at the day of harvest, which was before noticeable biomass amount was produced in YM-25. Top, blue font: “Lactate +” strains. Bottom, grey font: “Lactate – (70)” strains. The medium with a “0” glycerol/lactate molar ratio refers to YM-25. Pink background: see Figure 2. The observed residual lactate concentration being higher than 20 g/L (i.e., the initial lactate concentration in each medium) indicates that water evaporation had occurred over the cultivation course. **B**, Statistical analysis of the differences in OD_600_ and residual lactate concentration as the function of initial glycerol/lactate molar ratio. Each bar represents the mean ± standard deviation (*n* = 6). *n*.*s*.: not statistically significant (*p*-value ≥ 0.05). ***: *p*-value < 0.001. **A, B**, Source data and statistical analysis: Figures S15-S16, Table S22-S25.

## Discussion

Given the limited understanding of *C. lusitaniae* and *W. versatilis* metabolism, it is not possible to rationalize their growth phenotypes. This is, in fact, one of the major hurdles in the use of non-conventional yeasts as metabolic engineering chassis ^15^. In contrast, as presented below, the plethora of knowledge about *S. cerevisiae* metabolism could reasonably assist us in postulating the molecular mechanisms for the observed phenomena, thus paving the way for future metabolic engineering endeavors.

### Potential genetic basis of “Lactate +” phenotype

The postulated metabolic flux distribution of *S. cerevisiae* grown on lactate is shown in Figure 6 (sans the glycerol utilization pathway). As discussed in Section 5.4.4.1 of ^14^, the estimated ratio of the total cytosolic NADPH produced to the total biomass synthesis fluxes indicates that, compared to the growth on glucose or galactose, the growth of lactate is severely limited by cytosolic NADPH. As such, it stands to reason that given the same growth conditions, “Lactate +” strains could somehow generate more cytosolic NADPH from the same amount of lactate compared to the “Lactate -” strains.

**Figure 6.**
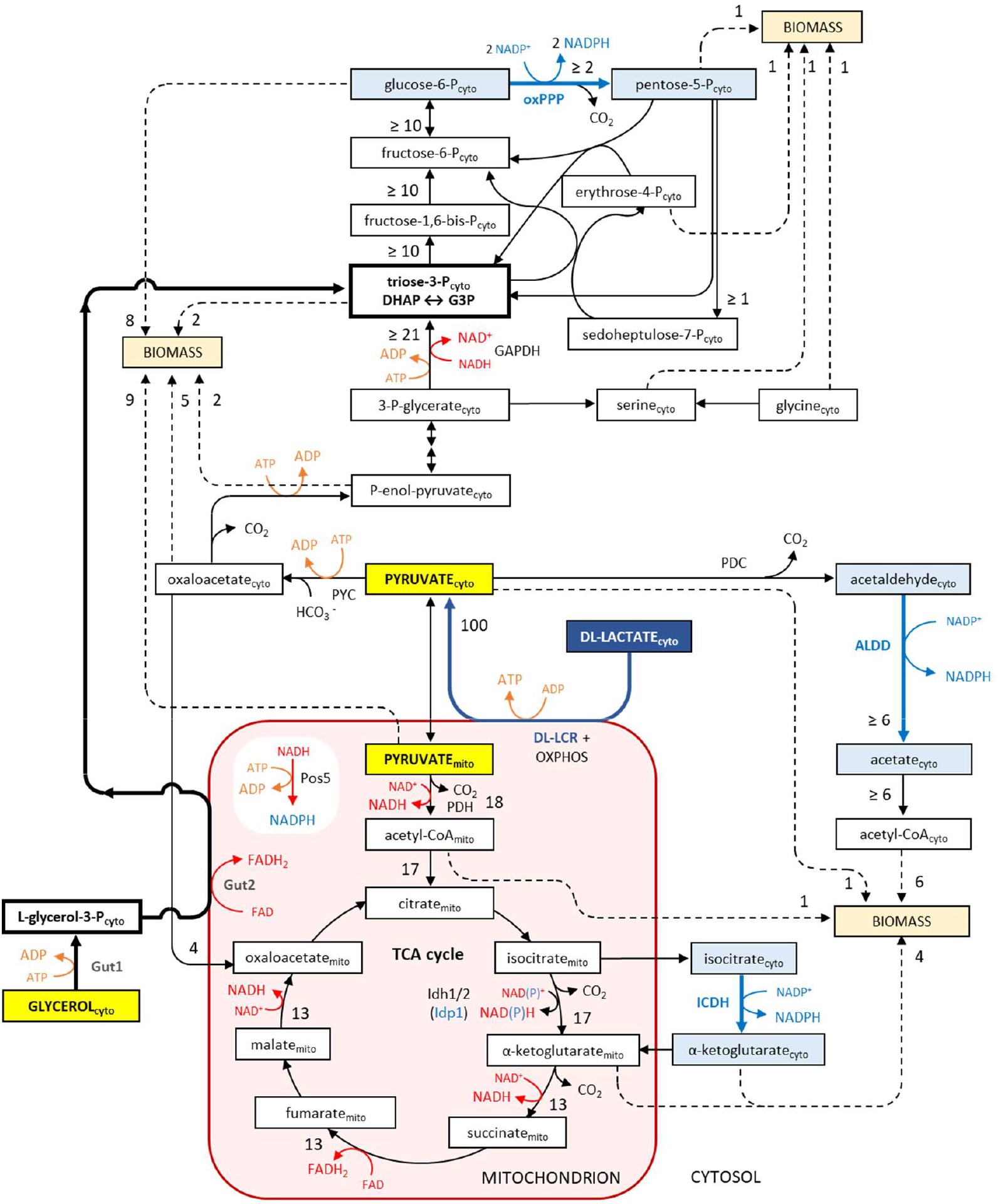
Postulated metabolic flux distribution of *S. cerevisiae* coutilizing lactate and glycerol. Adapted with modification from Figure 5I.2 in ^14^. cyto: cytosolic. mito: mitochondrial. PYC: pyruvate carboxylase. PDC: pyruvate decarboxylase. PDH: pyruvate dehydrogenase. DL-LCR: DL-Lactate:cytochrome c oxidoreductase. OXPHOS: oxidative phosphorylation system. Other abbreviations: see text. The light blue background and font in the cytosol respectively indicate metabolites and enzymes directly involved in cytosolic NADPH biosynthesis. The flux sizes are adapted for illustrative purposes from the published *in vivo* metabolic flux distribution of *S. cerevisiae* grown on pyruvate ^48^. Justification for the adoption: Note S4.

Cytosolic NADPH in *S. cerevisiae* can be generated through three routes (Figure 6): (*i*) the oxidative branch of the pentose phosphate pathway (**oxPPP**) catalyzed by Zwf1 and Gnd1/2 ^16,17^; (*ii*) the cytosolic aldehyde dehydrogenase (**ALDD**) catalyzed by Ald6/3/2 ^18,19^; and (*iii*) the cytosolic isocitrate dehydrogenase (**ICDH**) reaction catalyzed by Idp2 ^20^. Among the three routes, oxPPP is least likely to determine superior lactate growth, as the reaction is energetically expensive: it consumes one NADH and three ATP molecules for every cytosolic NADPH molecule generated from cytosolic pyruvate. ALDD is more energetically efficient as it merely involves decarboxylation, and ICDH is even more so, as it also generates a mitochondrial NADH molecule per reaction. Additionally, ICDH is superior to ALDD in that it does not require the formation of toxic acetaldehyde ^21^, and unlike acetate, the product (i.e., α-ketoglutarate) can directly participate in the TCA cycle without an activation step ^22^. In other words, the greater metabolic fate flexibility displayed by cytosolic α-ketoglutarate relative to cytosolic acetate imposes less (if any) penalty on the cell committing to the respective route.

Different *S. cerevisiae* strains may exhibit substantially distinct cytosolic NADPH production capacities. Figure 7A shows the median value and diversity index (defined as the ratio of standard deviation to median) of each cytosolic NADPH-producing enzyme when *S. cerevisiae* BY4742 was grown on 10 different carbon sources ^23^. The smaller the diversity index, the more similar the abundance across the carbon sources utilized; hence, the more likely it is to be constitutively expressed. In contrast, the higher the diversity index is, the more likely its expression is to be heavily regulated (read: induced or repressed) by carbon sources. As shown in Figure 7A, Zwf1, Gnd1, and Ald6 are the top three constitutive enzymes, while Ald3 is the most regulated one. Assuming that there is no interstrain diversity with regard to cytosolic NADPH production capacity, the abundance ratio of Ald6 to Zwf1 (i.e., the two constitutive enzymes) and Ald3 to Zwf1 (i.e., the heavily regulated one to the constitutive one) in cells grown on glucose should be conserved. In reality, the two ratios varied widely across various *S. cerevisiae* cultures ^24^: the fold difference between the highest and lowest values was 851 times for Ald6 (Figure 7B) and 468 times for Ald3 (Figure 7C). (See also Table S27).

**Figure 7.**
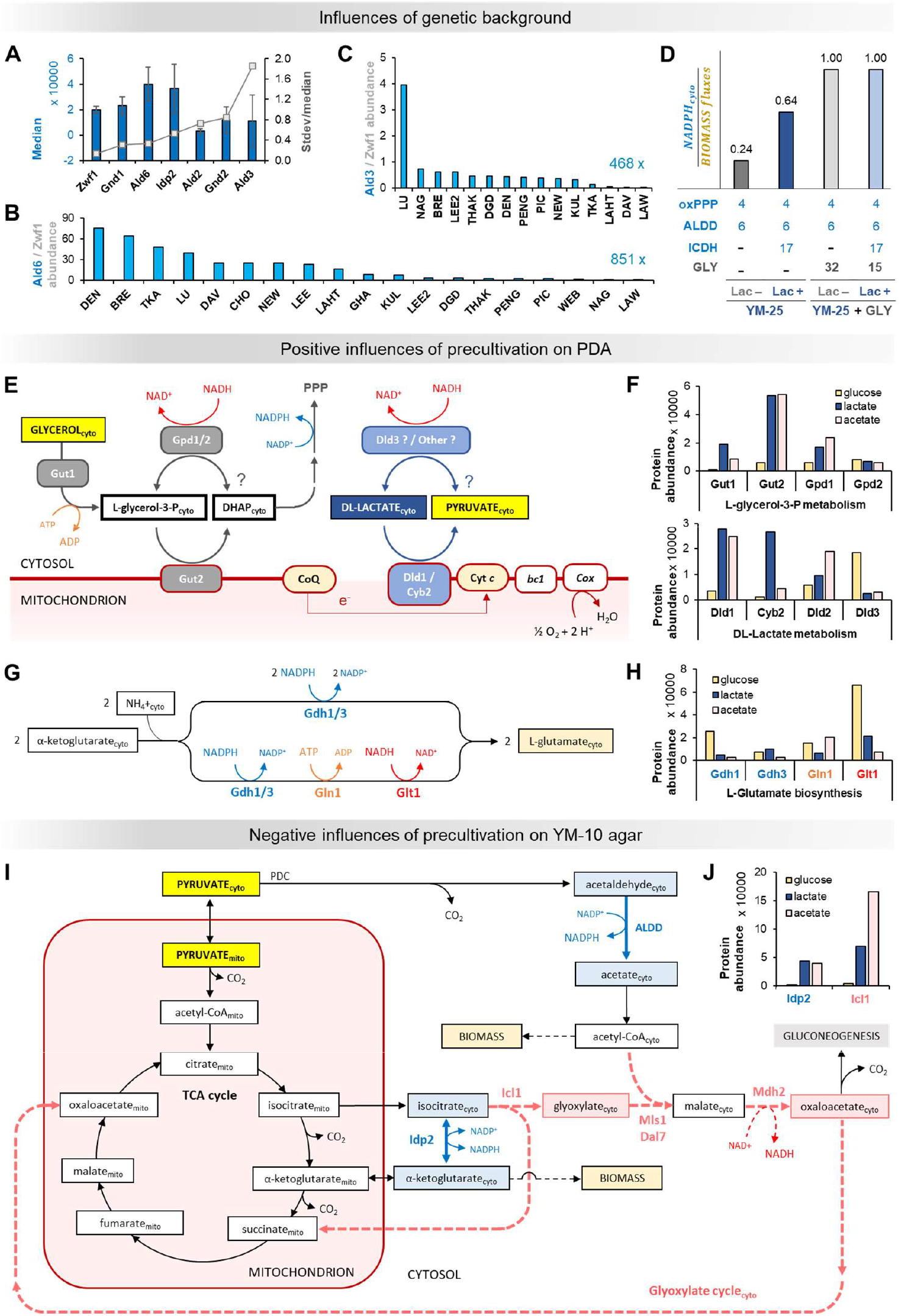
Postulated molecular mechanisms by which the genetic background and precultivation medium of *S. cerevisiae* can influence its growth on lactate—with and without glycerol supplementation. **A**, The abundance of proteins directly involved in cytosolic NADPH biosynthesis in *S. cerevisiae*. Source data: Table S27. **B, C**, The abundance of Ald6 (**B**) and Ald3 (**C**) relative to Zwf1 in various glucose-grown *S. cerevisiae* cultures. The label at the bottom right corner of each chart indicates the fold difference between the largest and the smallest ratios in the chart. Source data: Table S28. **D**, The estimated ratio of the total cytosolic NADPH to the total biomass synthesis fluxes of a lactate-grown *S. cerevisiae*, whose metabolic flux distribution is as shown in Figure 6, for each of the four hypothetical scenarios. The number below each bar: the amount of cytosolic NADPH contributed by the specified source. The size of the total biomass synthesis fluxes in all scenarios was 42 (Table S29). The number atop each bar indicates the ratio for the corresponding scenario. A ratio of 1.0 indicates theoretically uninterrupted exponential growth at the maximum specific growth rate. (See also Section 5.4.4.1 in ^14^ for clarity). GLY: glycerol utilization. Lac –: “Lactate –” strain. Lac +: “Lactate +” strain. Other abbreviations: see text. **E**, Reactions that may happen when *S. cerevisiae* is grown on a glycerol/lactate mixture. The “?” marks indicate reactions that have not been reported but are theoretically possible (see text for details). PPP: pentose phosphate pathway. CoQ: Coenzyme Q. Cyt *c*: Cytochrome *c. bc1*: Cytochrome *bc1* (complex III). *Cox*: cytochrome c oxidase (complex IV). **F**, The abundance of the eight proteins mentioned in (**E**) as a function of the carbon source used for growth. **G**, The two routes of cytosolic L-glutamate biosynthesis. Source data: Figure S17. **H**, The abundance of the four proteins directly involved in cytosolic L-glutamate biosynthesis as a function of the carbon source used for growth. **I**, The postulated interactions between a portion of the pyruvate utilization network and a portion of the glyoxylate cycle occurring in the cytosol. cyto, mito, light blue background and font: see Figure 6. The pink background and font respectively indicate metabolites and enzymes of the glyoxylate cycle that are relevant to the postulated scenario (see text for details). **J**, Idp2 and Icl1 abundances as a function of the carbon source used for growth. **F, H, J**, Source data: Table S31.

Different cytosolic NADPH production capacities should result in different cytosolic NADPH-to-biomass synthesis flux ratios. The ratio is believed to determine how long a strain can assume its maximum specific growth rate on lactate, and in turn, the total-growth rate (Section 5.4.4.1 in ^14^). To illustrate, using the postulated metabolic flux distribution of *S. cerevisiae* grown on lactate as a conceptual framework (Figure 6, Table S29), a complete absence of Idp2 activity will result in a flux ratio of 0.24, while the full activity of Idp2 will almost triple the flux ratio (Figure 7D, first and second bars). In fact, there is experimental evidence that a strain’s cytosolic NADPH production capacity can meaningfully affect its lactate growth phenotype. Boubekeur et al. ^13^ reported that *ALD4* and *PDA1* double deletion abolished the lactate growth capacity of strain YPH499 but not W303-1a, owing to its Ald6 activity being almost double that of the former.

In turn, a similar analysis can be conducted to predict the minimum amount of glycerol required to meet the cytosolic NADPH demand of different genetic backgrounds. The third and fourth bars of Figure 7D indicate that it is between 15-32 mol%, which is in close agreement with the most effective concentrations identified in this study (i.e., 10-49 mol%).

### Positive influences of precultivation on PDA

Compared to precultivation on YM-10 agar, precultivation on PDA substantially improved the growth yield of the “Lactate – (70)” strains in YM-25 + GLY by approximately 2.6 times (Figure 3C, right), making it similar to the growth yield attained by the “Lactate +” strains in the same media (i.e., approximately 70% of their growth yield in YM-21). Such a drastic improvement was not seen when the same “Lactate – (70)” strains were cultivated in YM-25 (Figure 3C, left), suggesting that the improvement originated not from lactate-but instead from glycerol-utilization.

Growth on PDA is essentially growth on glucose, whereas that on YM-10 agar is essentially growth on acetate. Consequently, it is reasonable to think that any improvement caused by precultivation on PDA on “Lactate – (70)” strains was mediated by proteins involved in glycerol utilization that are expressed differentially on glucose versus acetate. These proteins should determine how glycerol is metabolized by the first few generations, before the cell finishes reshaping its proteome to best accommodate its growth on lactate.

Glycerol utilization in *S. cerevisiae* is believed to be mediated by Gut1 and Gut2 ^25^ (Figure 6). At least in glucose-grown *S. cerevisiae*, the latter also collaborates with Gpd1 and Gpd2 to form the L-glycerol-3-P shuttle involved in cytosolic NADH regeneration ^26,27^ (Figure 7E). The abundance of Gut1, Gut2, and Gpd1 is much lower in cells grown on glucose than on acetate, with the most pronounced difference displayed by Gut2 (∼9 times less abundant), followed by Gut1 (∼7 times less abundant), and Gpd1 (∼4 times less abundant) ^23^. In contrast, the abundance of Gpd2 was ∼40% higher ^23^ (Figure 7F, top).

At a glance, the meager amounts of Gut1 and Gut2 provided by precultivation on PDA seem to be at odds with supposedly superior glycerol utilization. There are two possible explanations for this antithesis. First, it could be that their seemingly low abundance is actually the appropriate amount, as there is likely an inherent limit to the amount of glycerol-derived DHAP that can be processed by Zwf1 and Gnd1/2. From this perspective, any Gut1 excess may simply waste precious ATP through unrestrained glycerol phosphorylation, and any excess Gut2 may simply occupy the mitochondrial membrane area and/or titrate flavin otherwise available for other proteins, including those involved in lactate utilization and protein synthesis (*vide infra*). (Note that Gut2 is a flavoprotein ^28^.) Second, it could also be that the low amounts of Gut1 and Gut2 actually pave the way for the processing of some L-glycerol-3-P by Gpd1/2, thus effectively resulting in net cytosolic NADH generation. Cytosolic NADH is not copiously produced through the gluconeogenesis pathway in lactate-grown *S. cerevisiae* (Figure 6); however, its presence is required for the glyceraldehyde-3-P dehydrogenase (GAPDH) reaction. (Note that the cell cannot transport the mitochondrial NADH generated through the TCA cycle to the cytosol ^29^). One possible way lactate-grown cells produce cytosolic NADH is through the activity of NAD^+^-linked glutamate dehydrogenase (Gdh2) in a manner that essentially transforms cytosolic NADPH consumed by Gdh1/3 to cytosolic NADH (Figure 1 in ^30^). From the perspective of lactate-grown *S. cerevisiae* (that is cytosolic NADPH-limited), any mechanism that can generate cytosolic NADH without sacrificing any cytosolic NADPH, such as the Gpd1/2 reaction, will significantly improve biomass synthesis capacity (hence, growth yield). Similarly, the presence of cytosolic NADH also permits L-glutamate synthesis via the “NADPH-minimal” pathway, i.e., the one involving Gln1 and Glt1 on top of Gdh1/3 ^31–33^ (Figure 7G, bottom). Otherwise, the reaction should proceed exclusively through the first pathway, which solely involves Gdh1/3 and cytosolic NADPH (Figure 7G, top). Glt1 is present in a glucose-grown cell at a level that is ∼9 times higher than that in an acetate-grown cell ^23^—almost like the opposite of Gut2 (Figure 7H), thus providing a means for the “NADPH-minimal” L-glutamate synthesis pathway to occur should cytosolic NADH be available.

Given the potential of the second possible explanation (i.e., that the cell processes a substantial amount of L-glycerol-3-P not via Gut2 but instead via Gpd1/2 and couples it with the generation of cytosolic NADH), the question is whether Gpd1/2 can actually catalyze the postulated reverse reaction.

At pH 7.0, a Gpd isolated from *S. cerevisiae* H44-3D catalyzed the reverse reaction at 3% of the forward reaction rate ^34^. Similarly, at pH 7.2, the enzyme isolated from *S. cerevisiae* YSH 11-6B did so at 0.1% of the forward reaction rate ^35^. The specific identity of the enzyme in both cases was not explicitly stated; however, it was likely the Gpd1, as the molecular mass was 41,000 – 47,000 Da. (The theoretical molecular mass of Gpd2 is > 49,000 Da ^36^). Their study and others ^37,38^ indicated that its *K*_*m*_ for DHAP was 0.018 – 0.13 mM and for NADH 0.37 – 1.6 mM. The forward reaction was sensitive to acidic pH—no activity could be detected at pH 5.0—and high ionic strength ^38^. Meanwhile, the Gpd2 in the *gpd1Δ* W303-1A cell extract exhibited *K*_*m*_ for DHAP of 0.86 mM and NADH of 0.018 mM ^39^. Neither Gpd1 nor Gpd2 has been thoroughly characterized for their reverse reaction capacities. However, NAD^+^ strongly competes with NADH for the active site ^35^.

Based on the above-mentioned facts, there is a reasonable chance for Gpd1/2 to catalyze the proposed reverse reaction *in vivo*, especially when the cell is grown on a glycerol/lactate mixture dominated by lactate. Inside the cell, lactate may reduce the cytosolic pH to below 5.5 ^40^ and induce the entry of various countercations, effectively raising the ionic strength of the cytosol and thus inhibiting the forward-, but perhaps not the reverse-, reaction. Additionally, since the concentrations of L-glycerol-3-P and NAD^+^ at all times are expectedly higher than those of DHAP and cytosolic NADH (e.g., due to the reverse GAPDH reaction and rapid protein synthesis via the “NADPH-minimal” pathway, *vide supra*), there is less chance for the enzyme to catalyze the forward reaction. Under such conditions, the reverse reaction rate may be higher than a mere 3% of that of the forward reaction; alternatively, even a relatively slow rate may be enough to merely provide supplemental cytosolic NADPH according to the postulated scheme (Figure 6). (Note that it is lactate that generates nearly all biomass precursors and energy molecules.) The enzyme catalyzing the reverse reaction may be Gpd2, whose *K*_*m*_ for DHAP is slightly higher than that of Gpd1.

Relative to precultivation on YM-10 agar, precultivation on PDA also resulted in an approximately 60% higher growth yield of “Lactate +” strains cultivated in YM-25 (Figures 1C, 3C left). However, the improvement was no longer observed when the strains were cultivated in YM-25 + GLY (Figure 3C, right), mainly because the strains precultivated on YM-10 agar now managed to reach a comparable growth yield through glycerol coutilization. These observations indicated that in the “Lactate +” case, any growth advantage imparted by precultivation on PDA was mediated by proteins involved in lactate utilization that are differentially expressed on glucose versus acetate.

There are at least four flavoproteins involved in lactate utilization: Dld1, Cyb2, Dld2, and Dld3 ^28^ (Table S30). The abundance of the first three enzymes, which are mitochondrial, was substantially (∼3-7 times) lower in cells grown on glucose than on acetate, whereas the abundance of Dld3, which is cytosolic, was ∼6 times higher ^23^ (Figure 7F, bottom). Among the four, Dld3 is the only one capable of generating cytosolic NADH, i.e., through its hypothetical reverse reaction (Figure 7E).

Among the three Dld enzymes, Dld3 exhibits the lowest *K*_*m*_ for D-lactate (533 μM), which is almost an order of magnitude lower than the other two enzymes (i.e., 4450 - 5215 μM) ^41^. Combined with its high abundance on glucose, it is possible that Dld3 catalyzes the reverse reaction in glucose-grown “Lactate +” cells and in turn imparts growth advantage in YM-25. The extent of the advantage, however, is substantially less than what was seen for “Lactate – (70)” in YM-25 + GLY (i.e., ∼40% vs. ∼150%) (Figure 3C), perhaps because the main limitation of lactate growth is cytosolic NADPH (not NADH).

### Negative influences of precultivation on YM-10 agar

Relative to precultivation on PDA, precultivation of *S. cerevisiae* on YM-10 agar tended to result in a lower growth yield. The effect was most pronounced when the “Lactate – (70)” strains were cultivated in YM-25 + GLY, followed by the “Lactate +” strains cultivated in YM-25 (Figure 3C).

To rationalize the trend, it is useful to notice that acetate utilization in *S. cerevisiae* intersects with lactate utilization at the cytosolic isocitrate node (Figures 7I, S18), among others. In addition to being the substrate of Idp2 to generate cytosolic NADPH, cytosolic isocitrate in acetate-grown *S. cerevisiae* is also used by Icl1 to generate cytosolic glyoxylate, the key metabolite of the glyoxylate cycle.

The *K*_*m*_ value of Icl1 for isocitrate is around 1.2–2.4 mM ^42–44^, while that of Idp2 is around 0.04-0.22 mM ^43,45^ (Table S32). Given the same amount of isocitrate and enzymes, the forward flux of Idp2 should dominate. However, acetate-grown *S. cerevisiae* contains ∼4 times more Icl1 than Idp2 ^23^ (Figure 7J). Additionally, two cytosolic intermediates of the glyoxylate cycle, glyoxylate and oxaloacetate, as well as their putative condensation product oxalomalate, can inhibit the forward reaction of Idp2 by 5-50%, depending on the concentration ^43^. Idp2 also exhibits *K*_*m*_ values for α-ketoglutarate and NADPH that are very similar to those for D-isocitrate and NADP^+^, indicating its equal capability in catalyzing the reverse reaction (i.e., α-ketoglutarate reduction) ^45^ Based on the information, it is possible that in acetate-grown *S. cerevisiae* (*i*) any cytosolic isocitrate will predominantly be processed by Icl1 (instead of Idp2), and (*ii*) a substantial amount of cytosolic α-ketoglutarate released by the mitochondria will not be used for biomass synthesis, but instead to generate more cytosolic isocitrate to feed the glyoxylate cycle. The cytosolic NADPH fueling the reverse Idp2 reaction is likely to be synthesized by ALDD, effectively reducing the amount available for actual biomass synthesis. Altogether, these possibilities may explain the typically lower growth yield displayed by the YM-10 agar-precultivated cells.

Icl1 produced during the precultivation step should theoretically diminish over time due to cell division. However, *S. cerevisiae* grown on lactate reportedly contained Icl1 at a level comparable to that of Idp2 ^23^ (Figure 7J), indicating deliberate Icl1 expression by the cell. In fact, the same lactate-grown *S. cerevisiae* contained a substantial amount of the four enzymes involved in the cytosolic isocitrate-to-oxaloacetate conversion (i.e., Icl1, Mls1, Dal7, and Mdh2) at a level that was 14-27 times higher than that in glucose-grown cells ^23^ (Figure S18, Table S33). In contrast, the other four glyoxylate cycle enzymes (Mdh3, Cit2, Aco1, and Aco2) were only expressed at 0.8–5.8 times their level in glucose-grown cells ^23^. A possible explanation for this phenomenon is that when grown on lactate, *S. cerevisiae* needs to deliberately synthesize cytosolic oxaloacetate, either from cytosolic isocitrate or α-ketoglutarate, to feed the gluconeogenic pathway in a manner that also generates the much-needed cytosolic NADH (*vide supra*) ^46^. Note that the pyruvate carboxylation (PYC) route consumes ATP (Figure 6), while direct export of mitochondrial oxaloacetate to the cytosol does not generate any cytosolic NADH (besides necessitating pyruvate carboxylation or any alternative reactions to occur for anaplerosis purposes).

### Inconsistent ability of *S. cerevisiae* to grow solely on glycerol

Despite representing a wide range of genetic backgrounds and being precultivated in a manner that minimizes carbon catabolite repression, none of the 22 *S. cerevisiae* strains studied here demonstrated a “dose-dependent” growth profile on glycerol, in which the amount of biomass generated corresponded to the initial amount of limiting carbon source. The sporadic growth profile was in contrast to that of the two non-*S. cerevisiae* strains studied here (Figures 2D, S11) and even an *S. cerevisiae* strain cultivated at various lactate concentrations ^47^ (Figure S19). These observations confirm that by design, *S. cerevisiae* cannot utilize glycerol as the sole carbon and energy source until specific mutations eventually permit glycolysis and gluconeogenesis to co-occur (mechanism #1) and/or permit some endogenous processes, such as protein degradation, to provide various central metabolism intermediates at the required rates (mechanism #2). (See also Section 5.1.2 in ^14^). In turn, the inconsistency invalidates the apparent growth on glycerol observed in this study, as it must have been conducted by spontaneously emerged mutants as opposed to the wild-type strains initially inoculated. In other words, the apparent contribution of glycerol carbons to biomass must be ignored during the interaction analysis (Figure 4A). The possibility that the cells underwent the same mutation(s) when cultivated in a glycerol/lactate mixture is likely negligible. As shown in the growth profiles of PDA-precultivated “Lactate +” and “Lactate – (70)” strains at day 2.6 (Figure S9, for example, Y1136 and Y1613 compared with the two non-*S. cerevisiae* Y1125 and Y1728), there was still no appreciable growth in the YM-GLY media, even though substantial growth was already observed in the YM-25 + GLY media. This observation suggests that the presence of lactate (i.e., the genuine carbon and energy source) could dampen any directional mutation pressure toward glycerol utilization as the sole carbon and energy source.

### Beyond synergism: the saturation and antagonism in glycerol/lactate coutilization

Under certain conditions, especially at high glycerol loading, glycerol/lactate coutilization resulted in saturated synergism and antagonism (Figure 4C). Since the amount of lactate in this study was kept constant across the different glycerol/lactate mixtures, the causes must lie in the glycerol utilization pathway. The saturation effect may be caused by the maximum processing capacity of the oxPPP (Figure 7E). In such a case, regardless of the amount of glycerol entering the cell, once the oxPPP maximum capacity has been reached, no excess glycerol can result in more cytosolic NADPH. The antagonistic effect may be caused by ATP wasting due to excessive Gut1 activity. (Note that *S. cerevisiae* grown on lactate tends to produce a substantial amount of Gut1 ^23^, Figure 7F, top). Another possibility is competition between Gut2 and Dld1/Cyb2 for the mitochondrial membrane and/or Cytochrome *C* (Figure 7E). Excess intracellular glycerol may be processed by Gut2, which eventually transfers the electrons to Coenzyme Q, which then transfers them to Cytochrome *C*. On the other hand, *S. cerevisiae* grown on lactate gradually relies on mitochondrial membrane-bound Dld1/Cyb2 (instead of cytosolic Dld3) for processing lactate (Figure 7F). It may be difficult for Dld1/Cyb2 under excess glycerol to assemble onto the mitochondrial inner membrane and/or to interrupt the quasi-continuous electron flow from glycerol ^27^, effectively declining lactate utilization and hence the synthesis of biomass precursors and ATP. Note how the cell physiological state can modulate the outcome: given the same glycerol/lactate molar ratio, cells with a lower amount of Gut1 and/or Gut2 will have a lower chance of suffering from such an electron transport chain crisis. This may explain the diversity in the *S. cerevisiae* responses shown in Figure 4C.

Aside, it is tempting to surmise that the growth performance of CEN.PK113-7D reported in ^14^ (Figure S2) could have been better had the cell been precultivated not in liquid YM-10 but instead on PDA, and exposed not to a 3.08 glycerol/lactate molar ratio but instead a 0.1 or 0.5 one.

## Concluding remarks

This study aimed to investigate whether the glycerol cofeeding strategy, once improved lactate utilization in *S. cerevisiae* CEN.PK113-7D, could also benefit other yeast strains, preferably at a lower glycerol/lactate molar ratio. The growth profiles of 22 natural *S. cerevisiae* and one nonconventional strain (i.e., *W. versatilis*) isolates suggested that it was possible regardless of the strain’s basal lactate utilization capacity. Specifically for *S. cerevisiae*, the three highest growth yield improvements, reaching between 86% to 94% of the maximum theoretical yield relative to glucose, were achieved at 0.1 and ∼0.5 molar ratios. The empirically observed most effective molar ratios agree with the underlying theoretical framework. Importantly, the study reveals how precultivation on PDA (i.e., on glucose) tends to influence the growth yield positively, while precultivation on YM-10 agar (i.e., on acetate) influences it negatively. Additionally, a high glycerol loading may result in no further improvement and occasionally even reduce the growth yield. The existing knowledge of *S. cerevisiae* metabolism allows the postulation of molecular mechanisms behind the observed phenomena. In turn, the molecules and metabolic pathways putatively involved can serve as the targets for future metabolic engineering endeavors.

## Materials and methods

### Yeast strains

All strains used in this study were obtained from the Indonesian Culture Collection (InaCC) (inacc.brin.go.id). All except one (Y93) were isolated from various types of sources (such as traditional food or beverages) from different geographical regions in Indonesia. The strain pool was deliberately selected to cover a wide range of genetic makeup diversity. Each isolate was provided as a fresh culture on PDA (Oxoid CM0139) in a 100 x 15 mm disposable petri dish format. The species identity of each strain was confirmed by sequencing the 28S ribosomal RNA of the cytoplasmic ribosome large subunit (LSU), followed by the Nucleotide BLAST analysis (https://blast.ncbi.nlm.nih.gov/Blast.cgi). The sequencing revealed the actual identities of Y1125 and Y1728 as *Wickerhamiella versatilis* and *Clavispora lusitaniae*, respectively.

### Preparation of PDA and YM-10 agar plates

PDA was prepared from a commercial reagent (Himedia M096) according to the manufacturer’s instructions. Specifically, the sterile agar solution was prepared by dissolving 39 g of the commercial powder into 1 L of distilled water, followed by its autoclavation (along with a magnetic bar inside) at 121°C for 15 minutes. The magnetic bar-bearing autoclaved solution was cooled to 60°C by placing it in a 60 °C incubator until the pouring time.

The sterile YM-10 agar solution was prepared from two sterile solutions: (*i*) an autoclaved 4.4% agar solution and (*ii*) a filter-sterilized 2X YM-10 (pH 5.8) solution. The final pH value of the 2X solution was chosen to ensure that the resulting pH of the 1X YM-10 agar solution was 6.0.

The 4.4% agar solution was prepared in a final volume of 500 mL in a 1-L glass bottle and autoclaved together with a magnetic stir bar at 121°C for 15 min. The autoclaved agar solution (containing a magnetic stir bar) was placed in a 60°C incubator until the pouring time.

To prepare the 2X YM-10 solution, 20 g of glacial acetic acid (Table M1) was mixed with 400 ml of distilled water. KOH granules were added to the solution under continuous magnetic stirring until the pH reached a value of 5.0. The following compounds were then added in the order they were listed: 1.95 g of Na_2_HPO_4_ (Mw 141.96), 11.91 g of NaH_2_PO4.H_2_O (Mw 137.99), 2.27 g of urea, and 1.70 g of YNB–AA–AS (Table M1). Several drops of 2 M KOH solution and some volume of distilled water were added to the solution under constant magnetic stirring and pH monitoring until the medium reached a final volume of 500 mL and a final pH of 5.8. The final solution was incubated in a 60°C incubator until the temperature reached approximately 60°C.

**Table M1.** Description of key reagents used in this study.

| Reagent and molecular weight (Mw), if applicable | Source | Identifier |
| --- | --- | --- |
| PDA | Oxoid | CM0139 |
| PDA | Himedia | M096 |
| Agar | Himedia | PCT0901 |
| Glacial acetic acid | Sigma-Aldrich | 1.00063 |
| D-(+)-Glucose anhydrous | Himedia | PCT0603 |
| Lactic acid solution | Sigma-Aldrich | 252476 |
| Glycerol | Sigma-Aldrich | G7757 |
| Yeast nitrogen base without amino acids and ammonium sulfate (YNB – AA – AS, NutriSelect® Basic) | Sigma-Aldrich | Y1251 |
| Urea (Mw 60.06) | Sigma-Aldrich | 1.08487 |
| NaH<sub>2</sub>PO<sub>4</sub>.H<sub>2</sub>O (Mw 137.99) | Sigma-Aldrich | 1.06346 |
| Na<sub>2</sub>HPO<sub>4</sub> (Mw 141.96) | Sigma-Aldrich | S0876 |
| KOH (Mw 56.11) | Sigma-Aldrich | 1.05012 |

To obtain sterile YM-10 agar solution, 2X YM-10 solution at approximately 60°C was filtered through a disposable bottle-top vacuum filter lined with a 0.22 μm sterile polyethersulfone membrane (LabSelect, 42311) that was attached to a bottle containing autoclaved 4.4% agar solution.

To prepare the agar plates, the 60°C sterile medium agar solution was first subjected to careful magnetic stirring, which ensured thorough mixing without air bubbles formation. The homogenous solution was then poured into sterile 100 mm × 15 mm polystyrene Petri dishes inside a laminar airflow bench at approximately 20 mL per dish. The plates were kept at room temperature to dry for approximately 3 days, after which they were stored inside a plastic bag in a 4°C refrigerator until use.

### Preparation of liquid media for experimental cultivation

The liquid media for the growth profiling experiment was prepared from the stock solutions described in Table M2 using reagents listed in Table M1. Those stock solutions were freshly prepared at the start of experimental media preparation and not filter-sterilized.

**Table M2.** Stock solutions for the preparation of experimental liquid media.

| Stock solution | Preparation |
| --- | --- |
| Glycerol, 50% w/w | 25 g glycerol +<br>25 g of distilled water |
| 10x Lactic acid pH 5.0 (KOH) | 1.18 g of Lactic acid (85% purity) +<br>KOH granules until the pH was 5.0 +<br>distilled water until the volume was 50 mL |
| 10x Na-phosphate buffer | 11.91 g of NaH<sub>2</sub>PO<sub>4</sub>·H<sub>2</sub>O (Mw 137.99) +<br>1.95 g of Na<sub>2</sub>HPO<sub>4</sub> (Mw 141.96) +<br>distilled water until the volume was 100 mL |
| 10x Urea | 2.27 g urea +<br>distilled water until the volume was 100 mL |
| 10x YNB – AA – AS | 1.7 g YNB – AA – AS +<br>distilled water until the volume was 100 mL |
| 2 M KOH | 11.2 g KOH +<br>distilled water until the volume was 100 mL |

To prepare the liquid media for experimental cultivation, the desired amount of carbon source(s), i.e., glucose, lactic acid, and/or glycerol as indicated in Table M3, was added into a 50 mL conical centrifugal tube, followed by the following in the order they were mentioned: a 1 cm magnetic bar, 5 mL of 10x Na-phosphate buffer, 5 mL of 10x Urea, 5 mL of 10x YNB – AA – AS, and distilled water until the volume was approximately 40 mL. Under constant magnetic stirring and pH monitoring, several drops of 2 M KOH were added until the solution’s pH was 6.0. Distilled water was then added until the final volume was 50 mL.

**Table M3.**
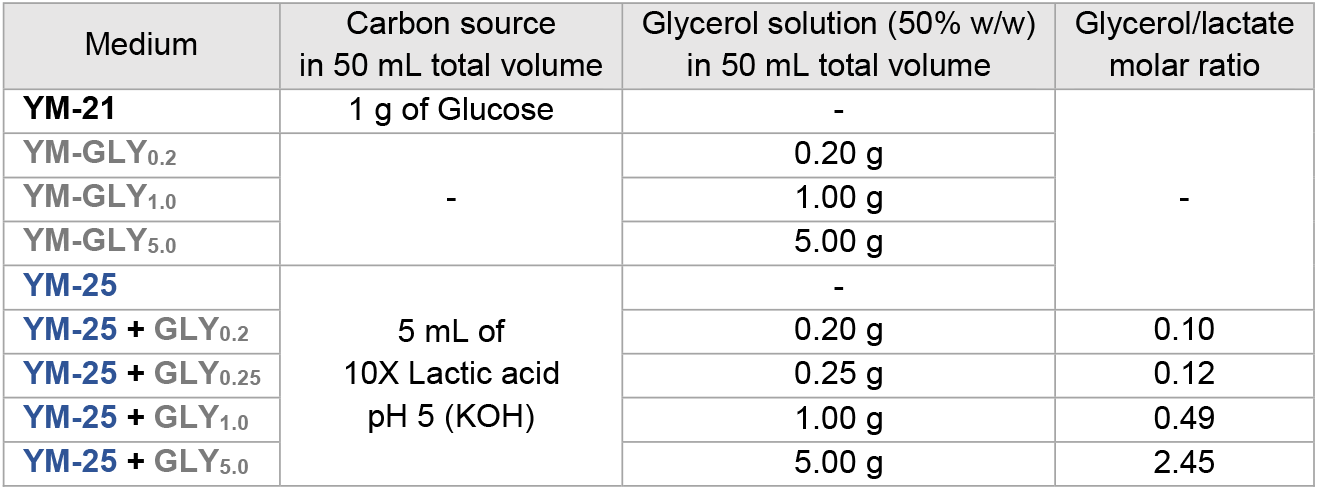
Carbon source composition of each liquid medium for experimental cultivation.

Each medium was filter-sterilized into a sterile 50 mL conical tube using a 50 mL disposable syringe and a sterile 0.45 μm nylon filter inside a laminar airflow bench. The filter-sterilized media were immediately used or sealed with a piece of parafilm and stored at room temperature until use.

### Precultivation on PDA and YM-10 agar plates

The cryostock of the intended strain was scrapped with a sterile disposable inoculating loop and subsequently quadrant-streaked onto a relevant agar plate. The plate was incubated at 30°C for several days until single colonies were present. The plate was then sealed with a piece of parafilm and stored in 4°C fridge until use.

### Experimental cultivation in 24-deep well plates

Two-and-a-half milliliters of each of the experimental media were pipetted into the intended well in the 24-deep well plate screening plates (Axygen, P-DW-10ML-24-C) according to the map in Figure S4. To create the inoculum, three single colonies on the PDA plate were individually touched with a sterile pipette tip and resuspended in 900 μl filter-sterilized distilled water. The inoculum was mixed through repeated pipetting and distributed into each well on the screening plate at as much as 25 μL per well. The completely inoculated screening plate was covered with a sterile breathable membrane (Axygen, BF-400-S) and placed in a shaker incubator (Taitec, BioShaker BR-43FM) set at 30 °C and 250 rpm. At the end of cultivation, 100 μL of individual cultures were placed in the designated well in a clear 96-well microtiter plate according to the map in Figures S5-S7. The absorbance at 600 nm was read by a microtiter plate reader (Tecan, Infinite 200Pro). The OD_600_ of individual cultures was determined through subtraction of OD_600_ of distilled water located in the same well.

### Analysis of residual lactate concentration in spent media via high-performance liquid chromatography (HPLC)

At the end of cultivation, one mL of culture of interest was filtered through a 0.22 μm hydrophilic PTFE syringe filter (Labfil C0000608) into an HPLC vial. The HPLC analyses were conducted on a Shimadzu liquid chromatograph system (Shimadzu Corp, Kyoto, Japan) equipped with a Coregel 87H column and a diode array detector (DAD). The column was maintained at 80°C. The mobile phase was 5 mM H_2_SO_4_ flown at the rate of 0.6 mL per minute. The DAD detector was applied to acquire spectral information within a range of 190 to 400 nm. (The optimum wavelength for lactate was 211 nm). Results were acquired and processed by the Shimadzu Labstation software (Shimadzu Corp).

## Acknowledgements

This study was supported by the Research Organization for Life Sciences and Environment of BRIN (grant no. 50/III.5/HK/2024). W.N. was supported by the Human Resource Capacity Development Program of BRIN (grant no. 166/HK/II/2024). The authors thank Radityo Pangestu, PhD (Research Center for Genetic Engineering of BRIN) for suggestions regarding quantitative HPLC strategies. The authors acknowledge the facilities, scientific, and technical support from the Advanced Characterization Laboratories Cibinong – Integrated Laboratory of Bioproducts (iLaB), the Indonesian Culture Collection (InaCC), and the Genomic Laboratories of BRIN through the E-Layanan Sains (ELSA) system.

## Author contributions

W.N. reviewed the literature, proposed the central hypotheses, designed and performed the experiments, analyzed the data, presented the experimental results, and wrote the manuscript with input from all authors. A. K. reviewed the yeast growth data. O.F. reviewed the mathematical analysis and contributed additional funding for data acquisition. W.F. contributed additional funding for data analysis, supervised the research, and administered the grants. All authors have read, edited, and approved the manuscript.

