## Supplemental Information for "Synergistic Glycerol Coutilization Promotes Metabolic Fitness of *Saccharomyces cerevisiae* on Lactate"

#### List of supplementary notes

### Note S1. Potential abundance of wet waste-derived lactic acid.

To obtain a realistic estimate of the abundance of lactic acid (LA) that can potentially be derived from wet waste,

- the recent and comprehensive 2023 billion-ton report assessing the amount of renewable carbon sources in the U.S. [1] will be used as the data source for illustrating the wet waste abundance;
- the performance of a semi-continuous, pilot-scale MAAD reported by Wu et al. [2] will be used as the data source for wet waste-to-lactic acid conversion metrics; and
- the review by Skaggs et al. [3] will be used as the data source for the range of organic carbon content of wet waste.

To put the potential lactic acid abundance into perspective, it will be compared to the amount of sugar produced in the U.S. on a Cmol basis.

##### Table S1. Parameters used to calculate the potential abundance of wet waste-derived lactic acid in the U.S.

"Pubchem" refers to https://pubchem.ncbi.nlm.nih.gov/.

| Object | # | Parameter | Unit | Magnitude | Min. | Max. | Reference |
|---|---|---|---|---|---|---|---|
| Wet waste in the U.S. | 1 | Total fats, oils, grease (FOG) and other wet waste in the U.S. | $\left[\frac{dry\ short\ ton\ wet\ waste}{year}\right]$ | - | 3 + 32 = 35 million | 4 + 43 = 47 million | Table ES-2 in [1] Min.: near-term Max.: mature-market high |
| Wastewater feedstock for MAAD | 2 | Substrate (COD) concentration | $\left[\frac{g\ COD}{L\ wastewater}\right]$ | 80.0 ± 6.2 | 73.80 | 86.20 | Table A1 in the supplement of [2] |
| | 3 | Total organic carbon | $\left[\frac{g\ carbon_{wastewater}}{L\ wastewater}\right]$ | 28.0 ± 3.7 | 24.30 | 31.70 | |
| MAAD product stream | 4 | Total carboxylic acid (TCA) concentration | $\left[\frac{g\ TCA}{L\ MAAD\ product\ stream}\right]$ | 40.6 ± 1.1 | 39.50 | 41.70 | Table 1 in [2] |
| | 5 | Lactic acid (LA) | $\left[\frac{LA}{TCA}wt.\%\right]$ | 44.55 ± 6.88 | 37.67 | 51.43 | |
| | 6 | TCA yield | $\left[\frac{g\ COD_{TCA\ produced}}{g\ COD_{fed}}\right]$ | 0.76 ± 0.01 | 0.75 | 0.77 | |
| | 7 | LA yield | $\left[\frac{g\ COD_{LA\ produced}}{g\ COD_{fed}}\right]$ | 0.27 ± 0.01 | 0.26 | 0.28 | |
| General | 8 | LA molecular weight | $\left[\frac{g\ LA}{mol\ LA}\right]$ | 90.08 | - | - | Pubchem |
| | 9 | LA carbon content | $\left[\frac{mol\ carbon_{LA}}{mol\ LA}\right]$ | 3 | - | - | Pubchem |
| | 10 | Carbon atomic weight | $\left[\frac{g\ carbon}{mol\ carbon}\right]$ | 12.011 | - | - | Pubchem |
| | 11 | FOG and other wet waste carbon content | $\left[\frac{carbon_{wet\ waste}}{dry\ wet\ waste}wt.\%\right]$ | - | 38.8 | 76.3 | Table 2 in [3] Min.: dairy cows Max.: FOG (rendered animal fat) |
| | 12 | - | $\left[\frac{g}{dry\ short\ ton}\right]$ | 0.907 million | - | - | www.convertunits.com |
| | 13 | - | $\left[\frac{Cmol\ LA}{g\ LA}\right]$ | 0.033 | | | Pubchem |
| | 14 | - | $\left[\frac{Cmol\ sugar}{g\ sugar}\right]$ | 0.035 | | | Pubchem Sugar = sucrose ($C_{12}H_{22}O_{11}$, Mw 342.30) |

**Table S2. Intermediary- and final- calculation results.**

| # | Parameters operated | Unit | Magnitude | Min. | Max. | Reference |
|---|---|---|---|---|---|---|
| 15 | #5 x #4 | $\left[\frac{g\ LA_{produced}}{L\ MAAD\ product\ stream}\right]$ | 18.17 ± 4.65 | 14.88 | 21.45 | Equation S1 |
| 16 | #3 / #2 | $\left[\frac{g\ carbon_{wastewater}}{g\ COD}\right]$ | - | 0.33 | 0.37 | Equation S2 |
| 17 | #10 x #9 / #8 | $\left[\frac{g\ carbon_{LA}}{g\ LA}\right]$ | 0.40 | - | - | Equation S3 |
| 18 | #7 x #2 x #16 x #8 / #9 / #10 | $\left[\frac{g\ LA_{produced}}{L\ wastewater}\right]$ | 18.28 ± 3.46 | 15.83 | 20.73 | Equation S4 |
| 19 | #18 / #3 | $\left[\frac{g\ LA_{produced}}{g\ carbon_{wastewater}}\right] = \left[\frac{g\ LA_{produced}}{g\ carbon_{wet\ waste}}\right]$ | 0.65 | - | - | Equation S5 |
| 20 | #19 x #1 x #11 x #12 x #13 | $\left[\frac{Cmol\ LA_{produced}}{year}\right]_{the\ U.S.}$ | - | 0.26 x $10^{12}$ | 0.70 x $10^{12}$ | Equation S6 |
| 21 | - | $\left[\frac{Cmol\ sugar}{year}\right]_{the\ U.S.}$ | 0.29 x $10^{12}$ | - | - | Equation S7 |
| 22 | #20 / #21 | $\left[\frac{Cmol\ LA_{produced}}{Cmol\ sugar}\right]_{the\ U.S.}$ | - | 0.90 | 2.41 | - |

#### Pre-calculation: making sure the physical meaning of parameters in [2] is understood

Since various parameters listed in Table 2 in [2] were obtained from a semi-continuous process, the "volume" of the product stream may not be the same as the "volume" of feed. As such, in the following, we will first check if the amount of LA stated as wt% of TCA in the *MAAD product stream* (parameter #5) matches the amount calculated using the LA-specific yield information (parameter #7, which links the LA in the MAAD product stream to the organic carbon in the *wastewater feed*).

Approach 1. Amount of LA produced, present in the MAAD product stream, stated as wt% of TCA: #5 x #4

$$\left[\frac{\boldsymbol{g\ LA_{produced}}}{\boldsymbol{L\ MAAD\ product\ stream}}\right] = \left[\frac{LA}{TCA} wt.\%\right] \times \left[\frac{g\ TCA}{L\ MAAD\ product\ stream}\right] \quad \text{Equation S1}$$

$$= 37.67\% \times 39.50 = \mathbf{14.88} \quad \text{Min.}$$

$$= 51.43\% \times 41.70 = \mathbf{21.45} \quad \text{Max.}$$

Approach 2. Amount of LA produced from the total organic carbon in wastewater

The relationship between COD concentration and total organic carbon: #3/#2

$$\left[\frac{\boldsymbol{g\ carbon}}{\boldsymbol{g\ COD}}\right] = \frac{\left[\frac{g\ carbon}{L\ wastewater}\right]}{\left[\frac{g\ COD}{L\ wastewater}\right]} \quad \text{Equation S2}$$

$$= \frac{24.30}{73.80} = \mathbf{0.33} \quad \text{Min.}$$

$$= \frac{31.70}{86.20} = \mathbf{0.37} \quad \text{Max.}$$

The relationship between the mass of (organic) carbon and the mass of LA: #10 x #9 / #8

$$\left[\frac{\boldsymbol{g\ LA\ carbon}}{\boldsymbol{g\ LA}}\right] = \frac{\left[\frac{g\ carbon}{mol\ carbon}\right] \times \left[\frac{mol\ LA\ carbon}{mol\ LA}\right]}{\left[\frac{g\ LA}{mol\ LA}\right]} = \frac{12 \times 3}{90.08} = \mathbf{0.40} \quad \text{Equation S3}$$

The amount of LA produced $[g]$ per liter of wastewater: #7 x #2 x #16 x #8 / #9 / #10, assuming that COD of wastewater is identical to COD of LA.

$$\left[\frac{\boldsymbol{g\ LA_{produced}}}{\boldsymbol{L\ wastewater}}\right] = \frac{\left[\frac{\cancel{g\ COD_{LA\ produced}}}{\cancel{g\ COD_{fed}}}\right] \times \left[\frac{\cancel{g\ COD}}{L\ wastewater}\right] \times \left[\frac{\cancel{g\ carbon_{wastewater}}}{\cancel{g\ COD}}\right] \times \left[\frac{g\ LA}{\cancel{mol\ LA}}\right]}{\left[\frac{\cancel{mol\ carbon_{LA}}}{\cancel{mol\ LA}}\right] \times \left[\frac{\cancel{g\ carbon}}{\cancel{mol\ carbon}}\right]} \quad \text{Equation S4}$$

$$= \frac{0.26 \times 73.80 \times 0.33 \times 90.08}{3 \times 12.011} = \mathbf{15.83} \quad \text{Min.}$$

$$= \frac{0.28 \times 86.20 \times 0.37 \times 90.08}{3 \times 12.011} = \mathbf{20.73} \quad \text{Max.}$$

By comparing Approach 1 (Equation S1) and Approach 2 (Equation S4) (i.e., parameters #15 and #18 in Table S2), it can be seen that the two pre-calculation approaches generate pretty similar results with very minor differences. As such, it can be safely concluded that (*i*) the "volume"of the product stream is the same as the "volume"of feed, (*ii*) that the COD of wastewater is the same as the COD of LA (which is the basis of Equation S4), and (*iii*) the physical meaning of the parameters given is understood.

#### Calculating the potential abundance of wet waste-derived lactic acid produced per year in the U.S.

Amount of LA produced $[g]$ per amount of carbon in wastewater $[g]$: #18 / #3. The amount is assumed to be identical with the amount of LA produced per amount of carbon in any type of wet waste.

$$\left[\frac{\boldsymbol{g\ LA_{produced}}}{\boldsymbol{g\ carbon_{wastewater}}}\right] = \left[\frac{g\ LA_{produced}}{g\ carbon_{wet\ waste}}\right] = \frac{\left[\frac{g\ LA_{produced}}{L\ wastewater}\right]}{\left[\frac{g\ carbon_{wastewater}}{L\ wastewater}\right]} \quad \text{Equation S5}$$

$$= \frac{15.83}{24.30} = \mathbf{0.65} \quad \text{Min.}$$

$$= \frac{20.73}{31.70} = \mathbf{0.65} \quad \text{Max.}$$

Amount of LA produced $[Cmol]$ from wet waste in the U.S. per year: #19 x #1 x #11 x #12 x #13

$$\left[\frac{\boldsymbol{cmol\ LA_{produced}}}{\boldsymbol{year}}\right]_{\boldsymbol{the\ U.S.}}$$

$$= \left[\frac{\cancel{g\ LA_{produced}}}{\cancel{g\ carbon_{wet\ waste}}}\right] \times \left[\frac{\cancel{dry\ short\ ton\ wet\ waste}}{year}\right] \times \left[\frac{\cancel{g\ carbon_{wet\ waste}}}{\cancel{g\ dry\ wet\ waste}} wt.\%\right] \times \left[\frac{\cancel{g}}{\cancel{dry\ short\ ton}}\right] \times \left[\frac{Cmol\ LA}{\cancel{g\ LA}}\right] \quad \text{Equation S6}$$

$$= 0.65 \times 35\ million \times 38.8\% \times 0.907\ million \times 0.033 = \mathbf{0.26 \times 10^{12}} \quad \text{Min.}$$

$$= 0.65 \times 47\ million \times 76.3\% \times 0.907\ million \times 0.033 = \mathbf{0.70 \times 10^{12}} \quad \text{Max.}$$

#### Calculating the amount of sugar $[Cmol]$ generated by agriculture per year in the U.S.

In 2024/2025, the amount of sugar produced in the U.S. is 8.42 million metric tons (https://www.fas.usda.gov/data/production/commodity/0612000). Assuming that the sugar is sucrose, then the C-mol equivalent of the sugar can be calculated using parameter #14 as shown in Equation S7.

$$\left[\frac{\boldsymbol{Cmol\ sugar_{produced}}}{\boldsymbol{year}}\right]_{\boldsymbol{the\ U.S.}}$$

$$= \frac{8.42\ million\ metric\ tons\ sugar}{year} \times \frac{10^6\ g}{metric\ ton} \times 0.035 \frac{Cmol\ sugar}{g\ sugar} = \mathbf{0.29 \times 10^{12}} \quad \text{Equation S7}$$

**Figure S1. Potentials of wet waste-derived lactic acid as a feedstock for yeast-based biorefineries.**

The grey background identifies a chemical process. The green background identifies a microbial process and the green font identifies the corresponding microbe(s). Thick red lines surround the areas where this study's findings can potentially contribute. The engineering of the primary metabolism of the yeast(s) is optional from the perspective of single-cell protein and yeast extract production. In contrast, an engineered metabolism is imperative for the production of non-native secondary metabolites such as designer hydrocarbons for use as drop-in and performance-advantaged transportation fuel components [4,5].

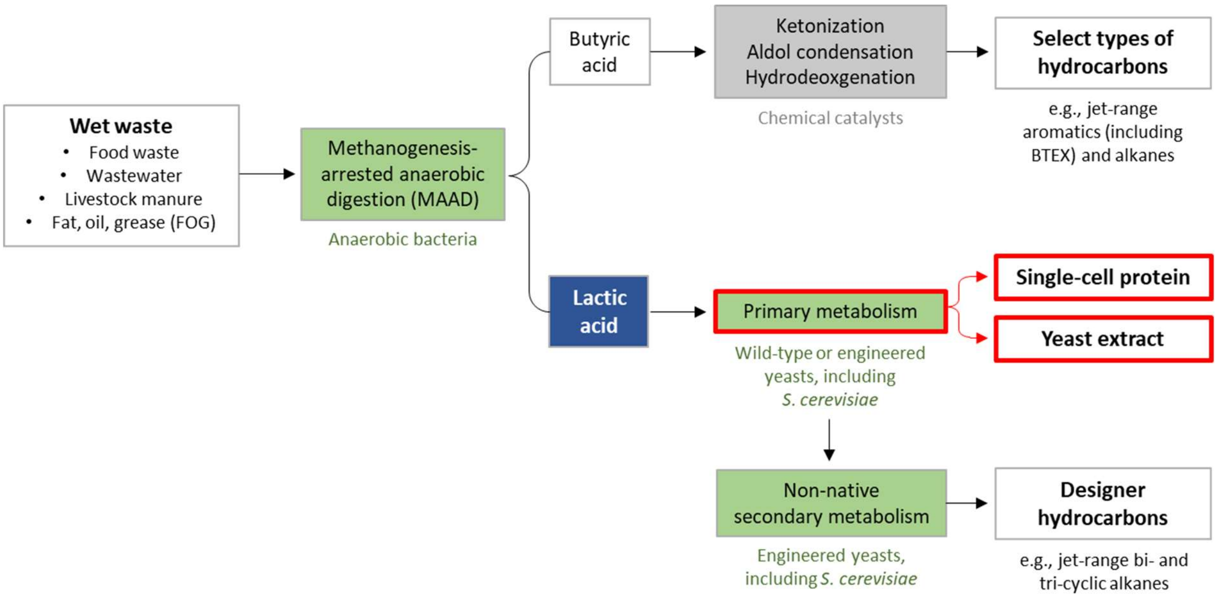

#### Note S2. Delimitation of the use of “lactic acid” versus “lactate” in this manuscript.

In this manuscript, the term “lactic acid” refers to the compound in the MAAD product stream, whereas the term “lactate” refers to the compound present in the defined minimal medium (pH 6.0) used for cultivating yeasts.

#### Table S3. Growth performance of several *S. cerevisiae* strains on lactate relative to their growth on glucose as reported in the literature.

The values listed represent the average of two or more biological replicates as described in the original study. “Total growth” is defined as the difference between the initial and the maximum $OD_{600}$ equivalent values of the culture as observed throughout the cultivation course [6]. “Total-growth rate” is defined as the total growth divided by the hour needed by the culture to reach the total growth.

| Strain | $[glucose]$ | $[lactate]$ | Growth parameter $x$ | Unit | $x_{glucose}$ | $x_{lactate}$ | $\frac{x_{lactate}}{x_{glucose}}$ | Reference(s) |
|---|---|---|---|---|---|---|---|---|
| CBS8066 | 1.0% w/v | 0.5% w/v | Specific growth rate | $[h^{-1}]$ | 0.44 | < 0.05 or absent | Negligible | Table 2, Section 3.1 in [7] |
| BAY.17 | | | | | 0.42 | | | |
| X2180 | | | | | 0.34 | | | |
| CEN.PK122 | | | | | 0.41 | | | |
| YPH499 | 0.5% w/v | 0.5% w/v | Doubling time | $[h]$ | 3.5 | 5.0 | 143% | Table 2 in [8] |
| W303-1a | | | | | 2.0 | 4.5 | 225% | |
| CEN.PK113-7D | 2.0% w/v | 2.0% w/v | Total growth | $\begin{bmatrix} OD_{600} \\ equiv. \end{bmatrix}$ | 19.38 | 6.61 | 34% | Figures 5R.2.A, 5R.3.A, 5R.11.A, 5R.12.A in [9] |
| | | | Total-growth rate | $[h^{-1}]$ | 0.321 | 0.058 | 18% | |
| | | | Maximum specific growth rate | $[h^{-1}]$ | 0.513 | 0.414 | 81 % | |

#### Figure S2. Growth profiles of *S. cerevisiae* CEN.PK113-7D in YM-21, YM-25, and YM-25 + GLY.

Adapted from the raw data forming Figures 5R.1 and 5R.10 in [9]. The pink background indicates precultivation in liquid YM-10. **A**, Linear-scale growth curves. **B**, Semilog-scale growth curves. **C**, Curves of growth relative to that in YM-21. Shown are the aggregated growth profiles of a representative biological replicate (i.e., Clone 2), which are very similar to those of the other two clones.

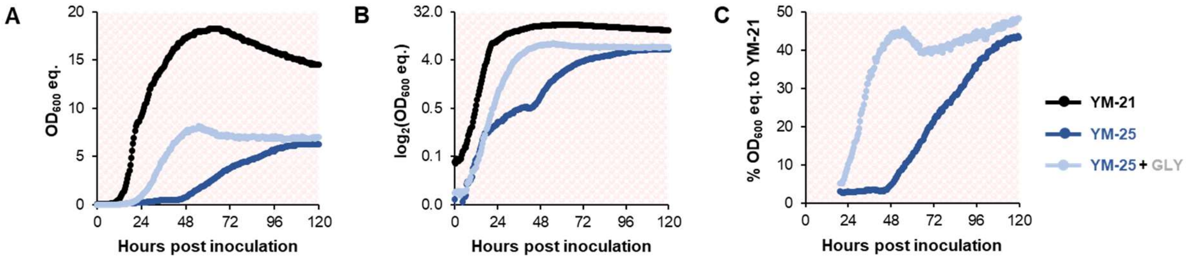

**Table S4. Major differences between this study and the referenced previous study.**

| Property | The referenced previous study [9] | This study |
|---|---|---|
| Strain(s) | *S. cerevisiae* CEN.PK113-7D (laboratory strain) | 24 natural isolates:<br>• 22 *S. cerevisiae*<br>• 1 *Wickerhamiella versatilis*<br>• 1 *Clavispora lusitaniae* |
| Precultivation medium | Liquid YM-10 | Solid agar media<br>• Potato infusion/dextrose<br>• YM-10 |
| Glycerol/lactate molar ratio | 3.08 (~3.0) | • 0.10 or 0.12 (~0.1)<br>• 0.49 (~0.5)<br>• 2.45 (~2.5) |
| Parallel characterization of growth in the corresponding glycerol minimal media | No | Yes |

**Figure S3. Flowchart of growth profiling experiments conducted in this study.**

A detailed description of the experiments is presented in the Methods section.

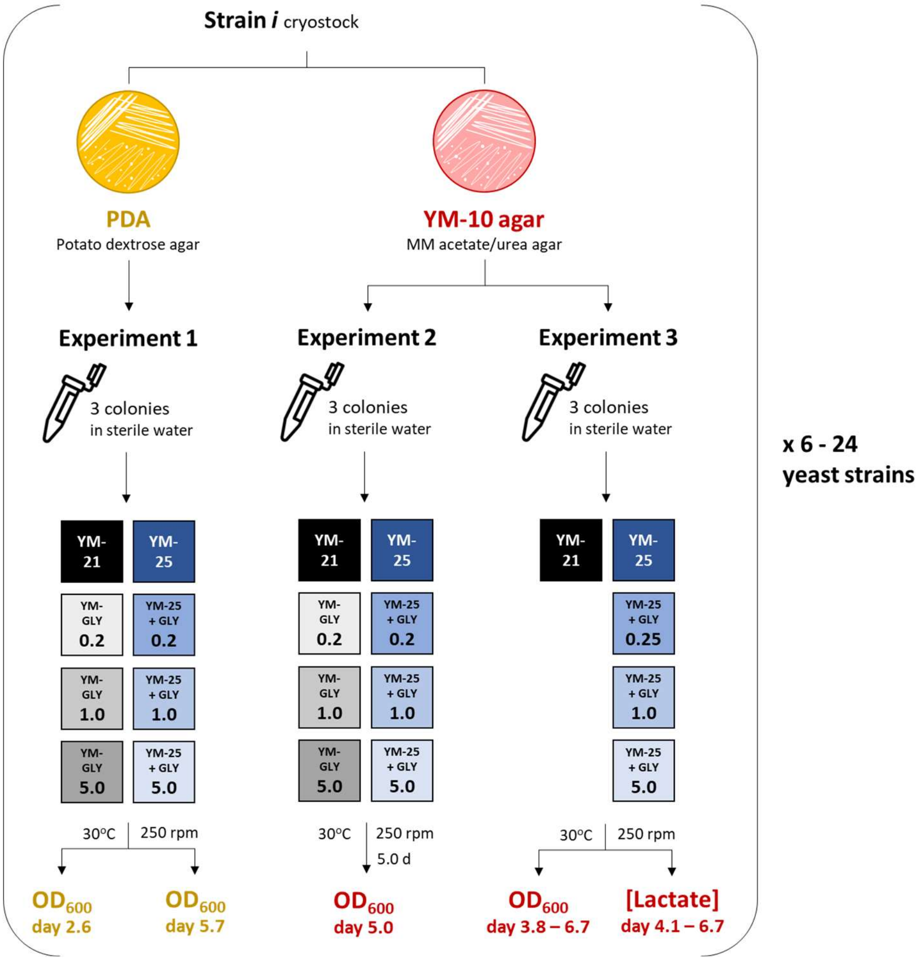

**Figure S4. Photographs of the 24-well cultivation plates from Experiment 2 at the end of cultivation.**

The bold blue font indicates "Lactate +" *S. cerevisiae* strains defined in Figure 3A. The bold red font indicates non-*S. cerevisiae* strains.

***Strains and media placement map***

| <> | 1 | 2 | 3 | 4 | 5 | 6 |
|---|---|---|---|---|---|---|
| | Strain (St.) 1 | | St. 2 | | St. 3 | |
| A | YM-21 | YM-25 | | | | |
| B | YM-GLY$_{0.2}$ | YM-25 + GLY$_{0.2}$ | | | | |
| C | YM-GLY$_{1.0}$ | YM-25 + GLY$_{1.0}$ | | | | |
| D | YM-GLY$_{5.0}$ | YM-25 + GLY$_{5.0}$ | | | | |

Y649 – Y655 – Y682

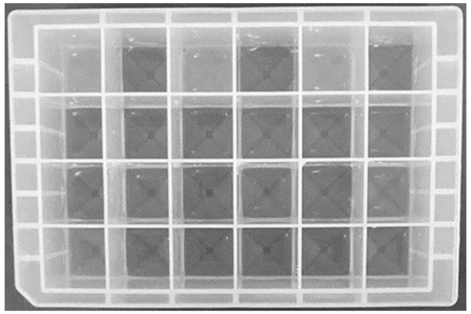

**Y683** – Y743 – Y93

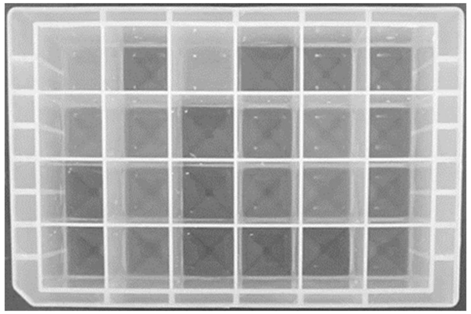

**Y1061** – Y1062 – Y1111

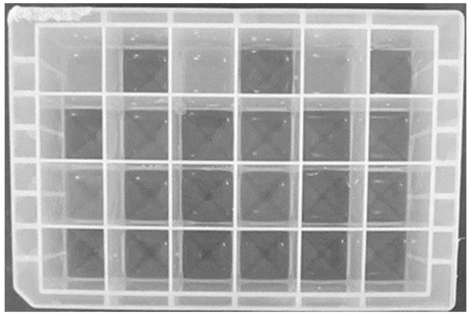

Y1122 – **Y1125** – Y1129

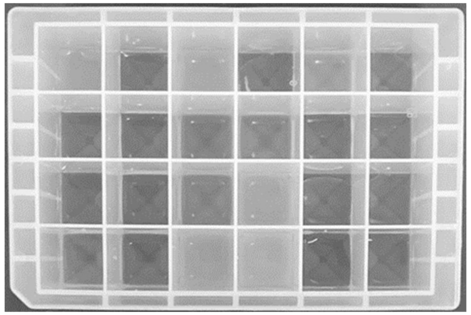

Y1136 – Y1139 – **Y1242**

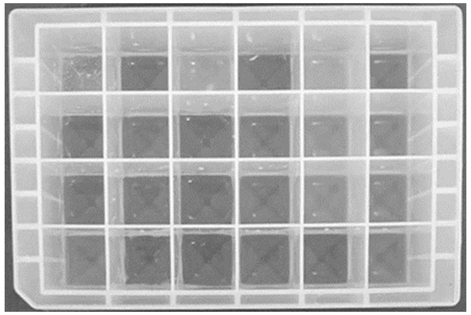

**Y1584** – **Y1613** – **Y1624**

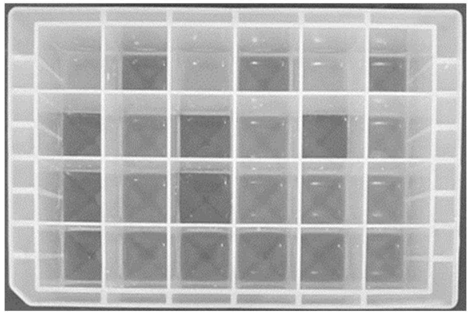

Y1672 – Y1674 – Y1703

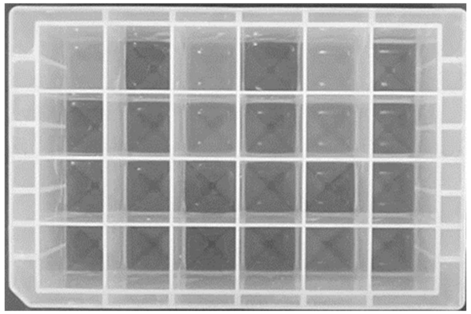

Y1705 - **Y1728** - Y634

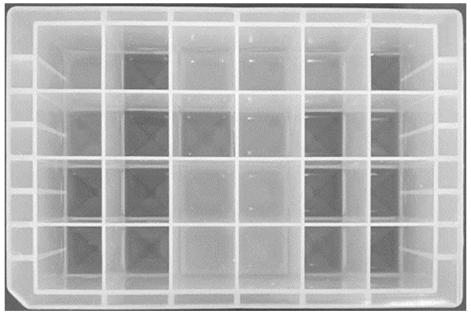

#### Figure S5. Microtiter plate reader-assisted $OD_{600}$ measurement of cultures from Experiment 1 – day 2.6.

**A**, Placement map of cultures of the same strain on a 96-well microtiter plate. The sequence and orientation are maintained and repeated throughout the plate as shown by the color code. **B** and **D**, Placement map of cultures of twelve different strains on microtiter plates 1 and 2, respectively. **C** and **E**, $OD_{600}$ readout of samples on microtiter plates 1 and 2, respectively. The values were substracted with $OD_{600}$ of distilled water in the same position (Figure S8) to obtain $OD_{600}$ of the corresponding cultures.

**A** **Medium placement map**

| <> | 1 | 2 | 3 | 4 | 5 | 6 | 7 | 8 | 9 | 10 | 11 | 12 |
|---|---|---|---|---|---|---|---|---|---|---|---|---|
| A | YM-21 | YM-25 | | | | | | | | | | |
| B | YM-GLY$_{0.2}$ | YM-25 + GLY$_{0.2}$ | | | | | | | | | | |
| C | YM-GLY$_{1.0}$ | YM-25 + GLY$_{1.0}$ | | | | | | | | | | |
| D | YM-GLY$_{5.0}$ | YM-25 + GLY$_{5.0}$ | | | | | | | | | | |
| E | | | | | | | | | | | | |
| F | | | | | | | | | | | | |
| G | | | | | | | | | | | | |
| H | | | | | | | | | | | | |

**B** **Strain placement map of plate 1**

<table>
<tr><th><></th><th>1</th><th>2</th><th>3</th><th>4</th><th>5</th><th>6</th><th>7</th><th>8</th><th>9</th><th>10</th><th>11</th><th>12</th></tr>
<tr><td>A</td><td colspan="2" rowspan="4">Y683</td><td colspan="2" rowspan="4">Y1242</td><td colspan="2" rowspan="4">Y1061</td><td colspan="2" rowspan="4">Y1111</td><td colspan="2" rowspan="4">Y1122</td><td colspan="2" rowspan="4">Y634</td></tr>
<tr><td>B</td></tr>
<tr><td>C</td></tr>
<tr><td>D</td></tr>
<tr><td>E</td><td colspan="2" rowspan="4">Y93</td><td colspan="2" rowspan="4">Y1125</td><td colspan="2" rowspan="4">Y1728</td><td colspan="2" rowspan="4">Y743</td><td colspan="2" rowspan="4">Y1139</td><td colspan="2" rowspan="4">Y1062</td></tr>
<tr><td>F</td></tr>
<tr><td>G</td></tr>
<tr><td>H</td></tr>
</table>

**C** **$OD_{600}$ of plate 1**

| <> | 1 | 2 | 3 | 4 | 5 | 6 | 7 | 8 | 9 | 10 | 11 | 12 |
|---|---|---|---|---|---|---|---|---|---|---|---|---|
| A | 1.2857 | 0.1227 | 1.1884 | 0.0618 | 1.0437 | 0.0597 | 0.6854 | 0.0645 | 0.8187 | 0.0573 | 0.9960 | 0.0746 |
| B | 0.0494 | 0.7893 | 0.1224 | 0.1269 | 0.0361 | 0.5908 | 0.0377 | 0.0570 | 0.0575 | 0.0413 | 0.1643 | 0.0731 |
| C | 0.0497 | 0.8231 | 0.0588 | 0.1641 | 0.0501 | 0.5588 | 0.0499 | 0.0683 | 0.0490 | 0.0486 | 0.0526 | 0.1347 |
| D | 0.0630 | 0.8953 | 0.0519 | 0.1445 | 0.0398 | 0.5880 | 0.0448 | 0.0728 | 0.0431 | 0.0574 | 0.0491 | 0.1013 |
| E | 0.0457 | 0.1068 | 0.9875 | 0.0494 | 1.4718 | 0.2105 | 1.2969 | 0.0611 | 1.0579 | 0.0408 | 1.3503 | 0.0556 |
| F | 0.2336 | 0.0860 | 0.2457 | 0.2256 | 0.5694 | 0.9338 | 0.0527 | 0.0963 | 0.0387 | 0.0397 | 0.0439 | 0.1211 |
| G | 0.0406 | 0.0809 | 0.2507 | 0.2000 | 1.4295 | 1.4798 | 0.0530 | 0.1152 | 0.0489 | 0.0390 | 0.0566 | 0.1614 |
| H | 0.0462 | 0.0825 | 0.2498 | 0.3793 | 1.5771 | 1.6703 | 0.0678 | 0.1859 | 0.0485 | 0.0675 | 0.0755 | 0.3171 |

**D** **Strain placement map of plate 2**

<table>
<tr><th><></th><th>1</th><th>2</th><th>3</th><th>4</th><th>5</th><th>6</th><th>7</th><th>8</th><th>9</th><th>10</th><th>11</th><th>12</th></tr>
<tr><td>A</td><td colspan="2" rowspan="4">Y649</td><td colspan="2" rowspan="4">Y1584</td><td colspan="2" rowspan="4">Y1613</td><td colspan="2" rowspan="4">Y1624</td><td colspan="2" rowspan="4">Y1674</td><td colspan="2" rowspan="4">Y1703</td></tr>
<tr><td>B</td></tr>
<tr><td>C</td></tr>
<tr><td>D</td></tr>
<tr><td>E</td><td colspan="2" rowspan="4">Y655</td><td colspan="2" rowspan="4">Y682</td><td colspan="2" rowspan="4">Y1671</td><td colspan="2" rowspan="4">Y1705</td><td colspan="2" rowspan="4">Y1129</td><td colspan="2" rowspan="4">Y1136</td></tr>
<tr><td>F</td></tr>
<tr><td>G</td></tr>
<tr><td>H</td></tr>
</table>

**E** **$OD_{600}$ of plate 2**

| <> | 1 | 2 | 3 | 4 | 5 | 6 | 7 | 8 | 9 | 10 | 11 | 12 |
|---|---|---|---|---|---|---|---|---|---|---|---|---|
| A | 0.9896 | 0.0509 | 1.2896 | 0.1199 | 1.2801 | 0.1061 | 1.3027 | 0.1122 | 1.1612 | 0.0428 | 1.2679 | 0.0604 |
| B | 0.0758 | 0.1856 | 0.0447 | 0.8194 | 0.0683 | 0.7901 | 0.0530 | 0.7375 | 0.0519 | 0.0382 | 0.0419 | 0.0420 |
| C | 0.0440 | 0.2766 | 0.0555 | 0.8993 | 0.0615 | 0.8459 | 0.0515 | 0.9001 | 0.0485 | 0.0507 | 0.0609 | 0.0586 |
| D | 0.0576 | 0.1464 | 0.0695 | 0.7887 | 0.0597 | 0.8352 | 0.0634 | 0.8713 | 0.0421 | 0.1194 | 0.0546 | 0.0801 |
| E | 1.0968 | 0.0536 | 1.2104 | 0.0542 | 1.2254 | 0.0439 | 1.2773 | 0.0499 | 0.9611 | 0.0371 | 0.6926 | 0.0505 |
| F | 0.0482 | 0.2299 | 0.3791 | 0.0798 | 0.0413 | 0.0628 | 0.0543 | 0.0824 | 0.0568 | 0.0350 | 0.0365 | 0.4011 |
| G | 0.0418 | 0.0752 | 0.0516 | 0.1327 | 0.0547 | 0.0560 | 0.0607 | 0.0918 | 0.0413 | 0.0354 | 0.0463 | 0.1485 |
| H | 0.0514 | 0.0617 | 0.0615 | 0.1300 | 0.0567 | 0.0772 | 0.0662 | 0.1552 | 0.0375 | 0.0357 | 0.0497 | 0.2185 |

#### Figure S6. Microtiter plate reader-assisted $OD_{600}$ measurement of cultures from Experiment 1 – day 5.7.

**A**, Placement map of cultures of the same strain on a 96-well microtiter plate. The sequence and orientation are maintained and repeated throughout the plate as shown by the color code. **B** and **D**, Placement map of cultures of twelve different strains on microtiter plates 1 and 2, respectively. **C** and **E**, $OD_{600}$ readout of samples on microtiter plates 1 and 2, respectively. The values were substracted with $OD_{600}$ of distilled water in the same position (Figure S8) to obtain $OD_{600}$ of the corresponding cultures.

##### A Medium placement map

| <> | 1 | 2 | 3 | 4 | 5 | 6 | 7 | 8 | 9 | 10 | 11 | 12 |
|---|---|---|---|---|---|---|---|---|---|---|---|---|
| A | YM-21 | YM-25 | | | | | | | | | | |
| B | YM-GLY$_{0.2}$ | YM-25 + GLY$_{0.2}$ | | | | | | | | | | |
| C | YM-GLY$_{1.0}$ | YM-25 + GLY$_{1.0}$ | | | | | | | | | | |
| D | YM-GLY$_{5.0}$ | YM-25 + GLY$_{5.0}$ | | | | | | | | | | |
| E | | | | | | | | | | | | |
| F | | | | | | | | | | | | |
| G | | | | | | | | | | | | |
| H | | | | | | | | | | | | |

##### B Strain placement map of plate 1

| <> | 1 | 2 | 3 | 4 | 5 | 6 | 7 | 8 | 9 | 10 | 11 | 12 |
|---|---|---|---|---|---|---|---|---|---|---|---|---|
| A | Y93 | | Y1125 | | Y1728 | | Y743 | | Y1139 | | Y1062 | |
| B | | | | | | | | | | | | |
| C | | | | | | | | | | | | |
| D | | | | | | | | | | | | |
| E | Y683 | | Y1242 | | Y1061 | | Y1111 | | Y1122 | | Y634 | |
| F | | | | | | | | | | | | |
| G | | | | | | | | | | | | |
| H | | | | | | | | | | | | |

##### C $OD_{600}$ of plate 1

| <> | 1 | 2 | 3 | 4 | 5 | 6 | 7 | 8 | 9 | 10 | 11 | 12 |
|---|---|---|---|---|---|---|---|---|---|---|---|---|
| A | 0.0896 | 0.2799 | 1.3396 | 0.0791 | 1.6220 | 1.3457 | 1.3346 | 0.0760 | 1.2176 | 0.0523 | 1.3980 | 0.0702 |
| B | 0.5795 | 0.5380 | 0.7080 | 0.5768 | 0.7757 | 1.5338 | 0.8529 | 0.9213 | 0.0409 | 0.1901 | 0.0589 | 0.8689 |
| C | 0.0506 | 0.5556 | 1.4583 | 1.3361 | 1.5470 | 1.6964 | 0.0897 | 0.9652 | 0.0467 | 0.0999 | 0.0915 | 0.8711 |
| D | 0.1324 | 0.5062 | 1.6474 | 1.8395 | 1.9087 | 2.0313 | 0.1725 | 0.6578 | 0.0735 | 0.2830 | 0.2816 | 0.7671 |
| E | 1.4549 | 0.7528 | 1.4210 | 0.6313 | 1.2541 | 0.2078 | 0.7802 | 0.0738 | 0.9517 | 0.0546 | 1.3092 | 0.0614 |
| F | 0.0676 | 1.1861 | 0.6084 | 0.8730 | 0.0408 | 0.9433 | 0.0548 | 0.5714 | 0.0703 | 0.5380 | 0.6788 | 0.6568 |
| G | 0.0689 | 1.2125 | 0.1020 | 0.9259 | 0.0471 | 0.8616 | 0.0531 | 0.6774 | 0.0499 | 0.2236 | 0.0699 | 0.7222 |
| H | 0.1948 | 1.0688 | 0.1415 | 0.7862 | 0.1036 | 0.7986 | 0.0642 | 0.1160 | 0.0482 | 0.2183 | 0.1143 | 0.8243 |

##### D Strain placement map of plate 2

| <> | 1 | 2 | 3 | 4 | 5 | 6 | 7 | 8 | 9 | 10 | 11 | 12 |
|---|---|---|---|---|---|---|---|---|---|---|---|---|
| A | Y655 | | Y682 | | Y1671 | | Y1705 | | Y1129 | | Y1136 | |
| B | | | | | | | | | | | | |
| C | | | | | | | | | | | | |
| D | | | | | | | | | | | | |
| E | Y649 | | Y1584 | | Y1613 | | Y1624 | | Y1674 | | Y1703 | |
| F | | | | | | | | | | | | |
| G | | | | | | | | | | | | |
| H | | | | | | | | | | | | |

##### E $OD_{600}$ of plate 2

| <> | 1 | 2 | 3 | 4 | 5 | 6 | 7 | 8 | 9 | 10 | 11 | 12 |
|---|---|---|---|---|---|---|---|---|---|---|---|---|
| A | 1.1427 | 0.0626 | 1.2764 | 0.0600 | 1.3817 | 0.0708 | 1.3534 | 0.0792 | 1.1225 | 0.0363 | 0.8894 | 0.0563 |
| B | 0.1239 | 0.2237 | 0.7249 | 0.6556 | 0.0689 | 0.1986 | 0.5917 | 0.5325 | 0.0713 | 0.0339 | 0.0468 | 0.6226 |
| C | 0.0556 | 0.3099 | 0.0744 | 0.7213 | 0.0822 | 0.0780 | 0.1177 | 1.2381 | 0.0532 | 0.0669 | 0.1246 | 0.6252 |
| D | 0.1125 | 0.0889 | 0.0902 | 0.7916 | 0.1388 | 0.1816 | 0.7595 | 0.9510 | 0.0385 | 0.0371 | 0.5368 | 0.1090 |
| E | 1.2233 | 0.0647 | 1.4889 | 1.0988 | 1.4917 | 0.9896 | 1.5262 | 0.9575 | 1.3218 | 0.0507 | 1.4285 | 0.0415 |
| F | 0.6527 | 0.4979 | 0.0762 | 1.2505 | 0.7425 | 1.2086 | 0.0941 | 1.2197 | 0.6544 | 0.0725 | 0.0515 | 0.0488 |
| G | 0.0775 | 0.7177 | 0.0700 | 1.1642 | 0.0858 | 1.1367 | 0.0871 | 1.2121 | 0.0529 | 0.3105 | 0.0526 | 0.1153 |
| H | 0.2699 | 0.9608 | 0.2355 | 1.0548 | 0.2015 | 1.1286 | 0.1889 | 1.1605 | 0.0701 | 0.2195 | 0.1176 | 0.1670 |

#### Figure S7. Microtiter plate reader-assisted $OD_{600}$ measurement of cultures from Experiment 2 – day 5.0.

**A**, Placement map of cultures of the same strain on a 96-well microtiter plate. The sequence and orientation are maintained and repeated throughout the plate as shown by the color code. **B** and **D**, Placement map of cultures of twelve different strains on microtiter plates 1 and 2, respectively. **C** and **E**, $OD_{600}$ readout of samples on microtiter plates 1 and 2, respectively. The values were substracted with $OD_{600}$ of distilled water in the same position (Figure S8) to obtain $OD_{600}$ of the corresponding cultures.

**A** **Medium placement map**

| <> | 1 | 2 | 3 | 4 | 5 | 6 | 7 | 8 | 9 | 10 | 11 | 12 |
|---|---|---|---|---|---|---|---|---|---|---|---|---|
| A | YM-21 | YM-25 | | | | | | | | | | |
| B | YM-$GLY_{0.2}$ | YM-25 + $GLY_{0.2}$ | | | | | | | | | | |
| C | YM-$GLY_{1.0}$ | YM-25 + $GLY_{1.0}$ | | | | | | | | | | |
| D | YM-$GLY_{5.0}$ | YM-25 + $GLY_{5.0}$ | | | | | | | | | | |
| E | | | | | | | | | | | | |
| F | | | | | | | | | | | | |
| G | | | | | | | | | | | | |
| H | | | | | | | | | | | | |

**B** **Strain placement map of plate 1**

<table>
<tr><th><></th><th>1</th><th>2</th><th>3</th><th>4</th><th>5</th><th>6</th><th>7</th><th>8</th><th>9</th><th>10</th><th>11</th><th>12</th></tr>
<tr><td>A</td><td colspan="2" rowspan="4">Y1136</td><td colspan="2" rowspan="4">Y1139</td><td colspan="2" rowspan="4">Y1242</td><td colspan="2" rowspan="4">Y1584</td><td colspan="2" rowspan="4">Y1613</td><td colspan="2" rowspan="4">Y1624</td></tr>
<tr><td>B</td></tr>
<tr><td>C</td></tr>
<tr><td>D</td></tr>
<tr><td>E</td><td colspan="2" rowspan="4">Y1061</td><td colspan="2" rowspan="4">Y1062</td><td colspan="2" rowspan="4">Y1111</td><td colspan="2" rowspan="4">Y1671</td><td colspan="2" rowspan="4">Y1674</td><td colspan="2" rowspan="4">Y1703</td></tr>
<tr><td>F</td></tr>
<tr><td>G</td></tr>
<tr><td>H</td></tr>
</table>

**C** **$OD_{600}$ of plate 1**

| <> | 1 | 2 | 3 | 4 | 5 | 6 | 7 | 8 | 9 | 10 | 11 | 12 |
|---|---|---|---|---|---|---|---|---|---|---|---|---|
| A | 1.1924 | 0.0794 | 1.3740 | 0.0909 | 1.3937 | 0.2275 | 1.5822 | 0.8451 | 1.5715 | 0.8601 | 1.6096 | 0.5793 |
| B | 0.0559 | 0.2605 | 0.0684 | 0.1562 | 0.8982 | 0.8767 | 0.1144 | 1.1729 | 0.0697 | 1.0634 | 0.0602 | 1.1966 |
| C | 0.0966 | 0.1902 | 0.0668 | 0.2574 | 0.8643 | 0.8487 | 0.1020 | 1.1442 | 0.0763 | 1.1260 | 0.7233 | 1.1906 |
| D | 0.5763 | 0.1679 | 0.0791 | 0.1580 | 0.2395 | 0.6945 | 0.1637 | 1.0056 | 0.4675 | 0.9597 | 0.5578 | 0.9717 |
| E | 1.4626 | 0.1335 | 1.5058 | 0.0947 | 1.2955 | 0.0636 | 1.4330 | 0.0549 | 1.3018 | 0.0456 | 1.2799 | 0.0486 |
| F | 0.0641 | 0.4387 | 0.1021 | 0.2646 | 0.0614 | 0.0564 | 0.0606 | 0.2757 | 0.7948 | 0.2326 | 0.7786 | 0.0683 |
| G | 0.7901 | 0.3170 | 0.0904 | 0.7145 | 0.0802 | 0.1731 | 0.1180 | 0.2114 | 0.0658 | 0.2222 | 0.0663 | 0.1040 |
| H | 0.1169 | 0.3201 | 0.1751 | 0.3134 | 0.0608 | 0.1082 | 0.1190 | 0.2476 | 0.0786 | 0.2197 | 0.5583 | 0.1886 |

**D** **Strain placement map of plate 2**

<table>
<tr><th><></th><th>1</th><th>2</th><th>3</th><th>4</th><th>5</th><th>6</th><th>7</th><th>8</th><th>9</th><th>10</th><th>11</th><th>12</th></tr>
<tr><td>A</td><td colspan="2" rowspan="4">Y649</td><td colspan="2" rowspan="4">Y655</td><td colspan="2" rowspan="4">Y682</td><td colspan="2" rowspan="4">Y1122</td><td colspan="2" rowspan="4">Y1125</td><td colspan="2" rowspan="4">Y1129</td></tr>
<tr><td>B</td></tr>
<tr><td>C</td></tr>
<tr><td>D</td></tr>
<tr><td>E</td><td colspan="2" rowspan="4">Y683</td><td colspan="2" rowspan="4">Y743</td><td colspan="2" rowspan="4">Y93</td><td colspan="2" rowspan="4">Y1705</td><td colspan="2" rowspan="4">Y1728</td><td colspan="2" rowspan="4">Y634</td></tr>
<tr><td>F</td></tr>
<tr><td>G</td></tr>
<tr><td>H</td></tr>
</table>

**E** **$OD_{600}$ of plate 2**

| <> | 1 | 2 | 3 | 4 | 5 | 6 | 7 | 8 | 9 | 10 | 11 | 12 |
|---|---|---|---|---|---|---|---|---|---|---|---|---|
| A | 1.3333 | 0.0622 | 1.2992 | 0.0616 | 1.3586 | 0.0637 | 1.4424 | 0.0667 | 1.4762 | 0.0454 | 1.1483 | 0.0441 |
| B | 0.0547 | 0.0552 | 0.0493 | 0.1420 | 0.0554 | 0.0718 | 0.0512 | 0.0588 | 0.6496 | 0.7396 | 0.0401 | 0.0366 |
| C | 0.0487 | 0.3886 | 0.0545 | 0.2608 | 0.1186 | 0.6328 | 0.0583 | 0.1223 | 0.6893 | 1.5069 | 0.0412 | 0.0395 |
| D | 0.4134 | 0.2117 | 0.0791 | 0.0947 | 0.0758 | 0.0622 | 0.3458 | 0.2380 | 1.6740 | 1.8659 | 0.0378 | 0.0390 |
| E | 1.6100 | 0.5211 | 1.5091 | 0.0727 | 0.1607 | 0.1510 | 1.4967 | 0.0735 | 1.6752 | 1.3934 | 1.2565 | 0.0586 |
| F | 0.9253 | 1.1649 | 0.0905 | 0.4486 | 0.7944 | 0.2277 | 0.8960 | 0.1013 | 0.7687 | 1.3413 | 0.1028 | 0.2876 |
| G | 0.1084 | 1.1639 | 0.0750 | 0.5374 | 0.6956 | 0.2823 | 0.1058 | 0.0962 | 1.6585 | 1.7444 | 0.0672 | 0.0708 |
| H | 0.1415 | 0.9624 | 0.1203 | 0.1019 | 0.0988 | 0.2727 | 0.1689 | 0.4787 | 1.9234 | 1.9640 | 0.5346 | 0.0560 |

**Figure S8. $OD_{600}$ of distilled water used as the well-specific blank for any microtiter plate reader-assisted culture $OD_{600}$ measurement in this study.**

The values were used to subtract the $OD_{600}$ readout of yeast culture samples in the same well, thus resulting in the culture's $OD_{600}$.

| <> | 1 | 2 | 3 | 4 | 5 | 6 | 7 | 8 | 9 | 10 | 11 | 12 |
|---|---|---|---|---|---|---|---|---|---|---|---|---|
| A | 0.0336 | 0.0359 | 0.0343 | 0.0362 | 0.0371 | 0.0390 | 0.0361 | 0.0384 | 0.0336 | 0.0365 | 0.0375 | 0.0419 |
| B | 0.0351 | 0.0352 | 0.0354 | 0.0331 | 0.0338 | 0.0336 | 0.0357 | 0.0351 | 0.0359 | 0.0332 | 0.0350 | 0.0326 |
| C | 0.0348 | 0.0373 | 0.0372 | 0.0368 | 0.0363 | 0.0382 | 0.0357 | 0.0368 | 0.0364 | 0.0363 | 0.0391 | 0.0385 |
| D | 0.0347 | 0.0357 | 0.0353 | 0.0349 | 0.0347 | 0.0570 | 0.0349 | 0.0357 | 0.0356 | 0.0356 | 0.0346 | 0.0334 |
| E | 0.0340 | 0.0370 | 0.0368 | 0.0344 | 0.0382 | 0.0353 | 0.0348 | 0.0376 | 0.0360 | 0.0347 | 0.0348 | 0.0350 |
| F | 0.0359 | 0.0354 | 0.0357 | 0.0346 | 0.0332 | 0.0339 | 0.0356 | 0.0368 | 0.0367 | 0.0345 | 0.0332 | 0.0334 |
| G | 0.0347 | 0.0381 | 0.0382 | 0.0351 | 0.0360 | 0.0362 | 0.0369 | 0.0368 | 0.0374 | 0.0343 | 0.0363 | 0.0343 |
| H | 0.0355 | 0.0382 | 0.0355 | 0.0351 | 0.0364 | 0.0373 | 0.0367 | 0.0359 | 0.0362 | 0.0356 | 0.0355 | 0.0367 |

#### Table S5. $OD_{600}$ of cultures from Experiment 1 – day 2.6.

The result of cell-wise subtraction of distilled water's $OD_{600}$ (Figure S8) from the corresponding samples' $OD_{600}$ (Figures S5C and S5E).

| Strain | YM-21 | YM-GLY$_{0.2}$ | YM-GLY$_{1.0}$ | YM-GLY$_{5.0}$ | YM-25 | YM-25 + GLY$_{0.2}$ | YM-25 + GLY$_{1.0}$ | YM-25 + GLY$_{5.0}$ |
|---|---|---|---|---|---|---|---|---|
| Y93 | 0.0117 | 0.1977 | 0.0059 | 0.0107 | 0.0698 | 0.0506 | 0.0428 | 0.0443 |
| Y634 | 0.9585 | 0.1293 | 0.0135 | 0.0145 | 0.0327 | 0.0405 | 0.0962 | 0.0679 |
| Y649 | 0.9560 | 0.0407 | 0.0092 | 0.0229 | 0.0150 | 0.1504 | 0.2393 | 0.1107 |
| Y655 | 1.0628 | 0.0123 | 0.0071 | 0.0159 | 0.0166 | 0.1945 | 0.0371 | 0.0235 |
| Y682 | 1.1736 | 0.3434 | 0.0134 | 0.0260 | 0.0198 | 0.0452 | 0.0976 | 0.0949 |
| Y683 | 1.2521 | 0.0143 | 0.0149 | 0.0283 | 0.0868 | 0.7541 | 0.7858 | 0.8596 |
| Y743 | 1.2621 | 0.0171 | 0.0161 | 0.0311 | 0.0235 | 0.0595 | 0.0784 | 0.1500 |
| Y1061 | 1.0066 | 0.0023 | 0.0138 | 0.0051 | 0.0207 | 0.5572 | 0.5206 | 0.5310 |
| Y1062 | 1.3155 | 0.0107 | 0.0203 | 0.0400 | 0.0206 | 0.0877 | 0.1271 | 0.2804 |
| Y1111 | 0.6493 | 0.0020 | 0.0142 | 0.0099 | 0.0261 | 0.0219 | 0.0315 | 0.0371 |
| Y1122 | 0.7851 | 0.0216 | 0.0126 | 0.0075 | 0.0208 | 0.0081 | 0.0123 | 0.0218 |
| Y1125 | 0.9507 | 0.2100 | 0.2125 | 0.2143 | 0.0150 | 0.1910 | 0.1649 | 0.3442 |
| Y1129 | 0.9251 | 0.0201 | 0.0039 | 0.0013 | 0.0024 | 0.0005 | 0.0011 | 0.0001 |
| Y1136 | 0.6578 | 0.0033 | 0.0100 | 0.0142 | 0.0155 | 0.3677 | 0.1142 | 0.1818 |
| Y1139 | 1.0219 | 0.0020 | 0.0115 | 0.0123 | 0.0061 | 0.0052 | 0.0047 | 0.0319 |
| Y1242 | 1.1541 | 0.0870 | 0.0216 | 0.0166 | 0.0256 | 0.0938 | 0.1273 | 0.1096 |
| Y1584 | 1.2553 | 0.0093 | 0.0183 | 0.0342 | 0.0837 | 0.7863 | 0.8625 | 0.7538 |
| Y1613 | 1.2430 | 0.0345 | 0.0252 | 0.0250 | 0.0671 | 0.7565 | 0.8077 | 0.7782 |
| Y1624 | 1.2666 | 0.0173 | 0.0158 | 0.0285 | 0.0738 | 0.7024 | 0.8633 | 0.8356 |
| Y1671 | 1.1872 | 0.0081 | 0.0187 | 0.0203 | 0.0086 | 0.0289 | 0.0198 | 0.0399 |
| Y1674 | 1.1276 | 0.0160 | 0.0121 | 0.0065 | 0.0063 | 0.0050 | 0.0144 | 0.0838 |
| Y1703 | 1.2304 | 0.0069 | 0.0218 | 0.0200 | 0.0185 | 0.0094 | 0.0201 | 0.0467 |
| Y1705 | 1.2425 | 0.0187 | 0.0238 | 0.0295 | 0.0123 | 0.0456 | 0.0550 | 0.1193 |
| Y1728 | 1.4336 | 0.5362 | 1.3935 | 1.5407 | 0.1752 | 0.8999 | 1.4436 | 1.6330 |

#### Table S6. $OD_{600}$ of cultures from Experiment 1 – day 5.7.

The result of cell-wise subtraction of distilled water's $OD_{600}$ (Figure S8) from the corresponding samples' $OD_{600}$ (Figures S6C and S6E).

| Strain | YM-21 | YM-GLY$_{0.2}$ | YM-GLY$_{1.0}$ | YM-GLY$_{5.0}$ | YM-25 | YM-25 + GLY$_{0.2}$ | YM-25 + GLY$_{1.0}$ | YM-25 + GLY$_{5.0}$ |
|---|---|---|---|---|---|---|---|---|
| Y93 | 0.0560 | 0.5444 | 0.0158 | 0.0977 | 0.2440 | 0.5028 | 0.5183 | 0.4705 |
| Y634 | 1.2744 | 0.6456 | 0.0336 | 0.0788 | 0.0264 | 0.6234 | 0.6879 | 0.7876 |
| Y649 | 1.1893 | 0.6168 | 0.0428 | 0.2344 | 0.0277 | 0.4625 | 0.6796 | 0.9226 |
| Y655 | 1.1091 | 0.0888 | 0.0208 | 0.0778 | 0.0267 | 0.1885 | 0.2726 | 0.0532 |
| Y682 | 1.2421 | 0.6895 | 0.0372 | 0.0549 | 0.0238 | 0.6225 | 0.6845 | 0.7567 |
| Y683 | 1.4209 | 0.0317 | 0.0342 | 0.1593 | 0.7158 | 1.1507 | 1.1744 | 1.0306 |
| Y743 | 1.2985 | 0.8172 | 0.0540 | 0.1376 | 0.0376 | 0.8862 | 0.9284 | 0.6221 |
| Y1061 | 1.2159 | 0.0076 | 0.0111 | 0.0672 | 0.1725 | 0.9094 | 0.8254 | 0.7613 |
| Y1062 | 1.3605 | 0.0239 | 0.0524 | 0.2470 | 0.0283 | 0.8363 | 0.8326 | 0.7337 |
| Y1111 | 0.7454 | 0.0192 | 0.0162 | 0.0275 | 0.0362 | 0.5346 | 0.6406 | 0.0801 |
| Y1122 | 0.9157 | 0.0336 | 0.0125 | 0.0120 | 0.0199 | 0.5035 | 0.1893 | 0.1827 |
| Y1125 | 1.3053 | 0.6726 | 1.4211 | 1.6121 | 0.0429 | 0.5437 | 1.2993 | 1.8046 |
| Y1129 | 1.0889 | 0.0354 | 0.0168 | 0.0029 | -0.0002 | 0.0007 | 0.0306 | 0.0015 |
| Y1136 | 0.8519 | 0.0118 | 0.0855 | 0.5022 | 0.0144 | 0.5900 | 0.5867 | 0.0756 |
| Y1139 | 1.1840 | 0.0050 | 0.0103 | 0.0379 | 0.0158 | 0.1569 | 0.0636 | 0.2474 |
| Y1242 | 1.3842 | 0.5727 | 0.0638 | 0.1060 | 0.5969 | 0.8384 | 0.8908 | 0.7511 |
| Y1584 | 1.4521 | 0.0405 | 0.0318 | 0.2000 | 1.0644 | 1.2159 | 1.1291 | 1.0197 |
| Y1613 | 1.4535 | 0.7093 | 0.0498 | 0.1651 | 0.9543 | 1.1747 | 1.1005 | 1.0913 |
| Y1624 | 1.4914 | 0.0585 | 0.0502 | 0.1522 | 0.9199 | 1.1829 | 1.1753 | 1.1246 |
| Y1671 | 1.3446 | 0.0351 | 0.0459 | 0.1041 | 0.0318 | 0.1650 | 0.0398 | 0.1246 |
| Y1674 | 1.2858 | 0.6177 | 0.0155 | 0.0339 | 0.0160 | 0.0380 | 0.2762 | 0.1839 |
| Y1703 | 1.3937 | 0.0183 | 0.0163 | 0.0821 | 0.0065 | 0.0154 | 0.0810 | 0.1303 |
| Y1705 | 1.3173 | 0.5560 | 0.0820 | 0.7246 | 0.0408 | 0.4974 | 1.2013 | 0.9153 |
| Y1728 | 1.5849 | 0.7419 | 1.5107 | 1.8740 | 1.3067 | 1.5002 | 1.6582 | 1.9743 |

#### Table S7. $OD_{600}$ of cultures from Experiment 2 – day 5.0.

The result of cell-wise subtraction of distilled water's $OD_{600}$ (Figure S8) from the corresponding samples' $OD_{600}$ (Figures S7C and S7E).

| Strain | YM-21 | YM-$GLY_{0.2}$ | YM-$GLY_{1.0}$ | YM-$GLY_{5.0}$ | YM-25 | YM-25 + $GLY_{0.2}$ | YM-25 + $GLY_{1.0}$ | YM-25 + $GLY_{5.0}$ |
|---|---|---|---|---|---|---|---|---|
| Y93 | 0.1225 | 0.7612 | 0.6596 | 0.0624 | 0.1157 | 0.1938 | 0.2461 | 0.2354 |
| Y634 | 1.2217 | 0.0696 | 0.0309 | 0.4991 | 0.0236 | 0.2542 | 0.0365 | 0.0193 |
| Y649 | 1.2997 | 0.0196 | 0.0139 | 0.3787 | 0.0263 | 0.02 | 0.3513 | 0.176 |
| Y655 | 1.2649 | 0.0139 | 0.0173 | 0.0438 | 0.0254 | 0.1089 | 0.224 | 0.0598 |
| Y682 | 1.3215 | 0.0216 | 0.0823 | 0.0411 | 0.0247 | 0.0382 | 0.5946 | 0.0052 |
| Y683 | 1.576 | 0.8894 | 0.0737 | 0.106 | 0.4841 | 1.1295 | 1.1258 | 0.9242 |
| Y743 | 1.4723 | 0.0548 | 0.0368 | 0.0848 | 0.0383 | 0.414 | 0.5023 | 0.0668 |
| Y1061 | 1.4286 | 0.0282 | 0.7554 | 0.0814 | 0.0965 | 0.4033 | 0.2789 | 0.2819 |
| Y1062 | 1.469 | 0.0664 | 0.0522 | 0.1396 | 0.0603 | 0.23 | 0.6794 | 0.2783 |
| Y1111 | 1.2573 | 0.0282 | 0.0442 | 0.0244 | 0.0283 | 0.0225 | 0.1369 | 0.0709 |
| Y1122 | 1.4063 | 0.0155 | 0.0226 | 0.3109 | 0.0283 | 0.0237 | 0.0855 | 0.2023 |
| Y1125 | 1.4426 | 0.6137 | 0.6529 | 1.6384 | 0.0089 | 0.7064 | 1.4706 | 1.8303 |
| Y1129 | 1.1108 | 0.0051 | 0.0021 | 0.0032 | 0.0022 | 0.004 | 0.001 | 0.0056 |
| Y1136 | 1.1588 | 0.0208 | 0.0618 | 0.5416 | 0.0435 | 0.2253 | 0.1529 | 0.1322 |
| Y1139 | 1.3397 | 0.033 | 0.0296 | 0.0438 | 0.0547 | 0.1231 | 0.2206 | 0.1231 |
| Y1242 | 1.3566 | 0.8644 | 0.828 | 0.2048 | 0.1885 | 0.8431 | 0.8105 | 0.6375 |
| Y1584 | 1.5461 | 0.0787 | 0.0663 | 0.1288 | 0.8067 | 1.1378 | 1.1074 | 0.9699 |
| Y1613 | 1.5379 | 0.0338 | 0.0399 | 0.4319 | 0.8236 | 1.0302 | 1.0897 | 0.9241 |
| Y1624 | 1.5721 | 0.0252 | 0.6842 | 0.5232 | 0.5374 | 1.164 | 1.1521 | 0.9383 |
| Y1671 | 1.3982 | 0.025 | 0.0811 | 0.0823 | 0.0173 | 0.2389 | 0.1746 | 0.2117 |
| Y1674 | 1.2658 | 0.7581 | 0.0284 | 0.0424 | 0.0109 | 0.1981 | 0.1879 | 0.1841 |
| Y1703 | 1.2451 | 0.7454 | 0.03 | 0.5228 | 0.0136 | 0.0349 | 0.0697 | 0.1519 |
| Y1705 | 1.4619 | 0.8604 | 0.0689 | 0.1322 | 0.0359 | 0.0645 | 0.0594 | 0.4428 |
| Y1728 | 1.6392 | 0.732 | 1.6211 | 1.8872 | 1.3587 | 1.3068 | 1.7101 | 1.9284 |

#### Table S8. $OD_{600}$ of cultures from Experiment 3 – day 3.8 – 6.7.

The result of cell-wise subtraction of distilled water's $OD_{600}$ (Figure S8) from the corresponding samples' $OD_{600}$. The harvest time, which varies for each strain, is indicated in the first column.

| Day | Strain | YM-21 | YM-25 | YM-25 + $GLY_{0.2}$ | YM-25 + $GLY_{1.0}$ | YM-25 + $GLY_{5.0}$ |
|---|---|---|---|---|---|---|
| 3.8 | Y634 | 0.9243 | 0.0062 | 0.3824 | 0.1584 | 0.0254 |
| 3.8 | Y649 | 1.0078 | 0.0056 | 0.2684 | 0.0607 | 0.0351 |
| 3.8 | Y655 | 1.0509 | 0.0056 | 0.2336 | 0.1412 | 0.0208 |
| 3.8 | Y682 | 1.0553 | 0.0051 | 0.1422 | 0.0416 | 0.0243 |
| 6.0 | Y683 | 1.3830 | 0.7839 | 1.1874 | 1.1904 | 1.0768 |
| 6.0 | Y743 | 1.3403 | 0.0215 | 0.8956 | 0.9293 | 0.4594 |
| 6.0 | Y1061 | 1.2600 | 0.9029 | 0.9354 | 1.0136 | 0.9524 |
| 6.0 | Y1062 | 1.4326 | 0.0280 | 0.9018 | 1.0266 | 0.5998 |
| 6.7 | Y1111 | 0.6878 | 0.0185 | 0.4072 | 0.3151 | 0.2538 |
| 6.7 | Y1122 | 0.7822 | 0.0302 | 0.2357 | 0.2756 | 0.1340 |
| 3.8 | Y1129 | 1.0623 | 0.0130 | -0.0004 | 0.0002 | 0.0001 |
| 3.8 | Y1136 | 1.1139 | 0.0012 | 0.3549 | 0.2995 | 0.1360 |
| 6.0 | Y1139 | 1.2612 | 0.0032 | 0.2320 | 0.0989 | 0.1892 |
| 6.0 | Y1242 | 1.3372 | 0.3735 | 0.6548 | 0.6527 | 0.4144 |
| 4.1 | Y1584 | 1.2624 | 0.0251 | 0.9367 | 0.7104 | 0.5233 |
| 4.1 | Y1613 | 1.3350 | 0.0204 | 0.8453 | 0.7542 | 0.5701 |
| 4.1 | Y1624 | 1.4281 | 0.0166 | 0.6530 | 0.5006 | 0.4334 |
| 4.1 | Y1671 | 1.3245 | 0.0048 | 0.0289 | 0.0389 | 0.1448 |
| 6.0 | Y1674 | 1.2759 | 0.0051 | 0.0837 | 0.3073 | 0.2507 |
| 6.7 | Y1703 | 1.4051 | 0.0207 | 0.1193 | 0.3087 | 0.2599 |
| 6.7 | Y1705 | 1.3702 | 0.0146 | 0.0422 | 0.2464 | 0.5607 |
| 6.0 | Y1728 | 1.5393 | 1.4013 | 1.5548 | 1.6176 | 1.8749 |

**Figure S9. $OD_{600}$ of cultures of the 24 strains studied across different experimental settings and cultivation periods.**

The source data are presented in Tables S5-S8. The yellow background indicates precultivation on PDA and pink on YM-10 agar.

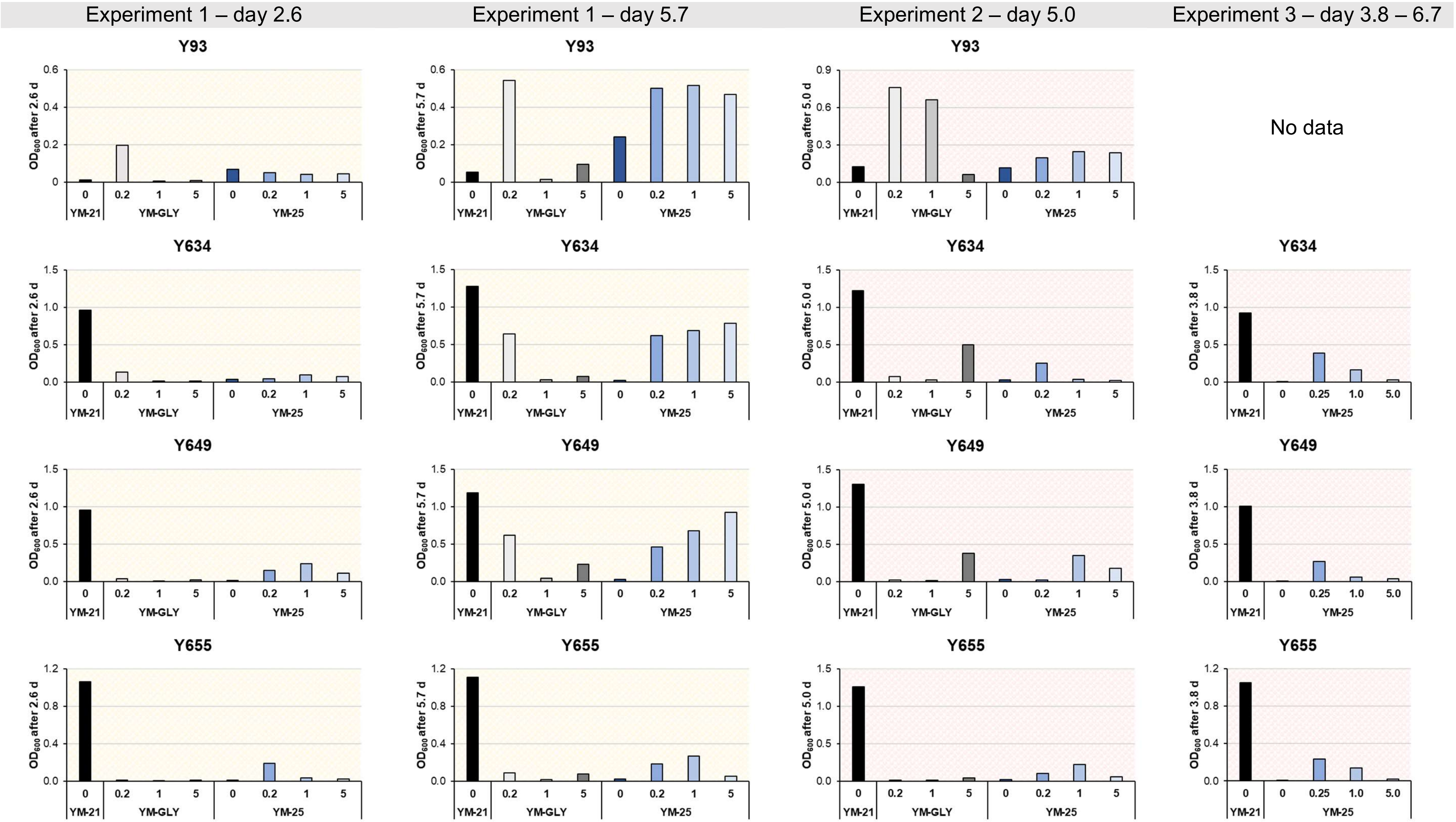

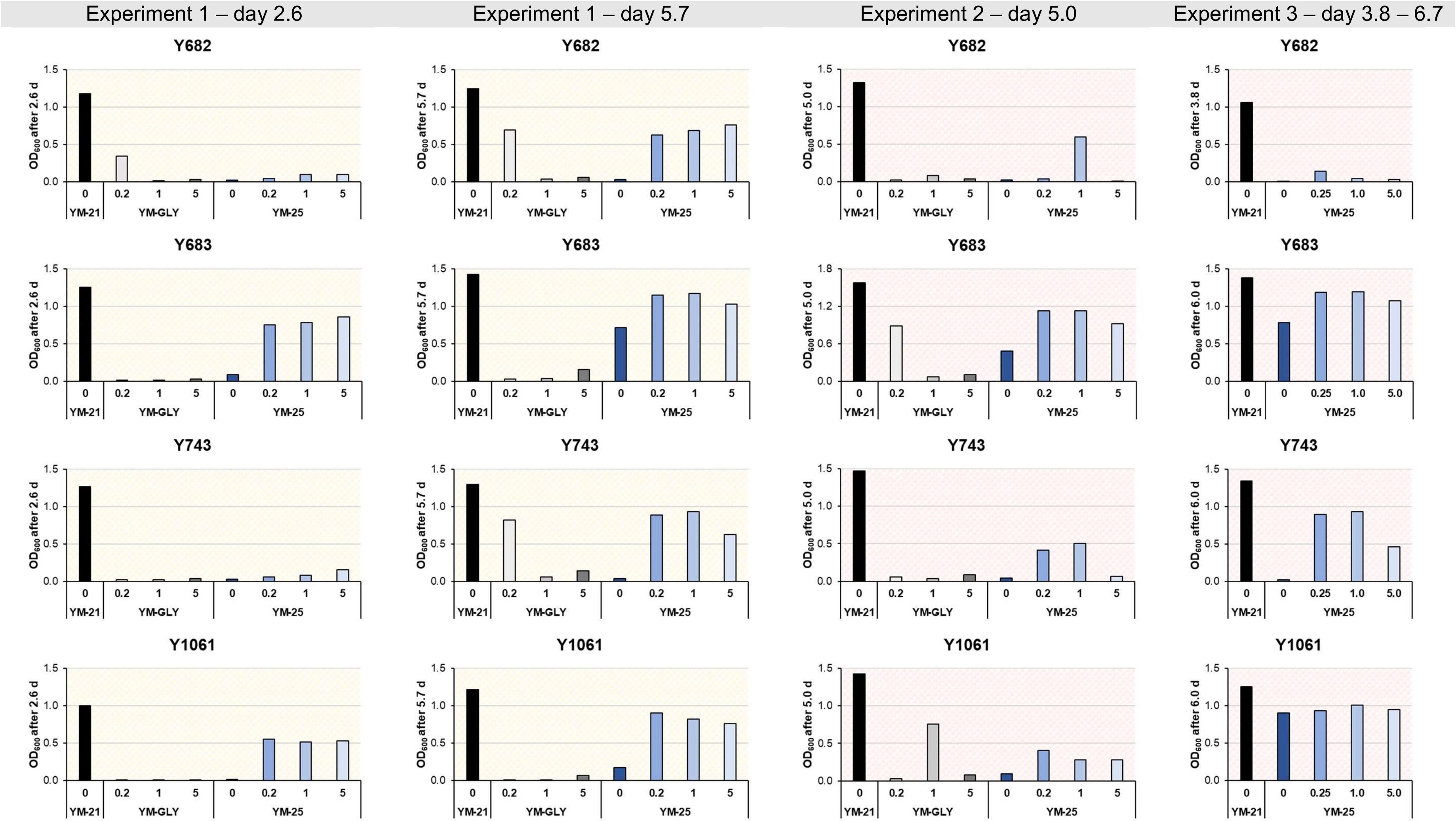
Experiment 1 – day 2.6
Experiment 1 – day 5.7
Experiment 2 – day 5.0
Experiment 3 – day 3.8 – 6.7
Y682
Y683
Y743
Y1061
OD600 after 2.6 d
OD600 after 5.7 d
OD600 after 5.0 d
OD600 after 3.8 d
OD600 after 6.0 d
0
0.2
1
5
0.25
1.0
5.0
YM-21
YM-GLY
YM-25

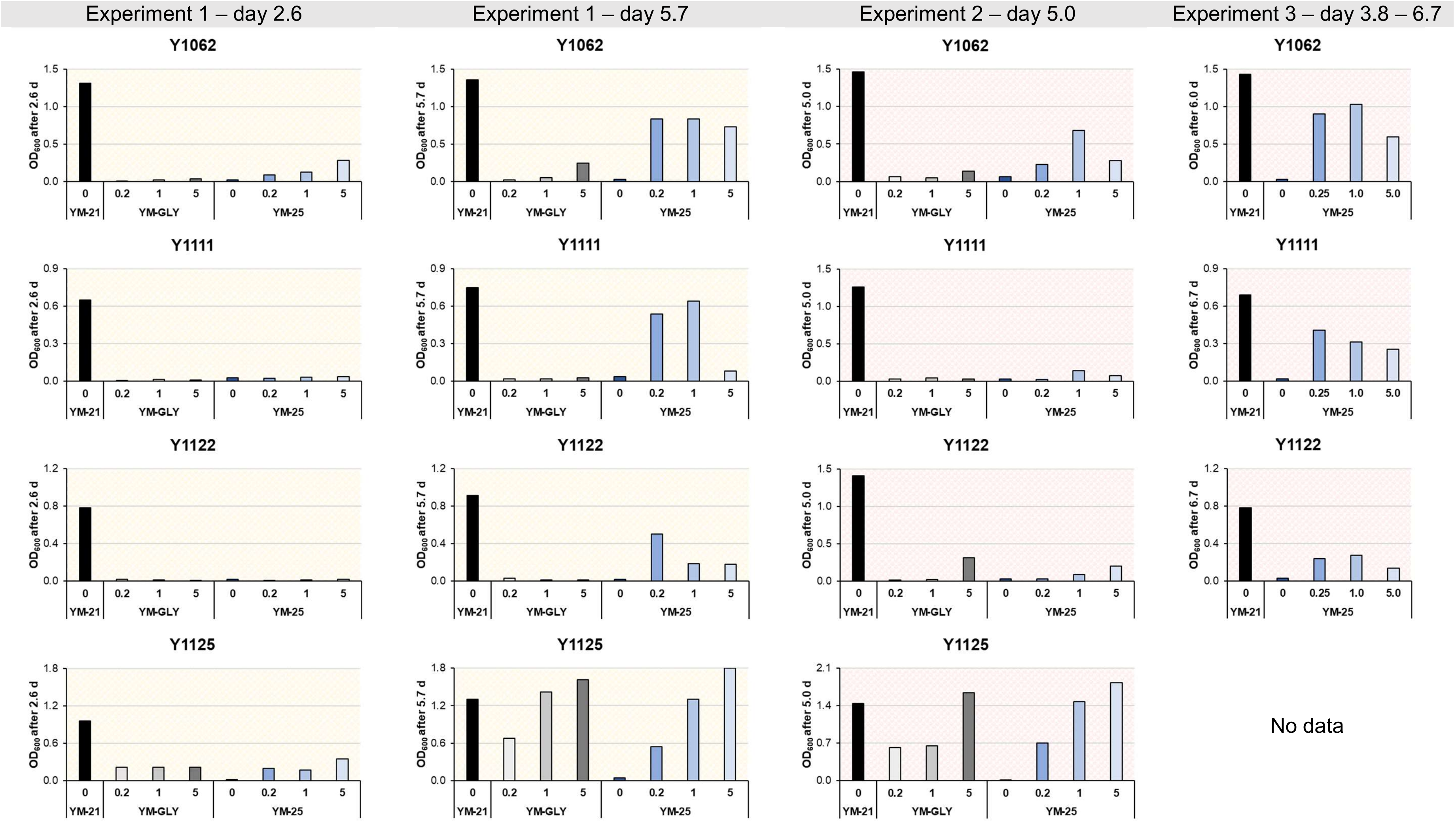
Experiment 1 – day 2.6
Experiment 1 – day 5.7
Experiment 2 – day 5.0
Experiment 3 – day 3.8 – 6.7
Y1062
Y1111
Y1122
Y1125
OD600 after 2.6 d
OD600 after 5.7 d
OD600 after 5.0 d
OD600 after 6.0 d
OD600 after 6.7 d
0
0.2
1
5
0.25
1.0
5.0
YM-21
YM-GLY
YM-25
No data

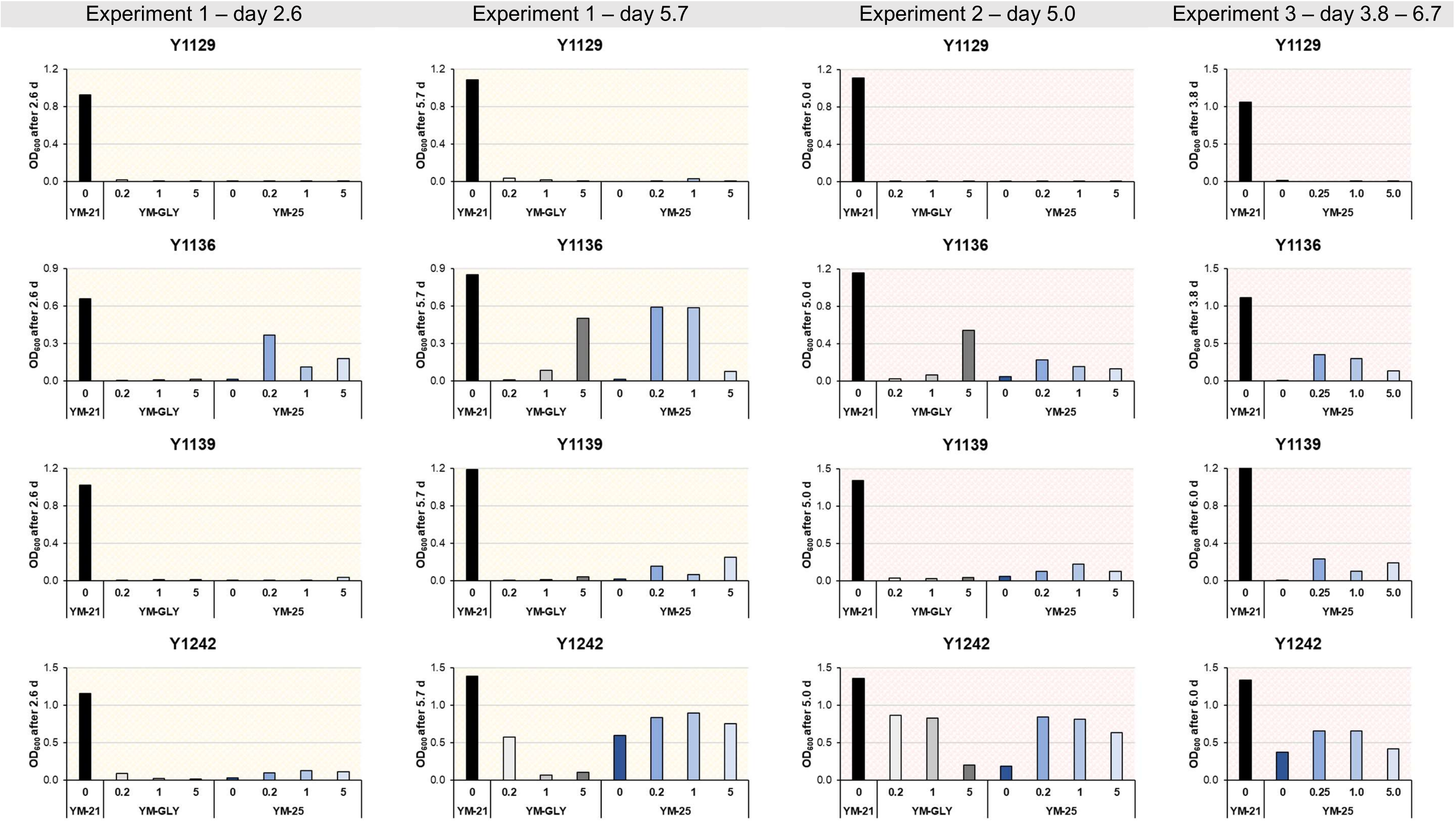
Experiment 1 – day 2.6
Experiment 1 – day 5.7
Experiment 2 – day 5.0
Experiment 3 – day 3.8 – 6.7
Y1129
Y1136
Y1139
Y1242
OD600 after 2.6 d
OD600 after 5.7 d
OD600 after 5.0 d
OD600 after 3.8 d
OD600 after 6.0 d
0
0.2
1
5
0.25
1.0
5.0
YM-21
YM-GLY
YM-25

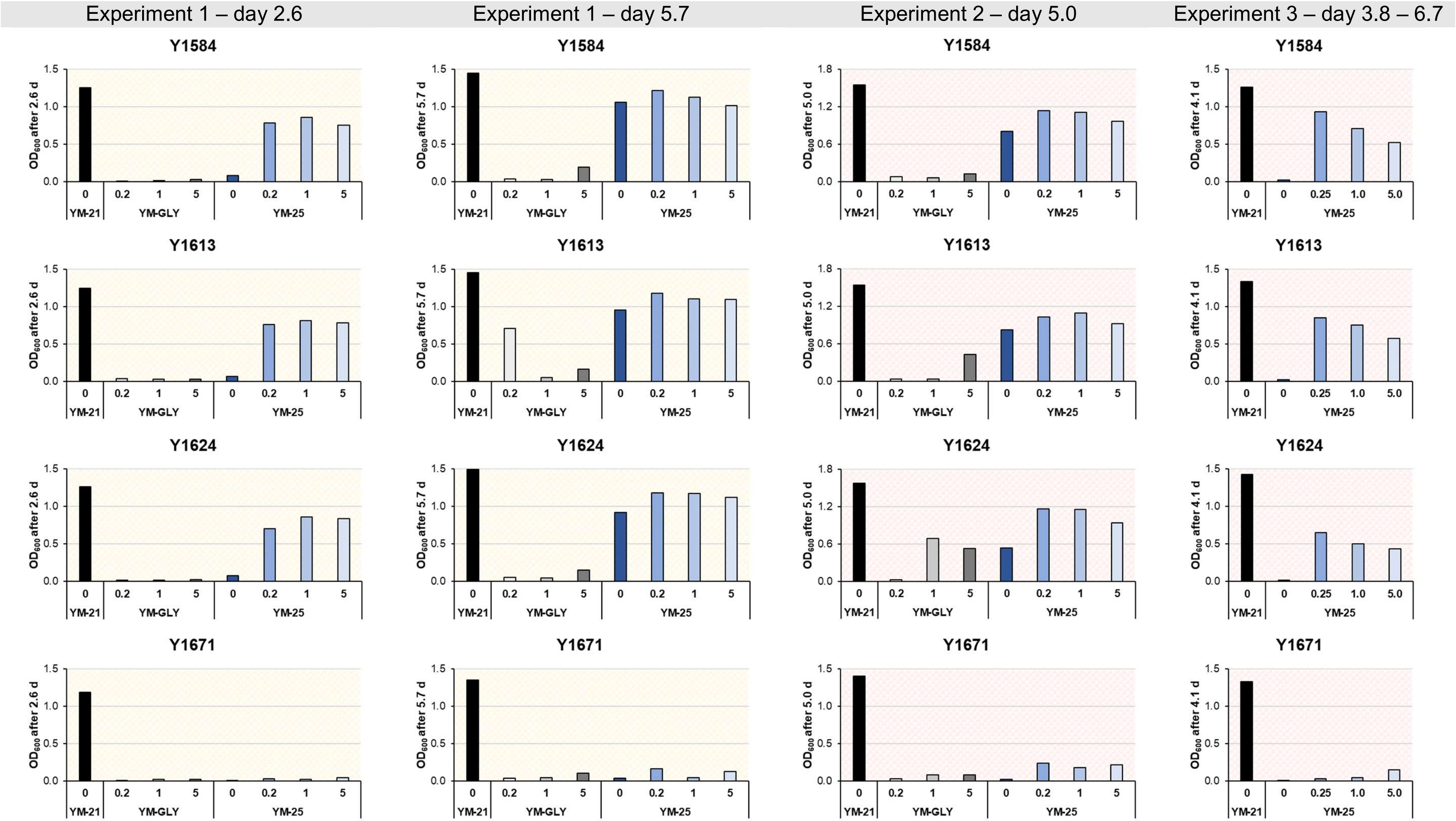
Experiment 1 – day 2.6
Experiment 1 – day 5.7
Experiment 2 – day 5.0
Experiment 3 – day 3.8 – 6.7
Y1584
Y1613
Y1624
Y1671
OD600 after 2.6 d
OD600 after 5.7 d
OD600 after 5.0 d
OD600 after 4.1 d
YM-21
YM-GLY
YM-25
0
0.2
1
5
0.25
1.0
5.0

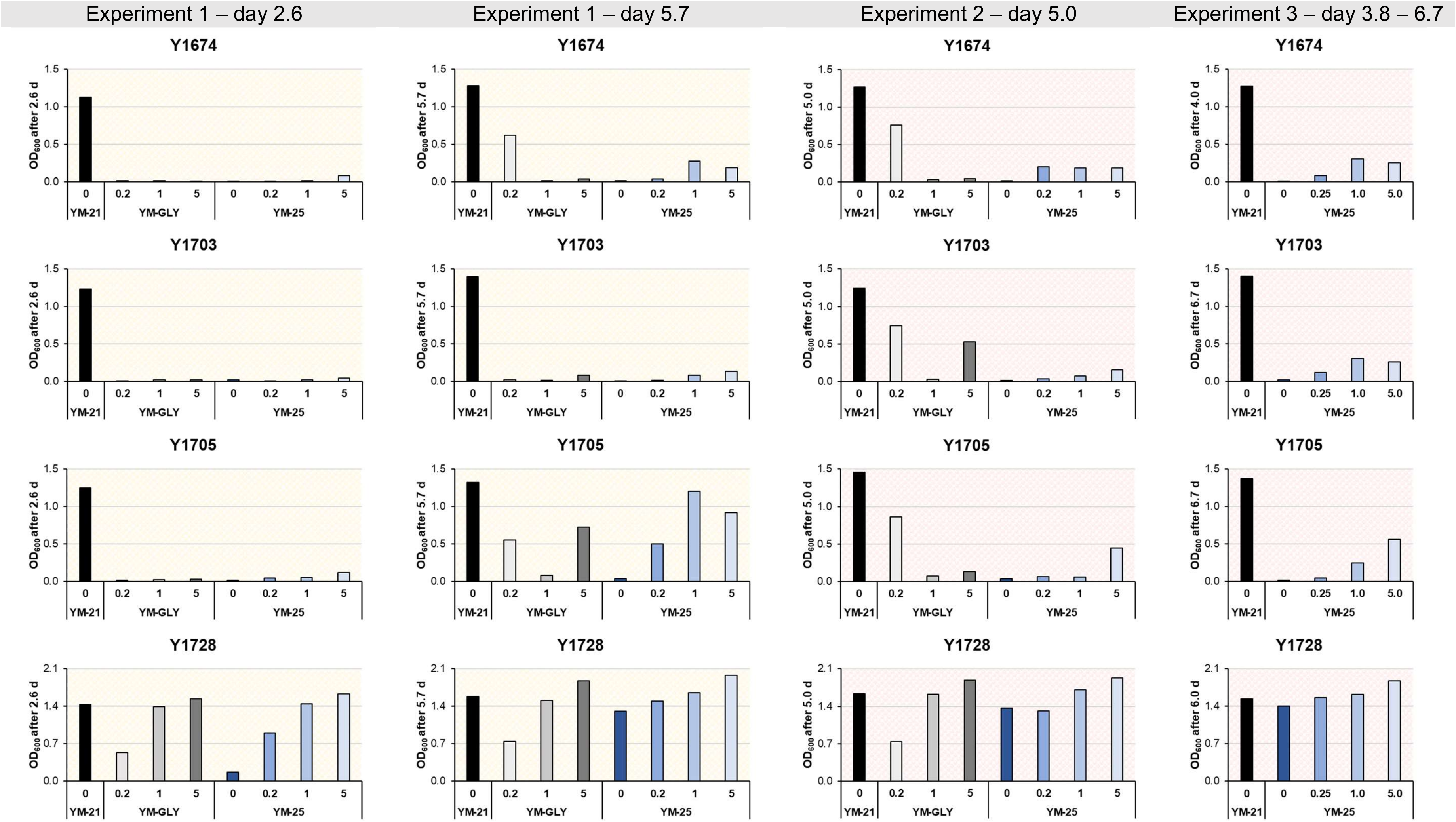
Experiment 1 – day 2.6
Experiment 1 – day 5.7
Experiment 2 – day 5.0
Experiment 3 – day 3.8 – 6.7
Y1674
Y1703
Y1705
Y1728
OD600 after 2.6 d
OD600 after 5.7 d
OD600 after 5.0 d
OD600 after 4.0 d
OD600 after 6.7 d
OD600 after 6.0 d
YM-21
YM-GLY
YM-25
0
0.2
1
5
0.25
1.0
5.0

**Table S9. $OD_{600}$ percentage of each strain in YM-25 relative to that in YM-21.**

The calculation is based on the values listed in Tables S6 and S7. The blue background indicates strains that are considered able to grow relatively well on lactate.

| Group | Strain | Precultivated on PDA (Exp. 1 – day 5.7) | Precultivated on YM-10 agar (Exp. 2 – day 5.0) |
|---|---|---|---|
| *S. cerevisiae* | Y93 | 435.71 | 94.45 |
| | Y634 | 2.07 | 1.93 |
| | Y649 | 2.33 | 2.02 |
| | Y655 | 2.41 | 2.01 |
| | Y682 | 1.92 | 1.87 |
| | Y683 | 50.38 | 30.72 |
| | Y743 | 2.90 | 2.60 |
| | Y1061 | 14.19 | 6.75 |
| | Y1062 | 2.08 | 4.10 |
| | Y1111 | 4.86 | 2.25 |
| | Y1122 | 2.17 | 2.01 |
| | Y1129 | 0.00 | 0.20 |
| | Y1136 | 1.69 | 3.75 |
| | Y1139 | 1.33 | 4.08 |
| | Y1242 | 43.12 | 13.90 |
| | Y1584 | 73.30 | 52.18 |
| | Y1613 | 65.66 | 53.55 |
| | Y1624 | 61.68 | 34.18 |
| | Y1671 | 2.37 | 1.24 |
| | Y1674 | 1.24 | 0.86 |
| | Y1703 | 0.47 | 1.09 |
| | Y1705 | 3.10 | 2.46 |
| Non-*S. cerevisiae* | Y1125 | 3.29 | 0.62 |
| | Y1728 | 82.45 | 82.89 |

**Table S10. Time-course $OD_{600}$ evolution of the seven relatively-well lactate growers in YM-21 and YM-25.**

Compiled from the data in Tables S5 and S6. The $OD_{600}$ at day 0, i.e., the start of the cultivation, was not measured but assumed to be 0.

| Group | Strain | Precultivated on PDA (Exp. 1 – day 5.7) | | | | | |
|---|---|---|---|---|---|---|---|
| | | YM-21 | | | YM-25 | | |
| | | 0 d | 2.6 d | 5.7 d | 0 d | 2.6 d | 5.7 d |
| *S. cerevisiae* | Y683 | 0 | 1.2521 | 1.4209 | 0 | 0.0868 | 0.7158 |
| | Y1061 | 0 | 1.0066 | 1.2159 | 0 | 0.0207 | 0.1725 |
| | Y1242 | 0 | 1.1541 | 1.3842 | 0 | 0.0256 | 0.5969 |
| | Y1584 | 0 | 1.2553 | 1.4521 | 0 | 0.0837 | 1.0644 |
| | Y1613 | 0 | 1.243 | 1.4535 | 0 | 0.0671 | 0.9543 |
| | Y1624 | 0 | 1.2666 | 1.4914 | 0 | 0.0738 | 0.9199 |
| Non *S. cerevisiae* | Y1728 | 0 | 1.4336 | 1.5849 | 0 | 0.1752 | 1.3067 |

#### Figure S10. Time-course $OD_{600}$ evolution of the four relatively-well lactate growers in YM-21 and YM-25.

The yellow background indicates precultivation on PDA. The source data are presented in Table S10.

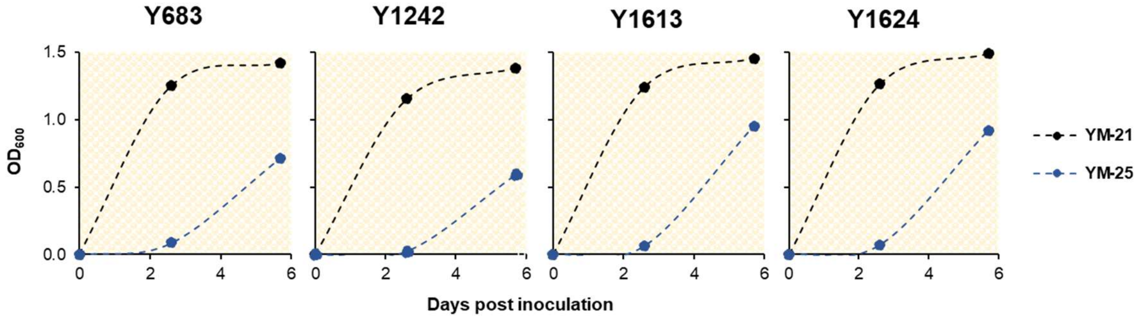

#### Table S11. $OD_{600}$ percentage of cultures grown in YM-GLY relative to that in YM-21.

The subcripted number following "GLY" indicates the concentration (% w/v) of glycerol in the respective medium. The calculation is based on the values listed in Tables S6 and S7.

| Group | Strain | Precultivated on YM-10 agar (Exp. 1 – day 5.7) | | | Precultivated on YM-10 agar (Exp. 2 – day 5.0) | | |
|---|---|---|---|---|---|---|---|
| | | YM-$GLY_{0.2}$ | YM-$GLY_{1.0}$ | YM-$GLY_{5.0}$ | YM-$GLY_{0.2}$ | YM-$GLY_{1.0}$ | YM-$GLY_{5.0}$ |
| *S. cerevisiae* | Y93 | 972.14 | 28.21 | 174.46 | 621.39 | 538.45 | 50.94 |
| | Y634 | 50.66 | 2.64 | 6.18 | 5.70 | 2.53 | 40.85 |
| | Y649 | 51.86 | 3.60 | 19.71 | 1.51 | 1.07 | 29.14 |
| | Y655 | 8.01 | 1.88 | 7.01 | 1.10 | 1.37 | 3.46 |
| | Y682 | 55.51 | 2.99 | 4.42 | 1.63 | 6.23 | 3.11 |
| | Y683 | 2.23 | 2.41 | 11.21 | 56.43 | 4.68 | 6.73 |
| | Y743 | 62.93 | 4.16 | 10.60 | 3.72 | 2.50 | 5.76 |
| | Y1061 | 0.63 | 0.91 | 5.53 | 1.97 | 52.88 | 5.70 |
| | Y1062 | 1.76 | 3.85 | 18.16 | 4.52 | 3.55 | 9.50 |
| | Y1111 | 2.58 | 2.17 | 3.69 | 2.24 | 3.52 | 1.94 |
| | Y1122 | 3.67 | 1.37 | 1.31 | 1.10 | 1.61 | 22.11 |
| | Y1129 | 3.25 | 1.54 | 0.27 | 0.46 | 0.19 | 0.29 |
| | Y1136 | 1.39 | 10.04 | 58.95 | 1.79 | 5.33 | 46.74 |
| | Y1139 | 0.42 | 0.87 | 3.20 | 2.46 | 2.21 | 3.27 |
| | Y1242 | 41.37 | 4.61 | 7.66 | 63.72 | 61.03 | 15.10 |
| | Y1584 | 2.79 | 2.19 | 13.77 | 5.09 | 4.29 | 8.33 |
| | Y1613 | 48.80 | 3.43 | 11.36 | 2.20 | 2.59 | 28.08 |
| | Y1624 | 3.92 | 3.37 | 10.21 | 1.60 | 43.52 | 33.28 |
| | Y1671 | 2.61 | 3.41 | 7.74 | 1.79 | 5.80 | 5.89 |
| | Y1674 | 48.04 | 1.21 | 2.64 | 59.89 | 2.24 | 3.35 |
| | Y1703 | 1.31 | 1.17 | 5.89 | 59.87 | 2.41 | 41.99 |
| | Y1705 | 42.21 | 6.22 | 55.01 | 58.85 | 4.71 | 9.04 |
| Non-*S. cerevisiae* | Y1125 | 51.53 | 108.87 | 123.50 | 42.54 | 45.26 | 113.57 |
| | Y1728 | 46.81 | 95.32 | 118.24 | 44.66 | 98.90 | 115.13 |

##### Table S12. Ratio of $OD_{600}$ percentage of cultures in YM-GLY / YM-21 relative to the maximum value.

The apparently-preferred glycerol concentration (i.e., the one resulting in the highest $OD_{600}$ value) is highlighted with yellow or pink, depending on the precultivation medium being PDA or YM-10. The calculation is based on the values listed in Table S11.

| Group | Strain | Precultivated on YM-10 agar (Exp. 1 – day 5.7) | | | | Precultivated on YM-10 agar (Exp. 2 – day 5.0) | | | |
|---|---|---|---|---|---|---|---|---|---|
| | | Max. % $OD_{600}$ in YM-GLY / YM-21 | YM-$GLY_{0.2}$ | YM-$GLY_{1.0}$ | YM-$GLY_{5.0}$ | Max. % $OD_{600}$ in YM-GLY / YM-21 | YM-$GLY_{0.2}$ | YM-$GLY_{1.0}$ | YM-$GLY_{5.0}$ |
| *S. cerevisiae* | Y93 | 972.14 | 1.0 | 0.0 | 0.2 | 621.4 | 1.0 | 0.9 | 0.1 |
| | Y634 | 50.66 | 1.0 | 0.1 | 0.1 | 40.9 | 0.1 | 0.1 | 1.0 |
| | Y649 | 51.86 | 1.0 | 0.1 | 0.4 | 29.1 | 0.1 | 0.0 | 1.0 |
| | Y655 | 8.01 | 1.0 | 0.2 | 0.9 | 3.5 | 0.3 | 0.4 | 1.0 |
| | Y682 | 55.51 | 1.0 | 0.1 | 0.1 | 6.2 | 0.3 | 1.0 | 0.5 |
| | Y683 | 11.21 | 0.2 | 0.2 | 1.0 | 56.4 | 1.0 | 0.1 | 0.1 |
| | Y743 | 62.93 | 1.0 | 0.1 | 0.2 | 5.8 | 0.6 | 0.4 | 1.0 |
| | Y1061 | 5.53 | 0.1 | 0.2 | 1.0 | 52.9 | 0.0 | 1.0 | 0.1 |
| | Y1062 | 18.16 | 0.1 | 0.2 | 1.0 | 9.5 | 0.5 | 0.4 | 1.0 |
| | Y1111 | 3.69 | 0.7 | 0.6 | 1.0 | 3.5 | 0.6 | 1.0 | 0.6 |
| | Y1122 | 3.67 | 1.0 | 0.4 | 0.4 | 22.1 | 0.1 | 0.1 | 1.0 |
| | Y1125 | 123.50 | 0.4 | 0.9 | 1.0 | 113.6 | 0.4 | 0.4 | 1.0 |
| | Y1129 | 3.25 | 1.0 | 0.5 | 0.1 | 0.5 | 1.0 | 0.4 | 0.6 |
| | Y1136 | 58.95 | 0.0 | 0.2 | 1.0 | 46.7 | 0.0 | 0.1 | 1.0 |
| | Y1139 | 3.20 | 0.1 | 0.3 | 1.0 | 3.3 | 0.8 | 0.7 | 1.0 |
| | Y1242 | 41.37 | 1.0 | 0.1 | 0.2 | 63.7 | 1.0 | 1.0 | 0.2 |
| | Y1584 | 13.77 | 0.2 | 0.2 | 1.0 | 8.3 | 0.6 | 0.5 | 1.0 |
| | Y1613 | 48.80 | 1.0 | 0.1 | 0.2 | 28.1 | 0.1 | 0.1 | 1.0 |
| | Y1624 | 10.21 | 0.4 | 0.3 | 1.0 | 43.5 | 0.0 | 1.0 | 0.8 |
| | Y1671 | 7.74 | 0.3 | 0.4 | 1.0 | 5.9 | 0.3 | 1.0 | 1.0 |
| | Y1674 | 48.04 | 1.0 | 0.0 | 0.1 | 59.9 | 1.0 | 0.0 | 0.1 |
| | Y1703 | 5.89 | 0.2 | 0.2 | 1.0 | 59.9 | 1.0 | 0.0 | 0.7 |
| | Y1705 | 55.01 | 0.8 | 0.1 | 1.0 | 58.9 | 1.0 | 0.1 | 0.2 |
| Non *S. cerevisiae* | Y1125 | 123.50 | 0.4 | 0.9 | 1.0 | 113.6 | 0.4 | 0.4 | 1.0 |
| | Y1728 | 118.24 | 0.4 | 0.8 | 1.0 | 115.1 | 0.4 | 0.9 | 1.0 |

**Table S13. Analysis of glycerol concentration preference agreement between cultures precultivated on PDA versus YM-10 agar.**

The determination is based on the values listed in Table S12.

| Group | Strain | Precultivated on PDA (Exp. 1 – day 5.7) | | | Precultivated on YM-10 agar (Exp. 2 – day 5.0) | | | Agreement assesment based on Exp. 1 and Exp. 2 | | | |
|---|---|---|---|---|---|---|---|---|---|---|---|
| | | YM-GLY$_{0.2}$ | YM-GLY$_{1.0}$ | YM-GLY$_{5.0}$ | YM-GLY$_{0.2}$ | YM-GLY$_{1.0}$ | YM-GLY$_{5.0}$ | YM-GLY$_{0.2}$ | YM-GLY$_{1.0}$ | YM-GLY$_{5.0}$ | No preference |
| *S. cerevisiae* | Y93 | 1 | | | 1 | | | 1 | | | |
| | Y634 | 1 | | | | | 1 | | | | 1 |
| | Y649 | 1 | | | | | 1 | | | | 1 |
| | Y655 | 1 | | | | | 1 | | | | 1 |
| | Y682 | 1 | | | | 1 | | | | | 1 |
| | Y683 | | | 1 | 1 | | | | | | 1 |
| | Y743 | 1 | | | | | 1 | | | | 1 |
| | Y1061 | | | 1 | | 1 | | | | | 1 |
| | Y1062 | | | 1 | | | 1 | | | 1 | |
| | Y1111 | | | 1 | | 1 | | | | | 1 |
| | Y1122 | 1 | | | | | 1 | | | | 1 |
| | Y1129 | 1 | | | 1 | | | 1 | | | |
| | Y1136 | | | 1 | | | 1 | | | 1 | |
| | Y1139 | | | 1 | | | 1 | | | 1 | |
| | Y1242 | 1 | | | 1 | | | 1 | | | |
| | Y1584 | | | 1 | | | 1 | | | 1 | |
| | Y1613 | 1 | | | | | 1 | | | | 1 |
| | Y1624 | | | 1 | | 1 | | | | | 1 |
| | Y1671 | | | 1 | | | 1 | | | 1 | |
| | Y1674 | 1 | | | 1 | | | 1 | | | |
| | Y1703 | | | 1 | 1 | | | | | | 1 |
| | Y1705 | | | 1 | 1 | | | | | | 1 |
| | TOTAL (*n* = 22) | 11 (50%) | - | 11 (50%) | 7 (32%) | 4 (18%) | 11 (50%) | 4 (18%) | - | 5 (23%) | 13 (59%) |
| Non *S. cerevisiae* | Y1125 | | | 1 | | | 1 | | | 1 | |
| | Y1728 | | | 1 | | | 1 | | | 1 | |
| | TOTAL (*n* = 2) | - | - | 2 (100%) | - | - | 2 (100%) | - | - | 2 (100%) | - |

#### Figure S11. Bar charts indicating YM-GLY preference of individual strains.

The yellow background indicates precultivation on PDA and pink on YM-10 agar. The source data are presented in Table S12.

*Both PDA- and YM-10-agar-precultivated clones prefer YM-GLY$_{0.2}$*

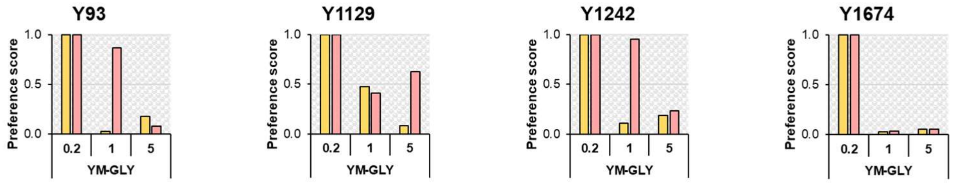

*Both PDA- and YM-10-agar-precultivated clones prefer YM-GLY$_{5.0}$*

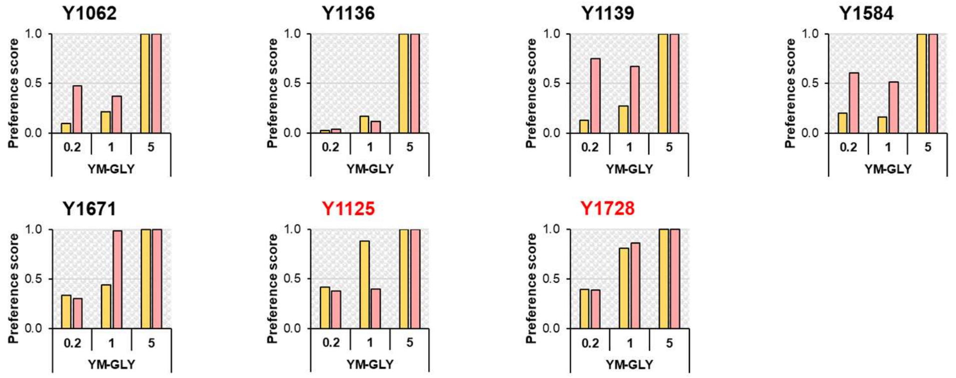

*PDA- and YM-10-agar-precultivated clones displayed different preferences for YM-GLY*

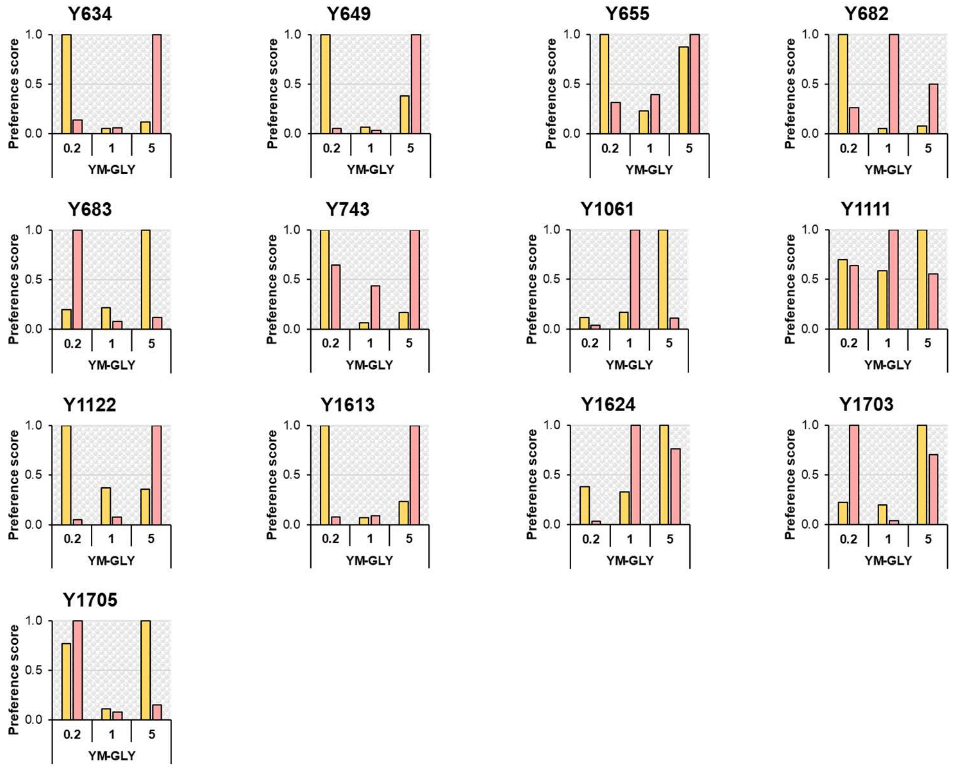

**Figure S12. $OD_{600}$ percentage of cultures grown in YM-GLY relative to that in YM-21 as a function of initial glycerol concentration.**

The yellow background indicates precultivation on PDA and pink on YM-10 agar. The source data are presented in Table S11.

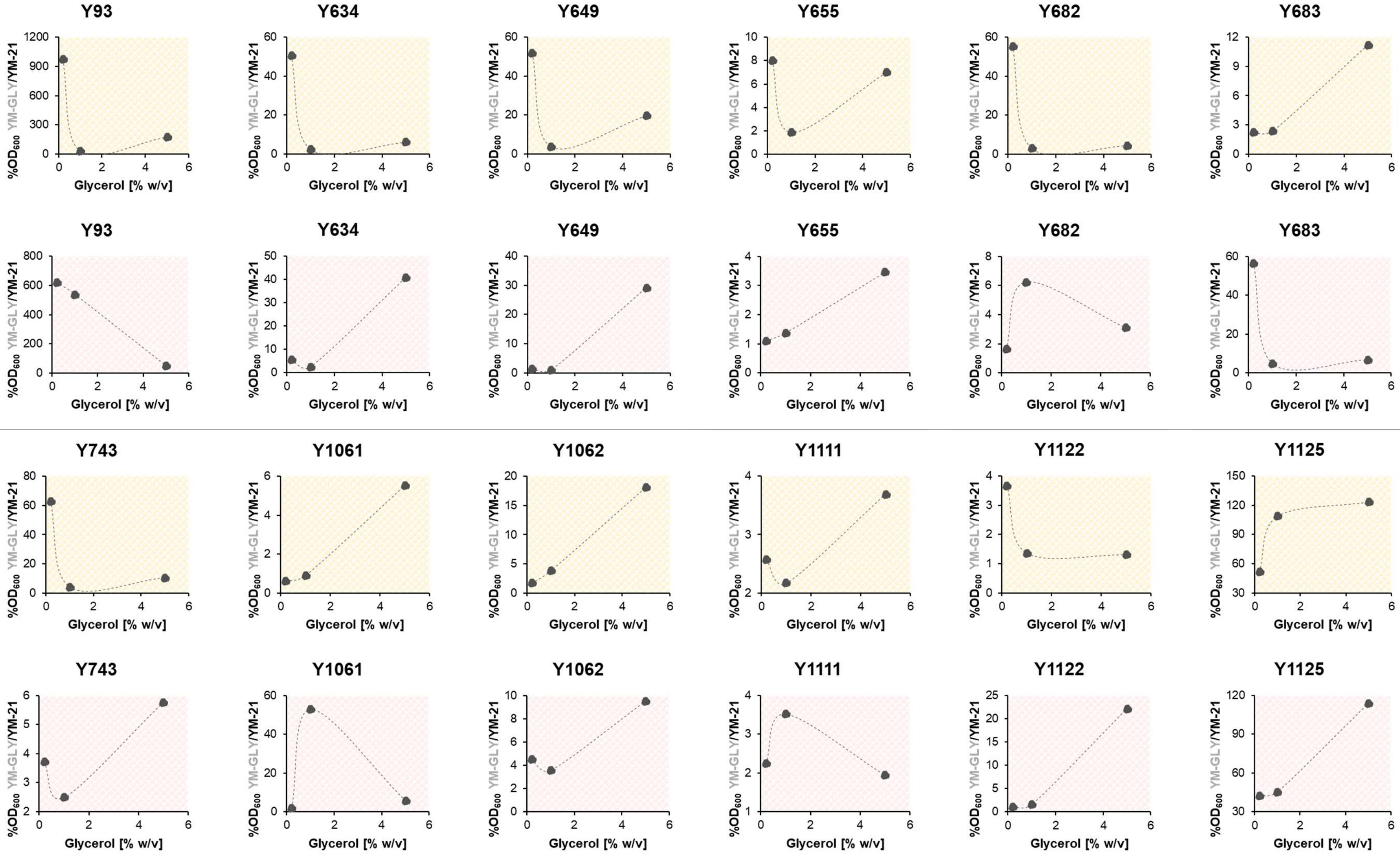

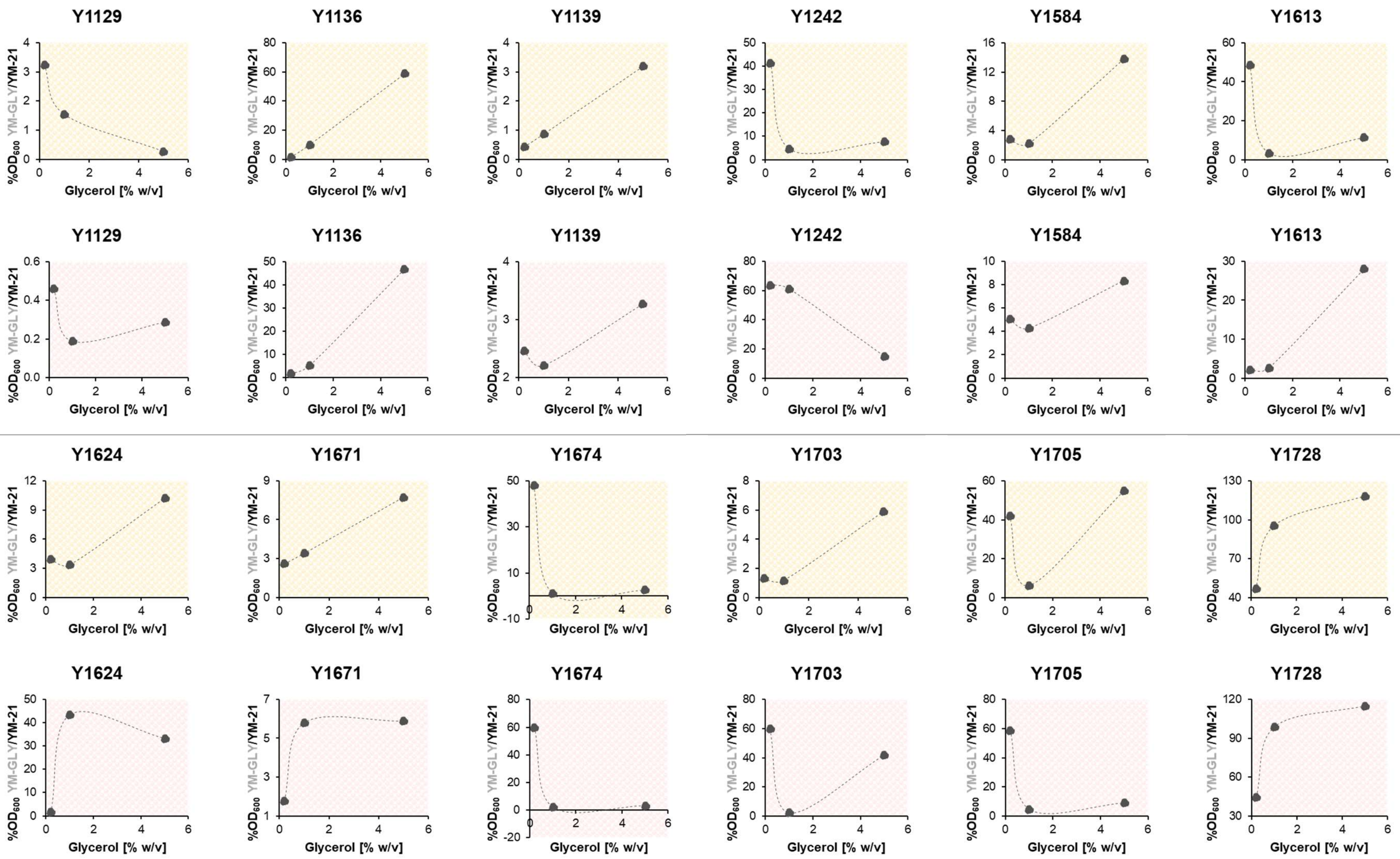
Y1129
Y1136
Y1139
Y1242
Y1584
Y1613
Y1129
Y1136
Y1139
Y1242
Y1584
Y1613
Y1624
Y1671
Y1674
Y1703
Y1705
Y1728
Y1624
Y1671
Y1674
Y1703
Y1705
Y1728
%OD600 YM-GLY/YM-21
Glycerol [% w/v]

#### Table S14. $OD_{600}$ percentage of cultures grown in YM-25 + GLY relative to that in YM-21.

The calculation is based on the values listed in Tables S6 and S7 .

| Group | Subgroup | Strain | Precultivated on PDA (Exp. 1 – day 5.7) | | | | Precultivated on YM-10 agar (Exp. 2 – day 5.0) | | | |
|---|---|---|---|---|---|---|---|---|---|---|
| | | | YM-25 + $GLY_{0.2}$ | YM-25 + $GLY_{1.0}$ | YM-25 + $GLY_{5.0}$ | Max. | YM-25 + $GLY_{0.2}$ | YM-25 + $GLY_{1.0}$ | YM-25 + $GLY_{5.0}$ | Max. |
| *S. cerevisiae* | **Outlier** | Y93 | 897.86 | 925.54 | 840.18 | 925.54 | 158.20 | 200.90 | 192.16 | 200.90 |
| | **Lactate – (15)** | Y1129 | 0.06 | 2.81 | 0.14 | 2.81 | 0.36 | 0.09 | 0.50 | 0.50 |
| | | Y1703 | 1.10 | 5.81 | 9.35 | 9.35 | 2.80 | 5.60 | 12.20 | 12.20 |
| | | Y1671 | 12.27 | 2.96 | 9.27 | 12.27 | 17.09 | 12.49 | 15.14 | 17.09 |
| | | Y1139 | 13.25 | 5.37 | 20.90 | 20.90 | 9.19 | 16.47 | 9.19 | 16.47 |
| | | Y1674 | 2.96 | 21.48 | 14.30 | 21.48 | 15.65 | 14.84 | 14.54 | 15.65 |
| | | Y655 | 17.00 | 24.58 | 4.80 | 24.58 | 8.61 | 17.71 | 4.73 | 17.71 |
| | **Lactate – (70)** | Y1122 | 54.99 | 20.67 | 19.95 | 54.99 | 1.69 | 6.08 | 14.39 | 14.39 |
| | | Y682 | 50.12 | 55.11 | 60.92 | 60.92 | 2.89 | 44.99 | 0.39 | 44.99 |
| | | Y1062 | 61.47 | 61.20 | 53.93 | 61.47 | 15.66 | 46.25 | 18.94 | 46.25 |
| | | Y634 | 48.92 | 53.98 | 61.80 | 61.80 | 20.81 | 2.99 | 1.58 | 20.81 |
| | | Y1136 | 69.26 | 68.87 | 8.87 | 69.26 | 19.44 | 13.19 | 11.41 | 19.44 |
| | | Y743 | 68.25 | 71.50 | 47.91 | 71.50 | 28.12 | 34.12 | 4.54 | 34.12 |
| | | Y649 | 38.89 | 57.14 | 77.58 | 77.58 | 1.54 | 27.03 | 13.54 | 27.03 |
| | | Y1111 | 71.72 | 85.94 | 10.75 | 85.94 | 1.79 | 10.89 | 5.64 | 10.89 |
| | | Y1705 | 37.76 | 91.19 | 69.48 | 91.19 | 4.41 | 4.06 | 30.29 | 30.29 |
| | **Lactate +** | Y1242 | 60.57 | 64.35 | 54.26 | 64.35 | 62.15 | 59.74 | 46.99 | 62.15 |
| | | Y1061 | 74.79 | 67.88 | 62.61 | 74.79 | 28.23 | 19.52 | 19.73 | 28.23 |
| | | Y1624 | 79.31 | 78.81 | 75.41 | 79.31 | 74.04 | 73.28 | 59.68 | 74.04 |
| | | Y1613 | 80.82 | 75.71 | 75.08 | 80.82 | 66.99 | 70.86 | 60.09 | 70.86 |
| | | Y683 | 80.98 | 82.65 | 72.53 | 82.65 | 71.67 | 71.43 | 58.64 | 71.67 |
| | | Y1584 | 83.73 | 77.76 | 70.22 | 83.73 | 73.59 | 71.63 | 62.73 | 73.59 |
| Non *S. cerevisiae* | **Lactate –** | Y1125 | 41.65 | 99.54 | 138.25 | 138.25 | 48.97 | 101.94 | 126.88 | 126.88 |
| | **Lactate +** | Y1728 | 94.66 | 104.62 | 124.57 | 124.57 | 79.72 | 104.33 | 117.64 | 117.64 |

#### Table S15. Datasets used to calculate statistical significance of differences between groups of cultures.

$GLY_{max}$: the concentration of glycerol inside the respective YM-25 + GLY that resulted in the highest $OD_{600}$. *n* : the number of the group's member, i.e., the group's sample size. The datasets compile the values listed in Tables S9 and S14.

| Row | *S. cerevisiae* group | Strain | Growth in YM-25 | | Growth in YM-25 + GLY | | | |
|---|---|---|---|---|---|---|---|---|
| | | | Precultivated on PDA (Exp. 1 – day 5.7) | Precultivated on YM-10 agar (Exp. 2 – day 5.0) | Precultivated on PDA (Exp. 1 – day 5.7) | $GLY_{max}$ | Precultivated on YM-10 agar (Exp. 2 – day 5.0) | $GLY_{max}$ |
| | | | Max. %$OD_{600}$ in YM-25 / YM-21 | Max. %$OD_{600}$ in YM-25 / YM-21 | %$OD_{600}$ in [YM-25 + $GLY_{max}$] / YM-21 | [% w/v] | Max. %$OD_{600}$ in [YM-25 + $GLY_{max}$] / YM-21 | [% w/v] |
| | | | ***1*** | ***2*** | ***3*** | ***3**** | ***4*** | ***4**** |
| ***a*** | **Lactate – (15)** (*n* = 6) | Y1129 | 0.00 | 0.20 | 2.81 | 1.0 | 0.50 | 5.0 |
| | | Y1703 | 0.47 | 1.09 | 9.35 | 5.0 | 12.20 | 5.0 |
| | | Y1671 | 2.37 | 1.24 | 12.27 | 0.2 | 17.09 | 0.2 |
| | | Y1139 | 1.33 | 4.08 | 20.90 | 5.0 | 16.47 | 1.0 |
| | | Y1674 | 1.24 | 0.86 | 21.48 | 1.0 | 15.65 | 0.2 |
| | | Y655 | 2.41 | 2.01 | 24.58 | 1.0 | 17.71 | 1.0 |
| | | **Mean** ± stdev | **1.30** ± 0.98 | **1.58** ± 1.36 | **15.23** ± 8.44 | Mode: 1.0 | **13.27** ± 6.55 | Mode: 5.0 |
| ***b*** | **Lactate – (70)** (*n* = 9) | Y1122 | 2.17 | 2.01 | 54.99 | 0.2 | 14.39 | 5.0 |
| | | Y682 | 1.92 | 1.87 | 60.92 | 5.0 | 44.99 | 1.0 |
| | | Y1062 | 2.08 | 4.10 | 61.47 | 0.2 | 46.25 | 1.0 |
| | | Y634 | 2.07 | 1.93 | 61.80 | 5.0 | 20.81 | 0.2 |
| | | Y1136 | 1.69 | 3.75 | 69.26 | 0.2 | 19.44 | 0.2 |
| | | Y743 | 2.90 | 2.60 | 71.50 | 1.0 | 34.12 | 1.0 |
| | | Y649 | 2.33 | 2.02 | 77.58 | 5.0 | 27.03 | 1.0 |
| | | Y1111 | 4.86 | 2.25 | 85.94 | 1.0 | 10.89 | 1.0 |
| | | Y1705 | 3.10 | 2.46 | 91.19 | 1.0 | 30.29 | 5.0 |
| | | **Mean** ± stdev | **2.57** ± 0.97 | **2.55** ± 0.82 | **70.52** ± 12.29 | Mode: 0.2 | **27.58** ± 12.58 | Mode: 1.0 |
| ***c*** | **Lactate +** (*n* = 6) | Y1242 | 43.12 | 13.90 | 64.35 | 1.0 | 62.15 | 0.2 |
| | | Y1061 | 14.19 | 6.75 | 74.79 | 0.2 | 28.23 | 0.2 |
| | | Y1624 | 61.68 | 34.18 | 79.31 | 1.0 | 74.04 | 0.2 |
| | | Y1613 | 65.66 | 53.55 | 80.82 | 0.2 | 70.86 | 0.2 |
| | | Y683 | 50.38 | 30.72 | 82.65 | 0.2 | 71.67 | 1.0 |
| | | Y1584 | 73.30 | 52.18 | 83.73 | 0.2 | 73.59 | 0.2 |
| | | **Mean** ± stdev | **51.39** ± 21.18 | **31.88** ± 19.20 | **77.61** ± 7.21 | Mode: 0.2 | **63.42** ± 17.77 | Mode: 0.2 |

**Table S16. Statistical significance of glycerol supplementation on *S. cerevisiae* growth in YM-25.**

Statistical significance (*p*-value) is calculated using the TTEST function (two-tailed distribution, paired) in Microsoft Excel. The datasets refer to those in Table S15.

| Precultivation | Precultivated on PDA (Exp. 1 – day 5.7) | | | Precultivated on YM-10 agar (Exp. 2 – day 5.0) | | |
|---|---|---|---|---|---|---|
| *S. cerevisiae* group | Lactate – (15) | Lactate – (70) | Lactate + | Lactate – (15) | Lactate – (70) | Lactate + |
| Datasets compared | ***1a*** vs. ***3a*** | ***1b*** vs. ***3b*** | ***1c*** vs. ***3c*** | ***2a*** vs. ***4a*** | ***2b*** vs. ***4b*** | ***2c*** vs. ***4c*** |
| *p*-value | 0.007 | 0.000 | 0.017 | 0.005 | 0.000 | 0.002 |
| Symbol of *p*-value | * | *** | * | ** | *** | ** |

**Table S17. Statistical significance of precultivation medium on *S. cerevisiae* growth in YM-25 vs. YM-25 + GLY.**

$GLY_{max}$: the concentration of glycerol inside the respective YM-25 + GLY that resulted in the highest $OD_{600}$. Statistical significance (*p*-value) is calculated using the TTEST function (two-tailed distribution, paired) in Microsoft Excel. The datasets refer to those in Table S15.

| Cultivation | Growth in YM-25 | | | Growth in YM-25 + $GLY_{max}$ | | |
|---|---|---|---|---|---|---|
| *S. cerevisiae* group | Lactate – (15) | Lactate – (70) | Lactate + | Lactate – (15) | Lactate – (70) | Lactate + |
| Datasets compared | ***1a*** vs. ***2a*** | ***1b*** vs. ***2b*** | ***1c*** vs. ***2c*** | ***3a*** vs. ***4a*** | ***3b*** vs. ***4b*** | ***3c*** vs. ***4c*** |
| *p*-value | 0.637 | 0.976 | 0.002 | 0.361 | 0.000 | 0.085 |
| Symbol of *p*-value | n.s. | n.s. | ** | n.s. | *** | n.s. |

**Table S18. Statistical significance of genetic background on *S. cerevisiae* growth in YM-25 vs. YM-25 + GLY.**

Statistical significance (*p*-value) is calculated using the TTEST function (two-tailed distribution, two-sample equal variance (homoscedastic)) in Microsoft Excel. The datasets refer to those in Table S15.

| Cultivation | Growth in YM-25 | | | | Growth in YM-25 + $GLY_{max}$ | | | |
|---|---|---|---|---|---|---|---|---|
| Precultivation | Precultivated on PDA (Exp. 1 – day 5.7) | | Precultivated on YM-10 agar (Exp. 2 – day 5.0) | | Precultivated on PDA (Exp. 1 – day 5.7) | | Precultivated on YM-10 agar (Exp. 2 – day 5.0) | |
| Lactate | - (15) vs. – (70) | - (70) vs. + | - (15) vs. – (70) | - (70) vs. + | - (15) vs. – (70) | - (70) vs. + | - (15) vs. – (70) | - (70) vs. + |
| Datasets compared | ***1a*** vs. ***1b*** | ***1b*** vs. ***1c*** | ***2a*** vs. ***2b*** | ***2b*** vs. ***2c*** | ***3a*** vs. ***3b*** | ***3b*** vs. ***3c*** | ***4a*** vs. ***4b*** | ***4b*** vs. ***4c*** |
| *p*-value | 0.028 | 0.000 | 0.104 | 0.000 | 0.000 | 0.228 | 0.024 | 0.001 |
| Symbol of *p*-value | * | *** | n.s. | *** | *** | n.s. | * | *** |

**Table S19. Differences between the theoretical additive $OD_{600}$ and the observed $OD_{600}$ of cultures grown in YM-25 + GLY.**

The calculation is based on the values listed in Tables S6 and S7 (for the theoretical additive values) and Table S14 (for the observed values). See also Equations 1-4 in the main text. The red font indicates a positive-value difference and grey font a negative-value difference.

| Group | Subgroup | Strain | Precultivated on PDA (Exp. 1 – day 5.7) | | | | | | Precultivated on YM-10 agar (Exp. 2 – day 5.0) | | | | | |
|---|---|---|---|---|---|---|---|---|---|---|---|---|---|---|
| | | | Theoretical additive | | | **Observed – theoretical additive** | | | Theoretical additive | | | **Observed –theoretical additive** | | |
| | | | YM-25 + YM-GLY$_{0.2}$ | YM-25 + YM-GLY$_{1.0}$ | YM-25 + YM-GLY$_{5.0}$ | **GLY$_{0.2}$** | **GLY$_{1.0}$** | **GLY$_{5.0}$** | YM-25 + YM-GLY$_{0.2}$ | YM-25 + YM-GLY$_{1.0}$ | YM-25 + YM-GLY$_{5.0}$ | **GLY$_{0.2}$** | **GLY$_{1.0}$** | **GLY$_{5.0}$** |
| *S. cerevisiae* | Outlier | Y93 | 0.7884 | 0.2598 | 0.3417 | **-0.2856** | **0.2585** | **0.1288** | 0.8769 | 0.7753 | 0.1781 | **-0.6831** | **-0.5292** | **0.0573** |
| | Lactate – (15) | Y1129 | 0.0354 | 0.0168 | 0.0029 | **-0.0347** | **0.0138** | **-0.0014** | 0.0073 | 0.0043 | 0.0054 | **-0.0033** | **-0.0033** | **0.0002** |
| | | Y1703 | 0.0248 | 0.0228 | 0.0886 | **-0.0094** | **0.0582** | **0.0417** | 0.7590 | 0.0436 | 0.5364 | **-0.7241** | **0.0261** | **-0.3845** |
| | | Y1671 | 0.0669 | 0.0777 | 0.1359 | **0.0981** | **-0.0379** | **-0.0113** | 0.0423 | 0.0984 | 0.0996 | **0.1966** | **0.0762** | **0.1121** |
| | | Y1139 | 0.0208 | 0.0261 | 0.0537 | **0.1361** | **0.0375** | **0.1937** | 0.0877 | 0.0843 | 0.0985 | **0.0354** | **0.1363** | **0.0246** |
| | | Y1674 | 0.6337 | 0.0315 | 0.0499 | **-0.5957** | **0.2447** | **0.134** | 0.7690 | 0.0393 | 0.0533 | **-0.5709** | **0.1486** | **0.1308** |
| | | Y655 | 0.1155 | 0.0475 | 0.1045 | **0.073** | **0.2251** | **-0.0513** | 0.0393 | 0.0427 | 0.0692 | **0.0696** | **0.1813** | **-0.0094** |
| | Lactate – (70) | Y1122 | 0.0535 | 0.0324 | 0.0319 | **0.45** | **0.1569** | **0.1508** | 0.0438 | 0.0509 | 0.3392 | **-0.0201** | **0.0346** | **-0.1369** |
| | | Y682 | 0.7133 | 0.061 | 0.0787 | **-0.0908** | **0.6235** | **0.678** | 0.0463 | 0.1070 | 0.0658 | **-0.0081** | **0.4876** | **-0.0606** |
| | | Y1062 | 0.0522 | 0.0807 | 0.2753 | **0.7841** | **0.7519** | **0.4584** | 0.1267 | 0.1125 | 0.1999 | **0.1033** | **0.5669** | **0.0784** |
| | | Y634 | 0.672 | 0.06 | 0.1052 | **-0.0486** | **0.6279** | **0.6824** | 0.0932 | 0.0545 | 0.5227 | **0.1610** | **-0.0180** | **-0.5034** |
| | | Y1136 | 0.0262 | 0.0999 | 0.5166 | **0.5638** | **0.4868** | **-0.441** | 0.0643 | 0.1053 | 0.5851 | **0.1610** | **0.0476** | **-0.4529** |
| | | Y743 | 0.8548 | 0.0916 | 0.1752 | **0.0314** | **0.8368** | **0.4469** | 0.0931 | 0.0751 | 0.1231 | **0.3209** | **0.4272** | **-0.0563** |
| | | Y649 | 0.6445 | 0.0705 | 0.2621 | **-0.182** | **0.6091** | **0.6605** | 0.0459 | 0.0402 | 0.4050 | **-0.0259** | **0.3111** | **-0.2290** |
| | | Y1111 | 0.0554 | 0.0524 | 0.0637 | **0.4792** | **0.5882** | **0.0164** | 0.0565 | 0.0725 | 0.0527 | **-0.0340** | **0.0644** | **0.0182** |
| | | Y1705 | 0.5968 | 0.1228 | 0.7654 | **-0.0994** | **1.0785** | **0.1499** | 0.8963 | 0.1048 | 0.1681 | **-0.8318** | **-0.0454** | **0.2747** |
| | Lactate + | Y1242 | 1.1696 | 0.6607 | 0.7029 | **-0.3312** | **0.2301** | **0.0482** | 1.0529 | 1.0165 | 0.3933 | **-0.2098** | **-0.2060** | **0.2442** |
| | | Y1061 | 0.1801 | 0.1836 | 0.2397 | **0.7293** | **0.6418** | **0.5216** | 0.1247 | 0.8519 | 0.1779 | **0.2786** | **-0.5730** | **0.1040** |
| | | Y1624 | 0.9784 | 0.9701 | 1.0721 | **0.2045** | **0.2052** | **0.0525** | 0.5626 | 1.2216 | 1.0606 | **0.6014** | **-0.0695** | **-0.1223** |
| | | Y1613 | 1.6636 | 1.0041 | 1.1194 | **-0.4889** | **0.0964** | **-0.0281** | 0.8574 | 0.8635 | 1.2555 | **0.1728** | **0.2262** | **-0.3314** |
| | | Y683 | 0.7475 | 0.75 | 0.8751 | **0.4032** | **0.4244** | **0.1555** | 1.3735 | 0.5578 | 0.5901 | **-0.2440** | **0.5680** | **0.3341** |
| | | Y1584 | 1.1049 | 1.0962 | 1.2644 | **0.111** | **0.0329** | **-0.2447** | 0.8854 | 0.8730 | 0.9355 | **0.2524** | **0.2344** | **0.0344** |
| Non *S. cerevisiae* | Lactate – | Y1125 | 0.7155 | 1.464 | 1.655 | **-0.1718** | **-0.1647** | **0.1496** | 0.6226 | 0.6618 | 1.6473 | **0.0838** | **0.8088** | **0.1830** |
| | Lactate + | Y1728 | 2.0486 | 2.8174 | 3.1807 | **-0.5484** | **-1.1592** | **-1.2064** | 2.0907 | 2.9798 | 3.2459 | **-0.7839** | **-1.2697** | **-1.3175** |

#### Figure S13. Analysis of interactions between glycerol- and lactate- utilizations.

The legend is described in Figure 4A in the main text. The number on the x-axis indicates the concentration (% w/v) of glycerol. The yellow background indicates precultivation on PDA and pink on YM-10 agar. The source data are presented in Tables S6, S7, and S19.

| **Precultivated on PDA** (Exp. 1 – day 5.7) | | **Precultivated on YM-10 agar** (Exp. 2 – day 5.0) | |
|---|---|---|---|
| Observed vs. theoretical additive $OD_{600}$ | Synergism analysis | Observed vs. theoretical additive $OD_{600}$ | Synergism analysis |

##### *S. cerevisiae* – Outlier

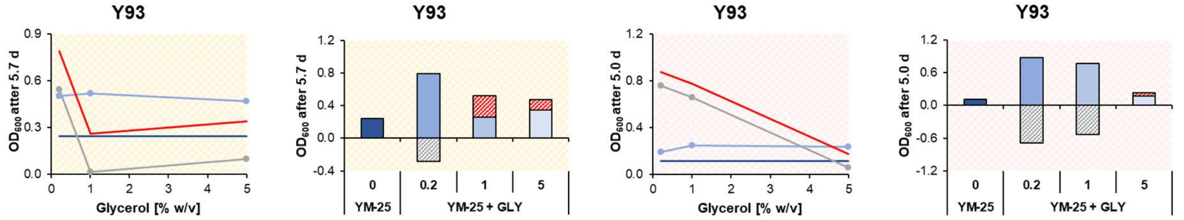

##### *S. cerevisiae* – Lactate - (15)

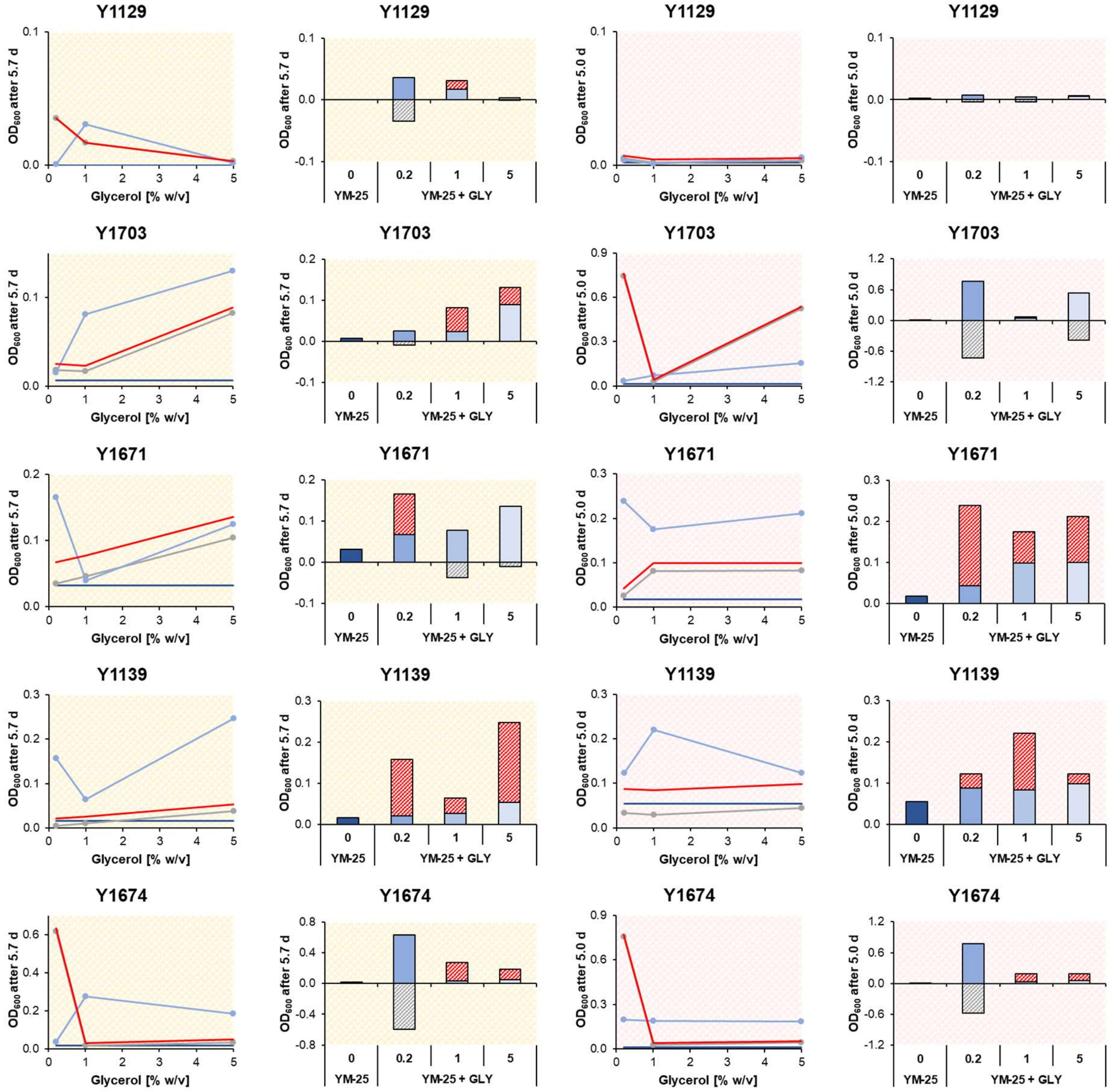

| **Precultivated on PDA** (Exp. 1 – day 5.7) | | **Precultivated on YM-10 agar** (Exp. 2 – day 5.0) | |
|---|---|---|---|
| Observed vs. theoretical additive $OD_{600}$ | Synergism analysis | Observed vs. theoretical additive $OD_{600}$ | Synergism analysis |

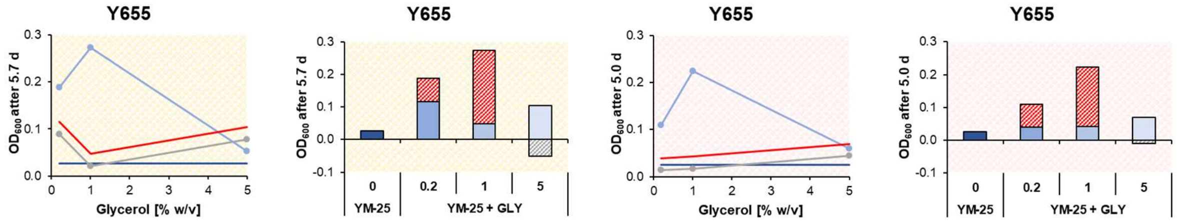

*S. cerevisiae* – Lactate – (70)

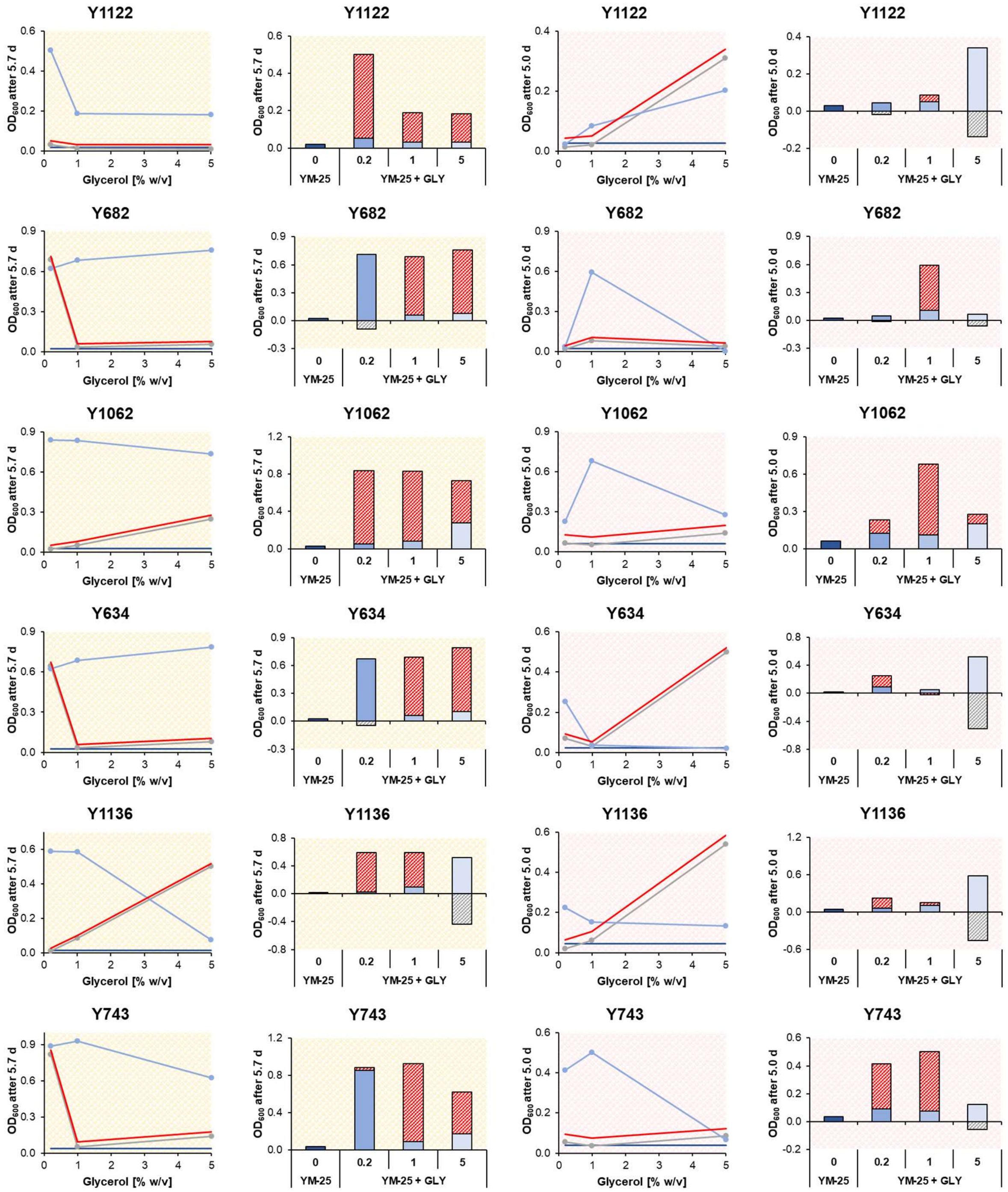

| **Precultivated on PDA** (Exp. 1 – day 5.7) | | **Precultivated on YM-10 agar** (Exp. 2 – day 5.0) | |
|---|---|---|---|
| Observed vs. theoretical additive $OD_{600}$ | Synergism analysis | Observed vs. theoretical additive $OD_{600}$ | Synergism analysis |

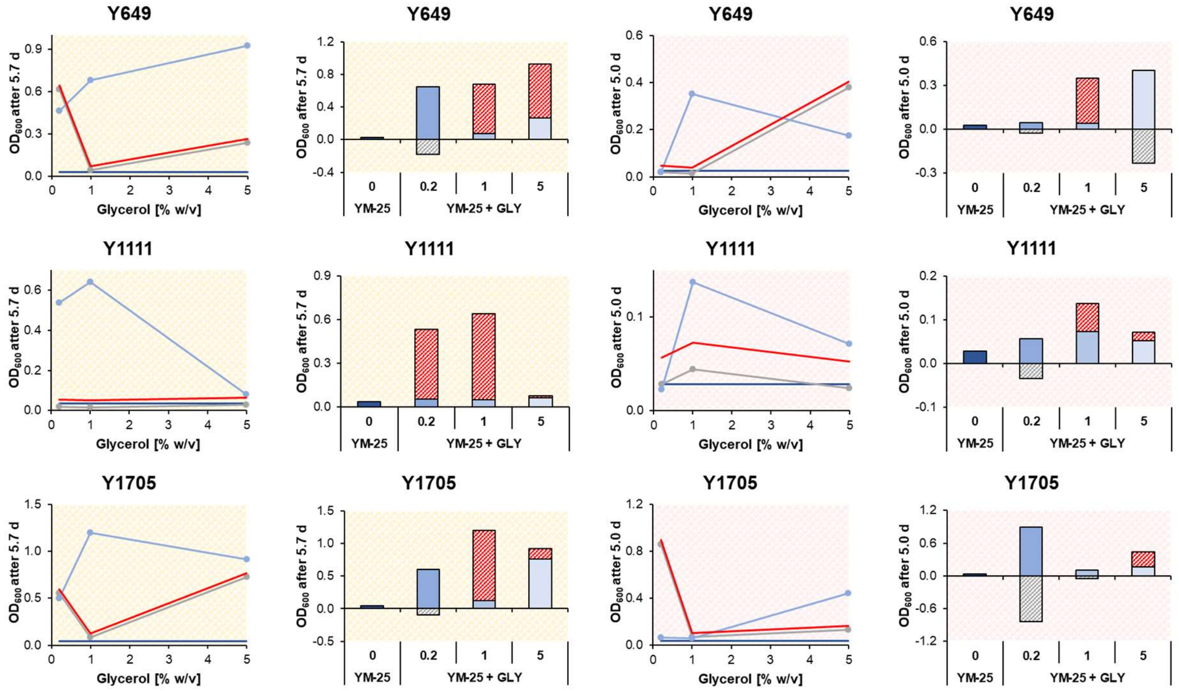

*S. cerevisiae* – **Lactate +**

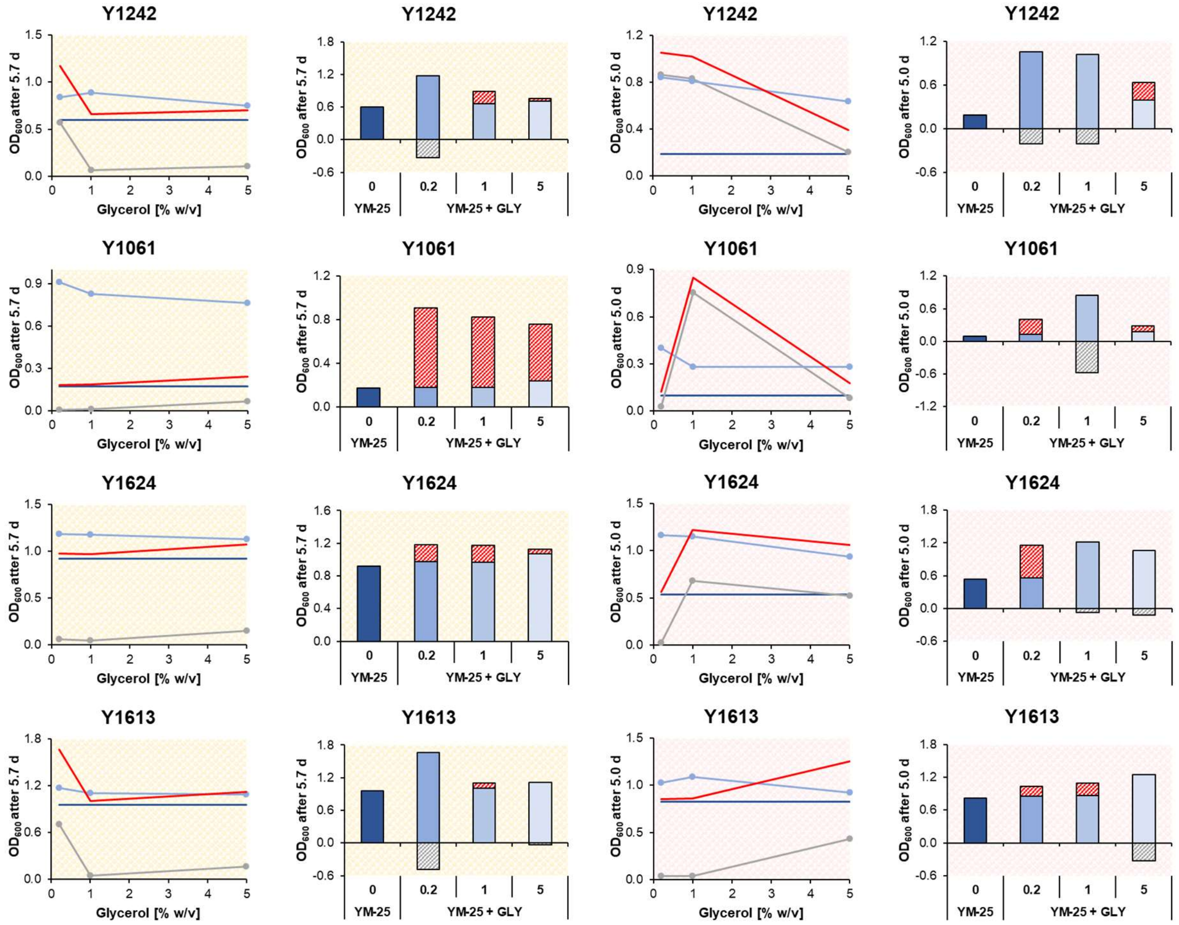

| **Precultivated on PDA** (Exp. 1 – day 5.7) | | **Precultivated on YM-10 agar** (Exp. 2 – day 5.0) | |
|---|---|---|---|
| Observed vs. theoretical additive $OD_{600}$ | Synergism analysis | Observed vs. theoretical additive $OD_{600}$ | Synergism analysis |

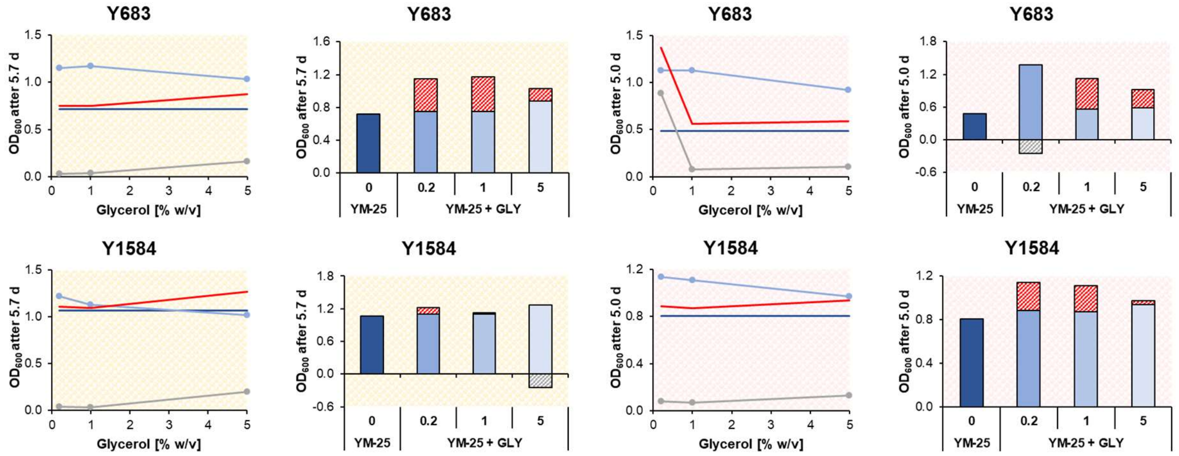

#### Non *S. cerevisiae* – Lactate -

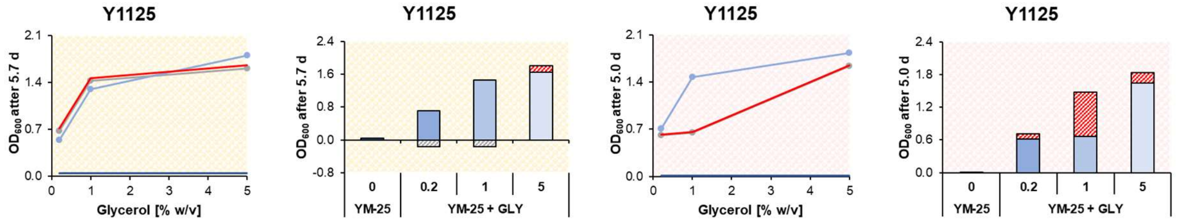

#### Non *S. cerevisiae* – Lactate +

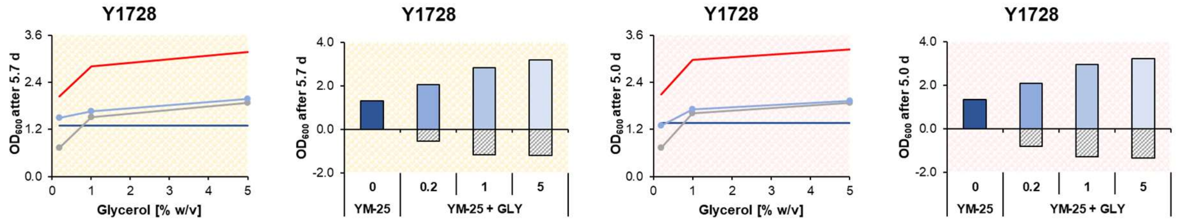

#### Table S20. Analysis of glycerol/lactate molar ratio that resulted in the highest $OD_{600}$ value.

The determination is based on the values listed in Table S14.

| Group | Subgroup | Strain | Highest $OD_{600}$ value | | | | | |
|---|---|---|---|---|---|---|---|---|
| | | | Precultivated on PDA (Exp. 1 – day 5.7) | | | Precultivated on YM-10 agar (Exp. 2 – day 5.0) | | |
| | | | YM-25 + $GLY_{0.2}$ | YM-25 + $GLY_{1.0}$ | YM-25 + $GLY_{5.0}$ | YM-25 + $GLY_{0.2}$ | YM-25 + $GLY_{1.0}$ | YM-25 + $GLY_{5.0}$ |
| *S. cerevisiae* | **Outlier** | Y93 | | 1 | | | 1 | |
| | **Lactate – (15)** | Y1129 | | 1 | | | | 1 |
| | | Y1703 | | | 1 | | | 1 |
| | | Y1671 | 1 | | | 1 | | |
| | | Y1139 | | | 1 | | 1 | |
| | | Y1674 | | 1 | | 1 | | |
| | | Y655 | | 1 | | | 1 | |
| | **Lactate – (70)** | Y1122 | 1 | | | | | 1 |
| | | Y682 | | | 1 | | 1 | |
| | | Y1062 | 1 | | | | 1 | |
| | | Y634 | | | 1 | 1 | | |
| | | Y1136 | 1 | | | 1 | | |
| | | Y743 | | 1 | | | 1 | |
| | | Y649 | | | 1 | | 1 | |
| | | Y1111 | | 1 | | | 1 | |
| | | Y1705 | | 1 | | | | 1 |
| | **Lactate +** | Y1242 | | 1 | | 1 | | |
| | | Y1061 | 1 | | | 1 | | |
| | | Y1624 | 1 | | | 1 | | |
| | | Y1613 | 1 | | | | 1 | |
| | | Y683 | | 1 | | 1 | | |
| | | Y1584 | 1 | | | 1 | | |
| | TOTAL (*n* = 22) | | **8** | **9** | **5** | **9** | **9** | **4** |
| | | | 36% | 41% | 23% | 41% | 41% | 18% |
| Non *S. cerevisiae* | **Lactate –** | Y1125 | | | 1 | | | 1 |
| | **Lactate +** | Y1728 | | | 1 | | | 1 |

**Figure S14. $OD_{600}$ in YM-25 or one of the three variants of YM-25 + GLY as a function of the medium's initial glycerol/lactate molar ratio.**

The yellow background indicates precultivation on PDA and pink on YM-10 agar. Source data: Tables S6 and S7.

*S. cerevisiae* - Outlier | *S. cerevisiae* – Lactate – (15)

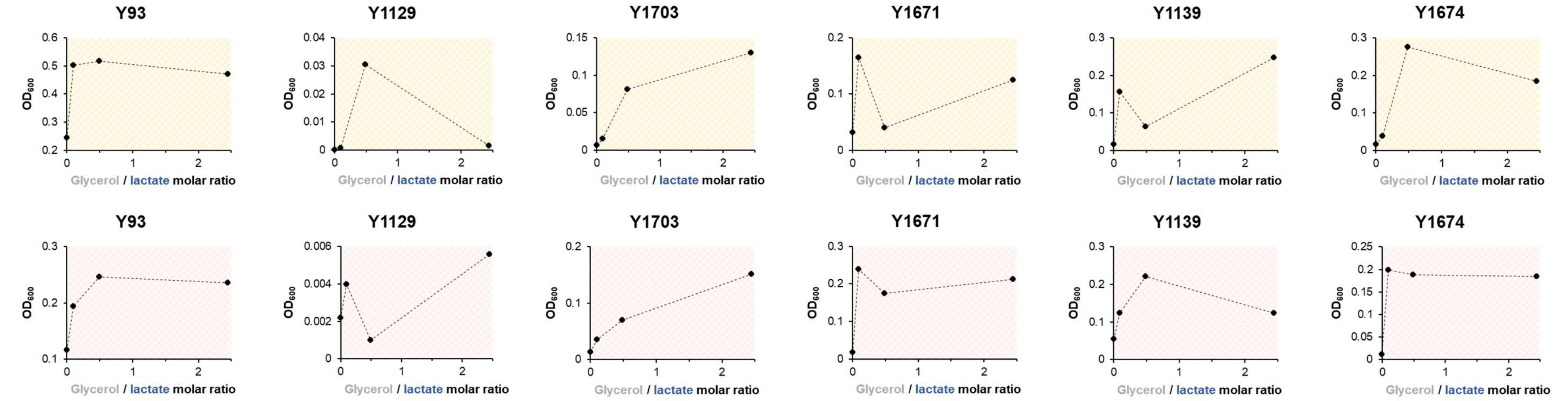

*S. cerevisiae* – Lactate – (15) | *S. cerevisiae* – Lactate – (70)

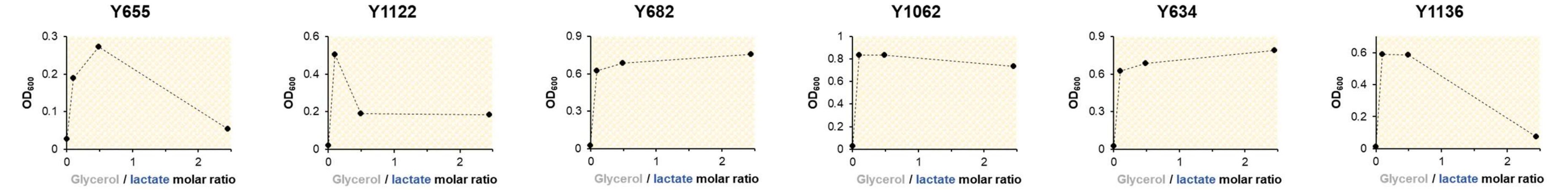

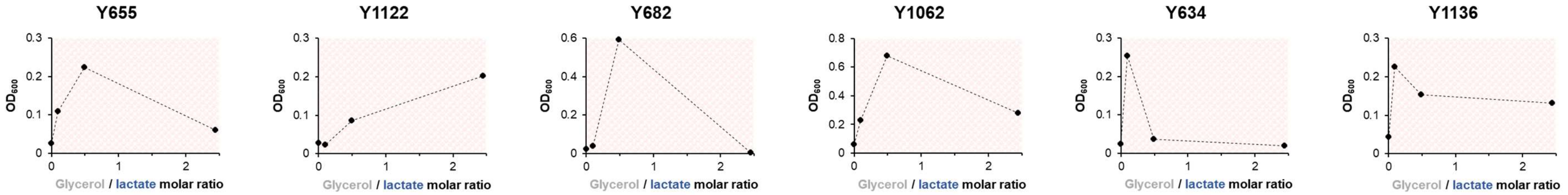

*S. cerevisiae* – Lactate – (70)

*S. cerevisiae* – Lactate +

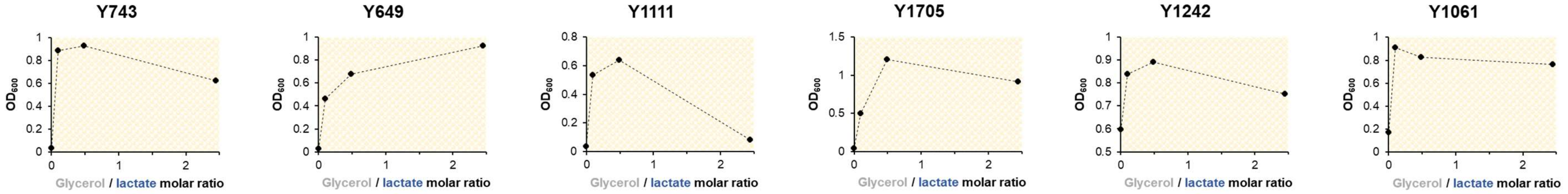

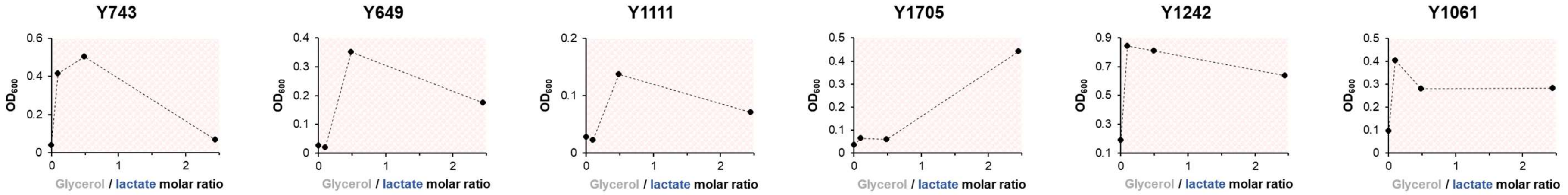

*S. cerevisiae* – Lactate +

Non *S. cerevisiae* – Lactate -

Non *S. cerevisiae* – Lactate +

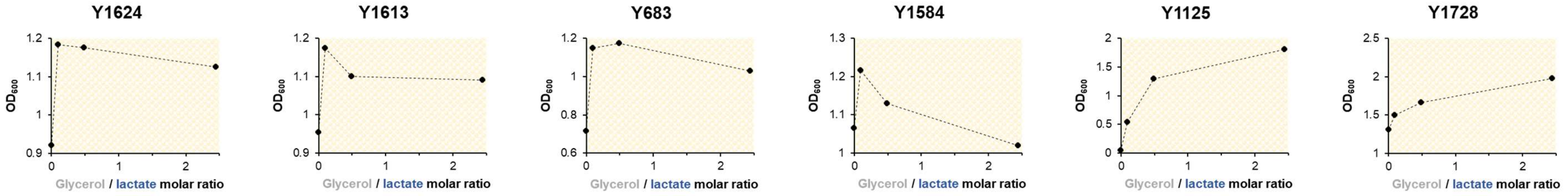

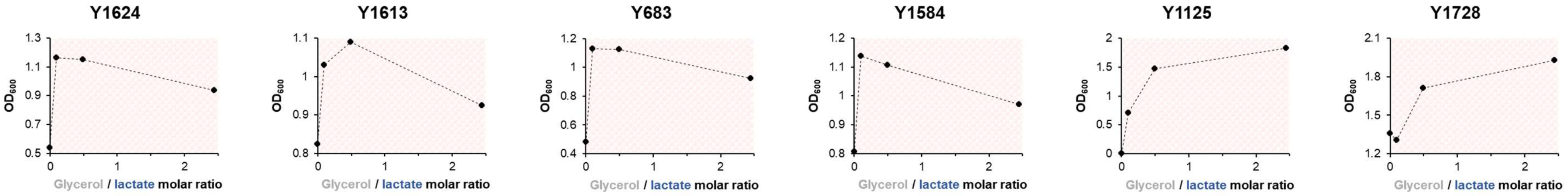
Y1624
Y1613
Y683
Y1584
Y1125
Y1728
OD600
Glycerol / lactate molar ratio

#### Note S3. Maximum theoretical growth yield in YM-25 and YM-GLY relative to that YM-21.

The maximum theoretical growth yield is the one solely determined by the substrate's chemical limit. In other words, here we assume that there is no additional limit imposed by the metabolic landscape used by the cell to extract carbon and energy from the substrate. One measure to describe a substrate's chemical limit is its heat of combustion [10].

**Table S21. Heat of combustion of the typical *S. cerevisiae* biomass and select substrates.**

The value for *S. cerevisiae* biomass is obtained from [10]. All other data are from the NIST Chemistry WebBook (https://webbook.nist.gov/chemistry/). If several values and/or ranges are listed, the largest mean (i.e., average) value is listed.

| Substrate | Chemical formula | Heat of combustion | |
|---|---|---|---|
| | | $\left[\frac{kJ}{mol}\right]$ | $\left[\frac{kJ}{Cmol}\right]$ |
| *S. cerevisiae* biomass | $C_1H_{1.8}O_{0.5}N_{0.2}$ | 560 | 560 |
| **Glucose** | $C_6H_{12}O_6$ | 2805.0 | 467.5 |
| **Lactic acid** | $C_3H_6O_3$ | 1362.8 | 454.3 |
| **Glycerol** | $C_3H_8O_3$ | 1654.1 | 551.4 |

Assuming an identical combustion efficiency between substrates, the maximum theoretical growth yield on a substrate relative to that on glucose can be calculated using the values in Table S21.

$$\frac{Max.\,theoretical\ growth\ yield_{\textbf{Lactic acid}}}{Max.\,theoretical\ growth\ yield_{\textbf{Glucose}}} = \frac{454.3\ \frac{kJ}{Cmol}}{467.5\ \frac{kJ}{Cmol}} = \sim 97\ \% \qquad \text{Equation S8}$$

$$\frac{Max.\,theoretical\ growth\ yield_{\textbf{Glycerol}}}{Max.\,theoretical\ growth\ yield_{\textbf{Glucose}}} = \frac{551.4\ \frac{kJ}{Cmol}}{467.5\ \frac{kJ}{Cmol}} = \sim 118\ \% \qquad \text{Equation S9}$$

The highest observed $OD_{600}$ percentage in YM-25 + GLY relative to YM-21 for *S. cerevisiae* was 91.19% for "Lactate – (70)" strains (i.e., Y1705 precultivated on PDA) and 83.73% for "Lactate +" strains (i.e., Y1584 precultivated on PDA). Those values are respectively ~94% and ~86% of the maximum theoretical value being ~97% (Equation S8). The lower-than-theoretical observed value indicates that the biomass synthesis process for *S. cerevisiae* on a glycerol/lactate mixture is still less efficient than that on glucose.

The $OD_{600}$ percentage in YM-$GLY_{5.0}$ relative to YM-21 for Y1728 (*C. lusitaniae*) and Y1125 (*W. versatilis*) was 118.24% and 123.50%, respectively, thus being comparable or slightly higher than the maximum theoretical value of ~118% (Equation S9). A higher-than-theoretical observed value indicates that biomass synthesis of those two strains on glycerol is more efficient than on glucose, which is indeed reported for a wide range of non-conventional yeasts [11–14].

**Figure S15. DL-Lactic acid calibration curve for quantitative HPLC analysis of spent media.**

Note that the unit of concentration was mistakenly written as “(ppm)” in the table; it should say “[g/L]”.

ID# : 1
Name : Lactic acid
Quantitative Method : External Standard
Function : f(x)=1.10791e+006*x+0
Rr1=0.9999991 Rr2=0.9999981

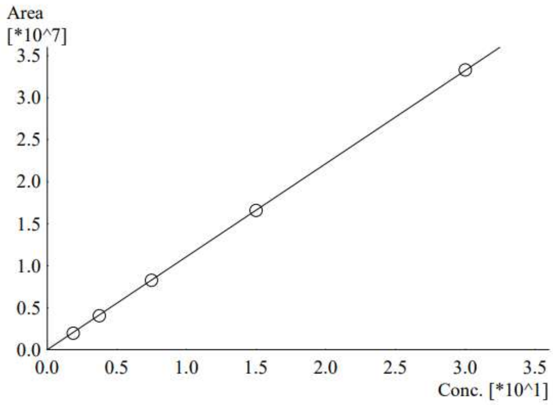

| # | Conc.(ppm) | MeanArea | Area |
|---|---|---|---|
| 1 | 1.875 | 1964648 | 1964648 |
| 2 | 3.75 | 4044468 | 4044468 |
| 3 | 7.5 | 8254797 | 8254797 |
| 4 | 15 | 16562509 | 16562509 |
| 5 | 30 | 33299804 | 33299804 |

**Figure S16. Chromatograms of residual (extracellular) lactate of select cultures from Experiment 3.**

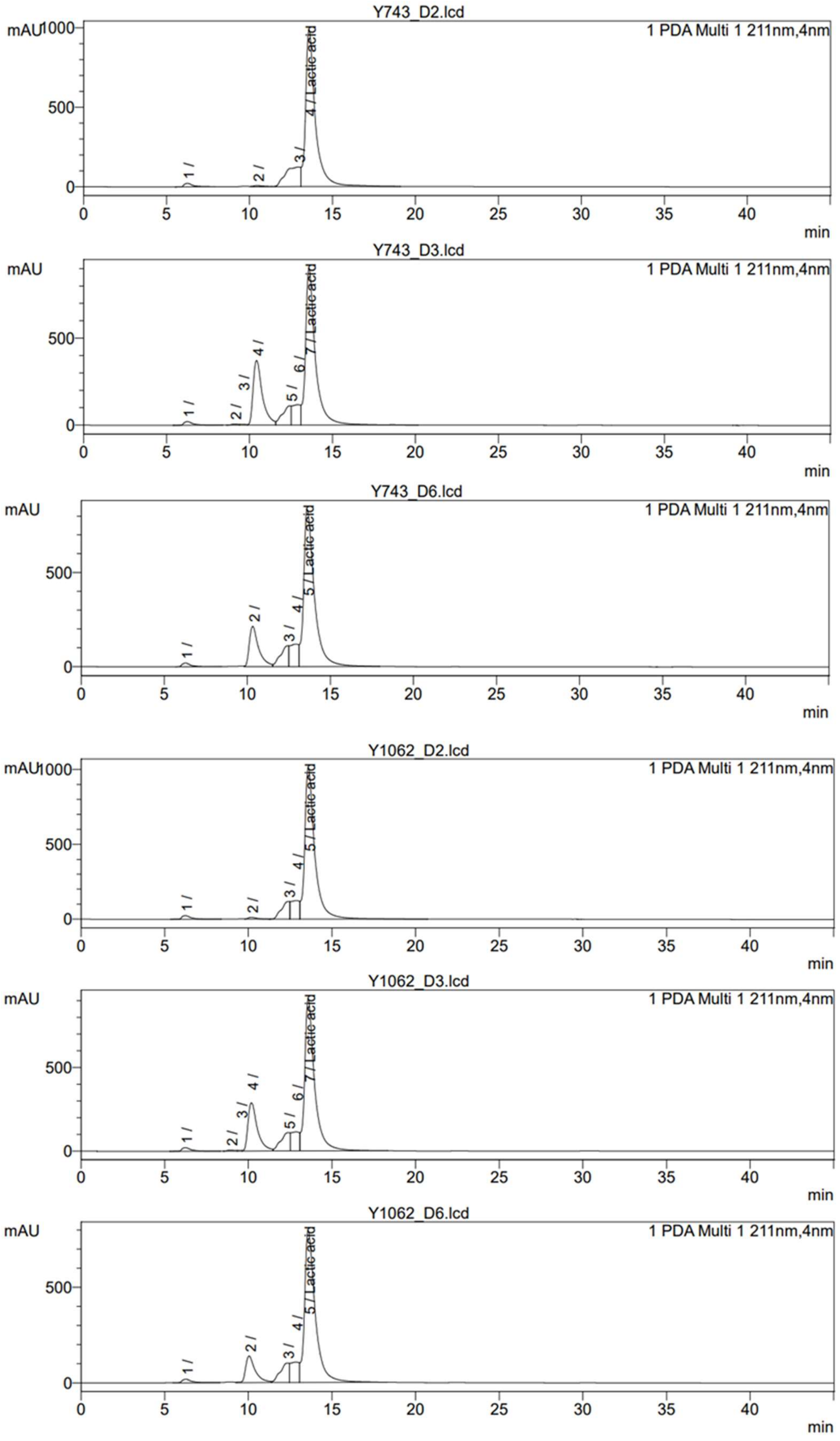

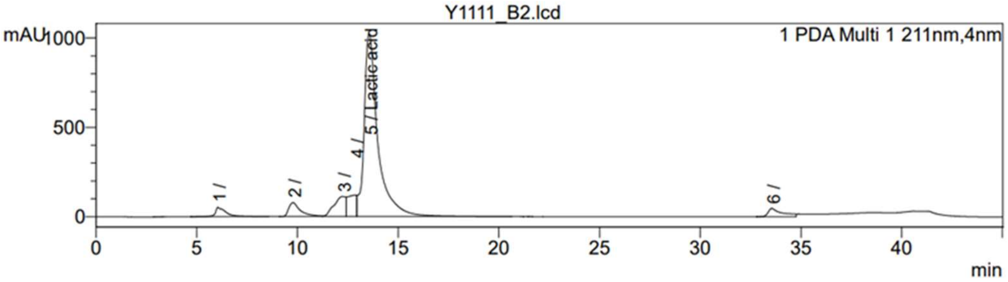
Y1111_B2.lcd
mAU
1 PDA Multi 1 211nm,4nm
5 / Lactic acid
min

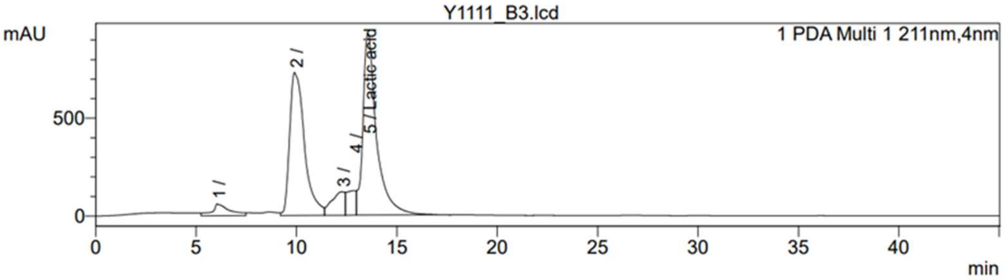
Y1111_B3.lcd
mAU
1 PDA Multi 1 211nm,4nm
5 / Lactic acid
min

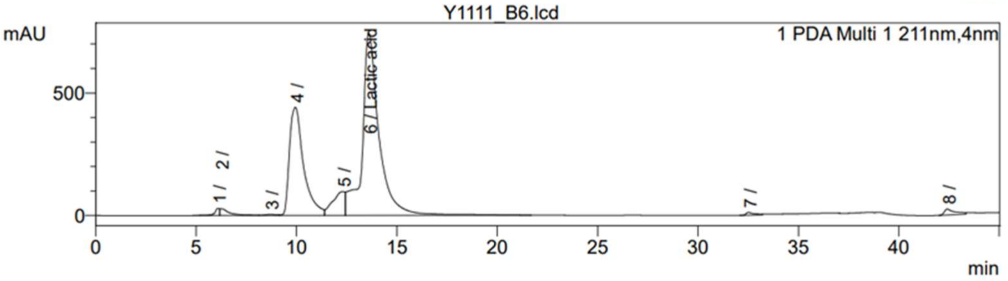
Y1111_B6.lcd
mAU
1 PDA Multi 1 211nm,4nm
6 / Lactic acid
min

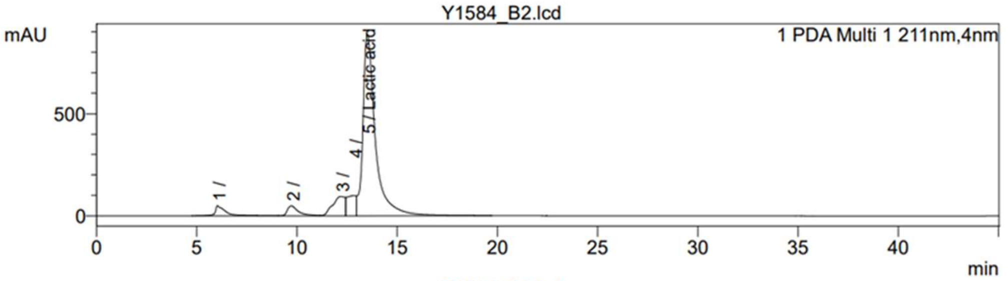
Y1584_B2.lcd
mAU
1 PDA Multi 1 211nm,4nm
5 / Lactic acid
min

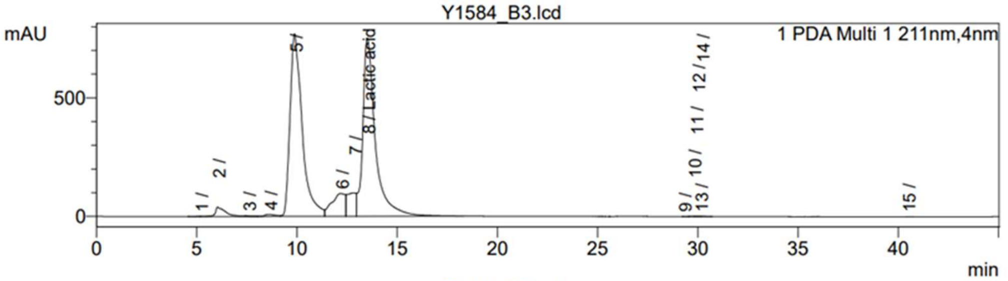
Y1584_B3.lcd
mAU
1 PDA Multi 1 211nm,4nm
8 / Lactic acid
min

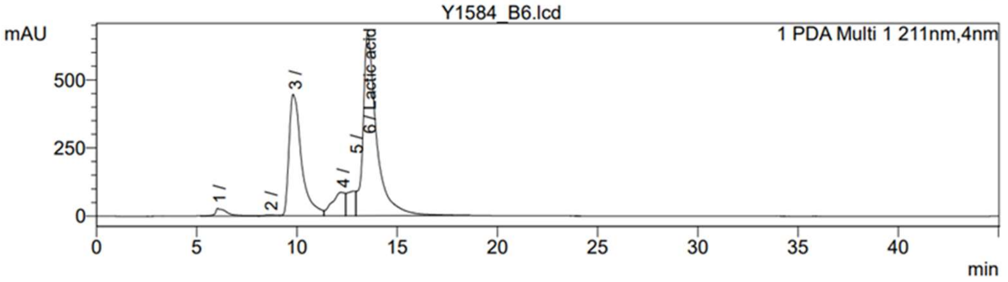
Y1584_B6.lcd
mAU
1 PDA Multi 1 211nm,4nm
6 / Lactic acid
min

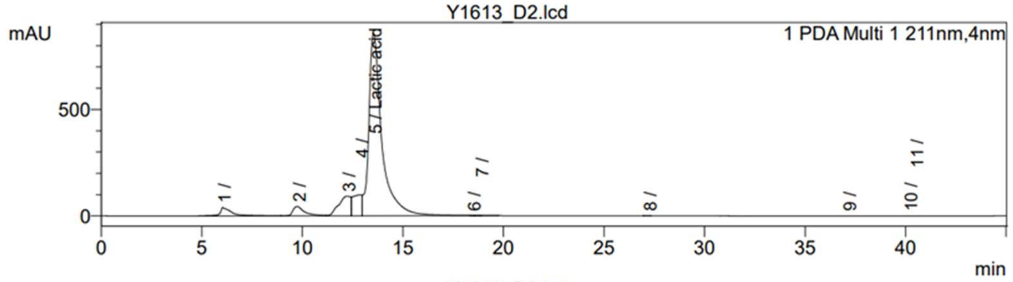
Y1613_D2.lcd
mAU
1 PDA Multi 1 211nm,4nm
1 /
2 /
3 /
4 /
5 / Lactic acid
6 /
7 /
8 /
9 /
10 /
11 /
min

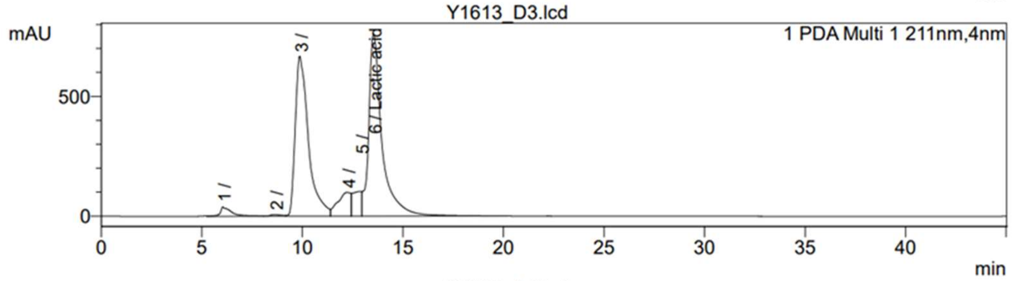
Y1613_D3.lcd
mAU
1 PDA Multi 1 211nm,4nm
1 /
2 /
3 /
4 /
5 /
6 / Lactic acid
min

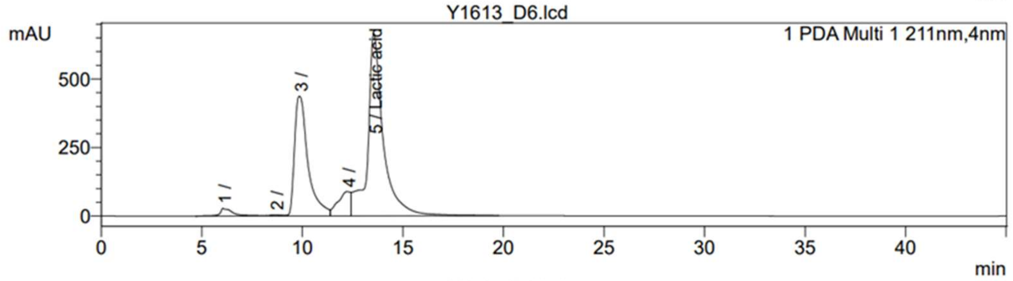
Y1613_D6.lcd
mAU
1 PDA Multi 1 211nm,4nm
1 /
2 /
3 /
4 /
5 / Lactic acid
min

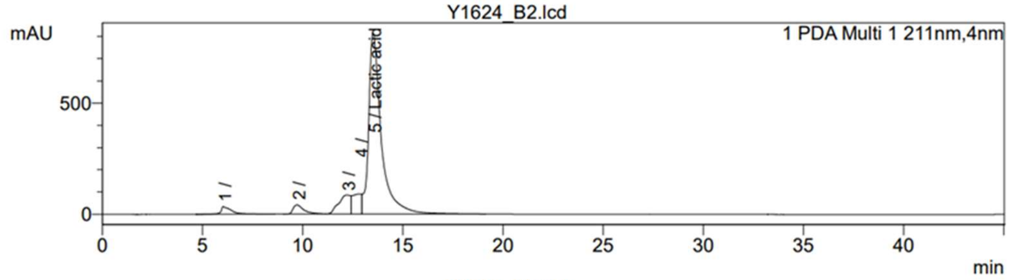
Y1624_B2.lcd
mAU
1 PDA Multi 1 211nm,4nm
1 /
2 /
3 /
4 /
5 / Lactic acid
min

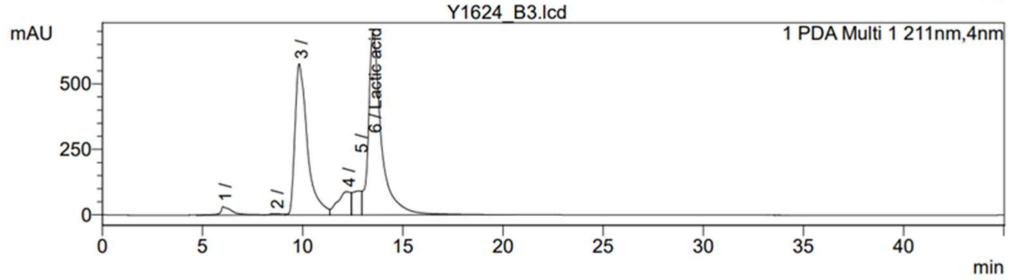
Y1624_B3.lcd
mAU
1 PDA Multi 1 211nm,4nm
1 /
2 /
3 /
4 /
5 /
6 / Lactic acid
min

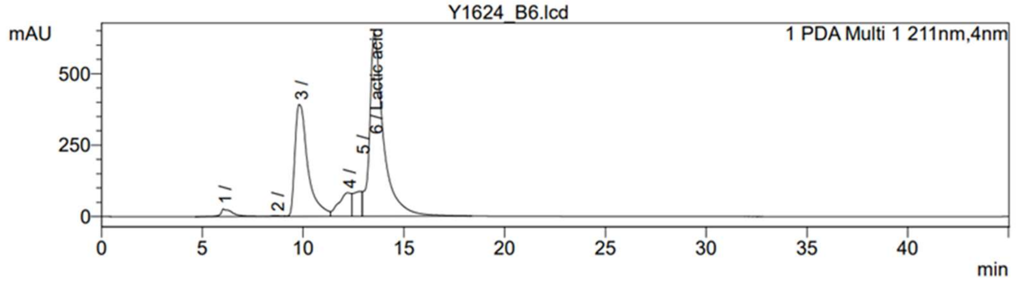
Y1624_B6.lcd
mAU
1 PDA Multi 1 211nm,4nm
1 /
2 /
3 /
4 /
5 /
6 / Lactic acid
min

**Table S22. Extracellular lactate concentration in spent media of select cultures from Experiment 3.**

Based on the chromatograms presented in Figure S16.

| Group | Strain | Medium | Glycerol/ lactate molar ratio | Sample name | Retention time [min] | Area | Height | [Lactate] [g/L] |
|---|---|---|---|---|---|---|---|---|
| **Lactate – (70)** | Y743 | YM-25 | 0 | Y743_D2 | 13.586 | 40515604 | 985899 | 36.569 |
| | | YM-25 + $GLY_{0.25}$ | 0.12 | Y743_D3 | 13.583 | 37362059 | 900661 | 33.723 |
| | | YM-25 + $GLY_{5.0}$ | 2.45 | Y743_D6 | 13.601 | 37473698 | 835311 | 33.824 |
| | Y1062 | YM-25 | 0 | Y1062_D2 | 13.583 | 42135665 | 1011882 | 38.032 |
| | | YM-25 + $GLY_{0.25}$ | 0.12 | Y1062_D3 | 13.579 | 37260680 | 911148 | 33.632 |
| | | YM-25 + $GLY_{5.0}$ | 2.45 | Y1062_D6 | 13.575 | 35382844 | 795641 | 31.937 |
| | Y1111 | YM-25 | 0 | Y1111_B2 | 13.560 | 48819164 | 1022794 | 44.064 |
| | | YM-25 + $GLY_{0.25}$ | 0.12 | Y1111_B3 | 13.561 | 44000023 | 928365 | 39.714 |
| | | YM-25 + $GLY_{5.0}$ | 2.45 | Y1111_B6 | 13.602 | 43235571 | 743407 | 39.024 |
| **Lactate +** | Y1584 | YM-25 | 0 | Y1584_B2 | 13.506 | 38189140 | 887976 | 34.470 |
| | | YM-25 + $GLY_{0.25}$ | 0.12 | Y1584_B3 | 13.507 | 32419448 | 752810 | 29.262 |
| | | YM-25 + $GLY_{5.0}$ | 2.45 | Y1584_B6 | 13.525 | 31411523 | 668618 | 28.352 |
| | Y1613 | YM-25 | 0 | Y1613_D2 | 13.534 | 38711803 | 859606 | 34.941 |
| | | YM-25 + $GLY_{0.25}$ | 0.12 | Y1613_D3 | 13.538 | 35103670 | 762544 | 31.685 |
| | | YM-25 + $GLY_{5.0}$ | 2.45 | Y1613_D6 | 13.549 | 35893931 | 666420 | 32.398 |
| | Y1624 | YM-25 | 0 | Y1624_B2 | 13.514 | 35621084 | 813919 | 32.152 |
| | | YM-25 + $GLY_{0.25}$ | 0.12 | Y1624_B3 | 13.512 | 31295520 | 694640 | 28.247 |
| | | YM-25 + $GLY_{5.0}$ | 2.45 | Y1624_B6 | 13.536 | 30584790 | 639847 | 27.606 |

**Table S23. Relative extracellular lactate concentration in spent media of select cultures from Experiment 3.**

Based on the data presented in Table S22.

| Group | Strain | $[Lactate]$ [g/L] | | | $Relative\ [Lactate] = \frac{[Lactate]_{YM-25+GLY}}{[Lactate]_{YM-25}}$ | |
|---|---|---|---|---|---|---|
| | | YM-25 | YM-25 + $GLY_{0.25}$ | YM-25 + $GLY_{5.0}$ | YM-25 + $GLY_{0.25}$ | YM-25 + $GLY_{5.0}$ |
| **Lactate – (70)** | Y743 | 36.569 | 33.723 | 33.824 | 0.922 | 0.925 |
| | Y1062 | 38.032 | 33.632 | 31.937 | 0.884 | 0.840 |
| | Y1111 | 44.064 | 39.714 | 39.024 | 0.901 | 0.886 |
| **Lactate +** | Y1584 | 34.470 | 29.262 | 28.352 | 0.849 | 0.823 |
| | Y1613 | 34.941 | 31.685 | 32.398 | 0.907 | 0.927 |
| | Y1624 | 32.152 | 28.247 | 27.606 | 0.879 | 0.859 |

#### Table S24. Relative $OD_{600}$ of select cultures from Experiment 3.

Based on the data presented in Table S8.

| Group | Strain | $[OD_{600}]$ | | | $Relative\ [OD_{600}] = \frac{[OD_{600}]_{YM-25+GLY}}{[OD_{600}]_{YM-25}}$ | |
|---|---|---|---|---|---|---|
| | | YM-25 | YM-25 + $GLY_{0.25}$ | YM-25 + $GLY_{5.0}$ | YM-25 + $GLY_{0.25}$ | YM-25 + $GLY_{5.0}$ |
| **Lactate – (70)** | Y743 | 0.0215 | 0.8956 | 0.4594 | 41.66 | 21.37 |
| | Y1062 | 0.0280 | 0.9018 | 0.5998 | 32.21 | 21.42 |
| | Y1111 | 0.0185 | 0.4072 | 0.2538 | 22.01 | 13.72 |
| **Lactate +** | Y1584 | 0.0251 | 0.9367 | 0.5233 | 37.32 | 20.85 |
| | Y1613 | 0.0204 | 0.8453 | 0.5701 | 41.44 | 27.95 |
| | Y1624 | 0.0166 | 0.6530 | 0.4334 | 39.34 | 26.11 |

**Table S25. Statistical analysis of relative $OD_{600}$ and relative extracellular lactate concentration in spent media of select cultures from Experiment 3.**

Based on the data presented in Tables S23 and S24. Statistical significance (*p*-value) is calculated using the TTEST function (two-tailed distribution, paired) in Microsoft Excel.

| Group | *n* | $Relative\ [OD_{600}]$ | | | | | $Relative\ [Lactate]$ | | | | |
|---|---|---|---|---|---|---|---|---|---|---|---|
| | | YM-25 + $GLY_{0.25}$ | | YM-25 + $GLY_{5.0}$ | | *p*-value | YM-25 + $GLY_{0.25}$ | | YM-25 + $GLY_{5.0}$ | | *p*-value |
| | | Average | Stdev | Average | Stdev | | Average | Stdev | Average | Stdev | |
| **Lactate – (70)** | 3 | 31.96 | 9.82 | 18.84 | 4.43 | 0.070 | 0.903 | 0.019 | 0.883 | 0.043 | 0.299 |
| **Lactate +** | 3 | 39.36 | 2.06 | 24.97 | 3.68 | 0.005 | 0.878 | 0.029 | 0.869 | 0.053 | 0.615 |
| **All strains** | 6 | 35.66 | 7.53 | 21.90 | 4.96 | 0.000 | 0.890 | 0.026 | 0.876 | 0.044 | 0.195 |

#### Note S4. Predicting the metabolic flux distribution of *S. cerevisiae* grown on lactate.

The maximum specific growth rates of *S. cerevisiae* CEN.PK113-7D on glucose, galactose, and pyruvate described in [9] were correlated relatively well with those of *S. cerevisiae* FY4 described in [15] (Figure 5R.17 in [9], $R^2 = 0.891$). The relatively strong correlation suggests that the *in vivo* metabolic flux distributions described in [15] can also describe to a reasonable extent the cultures described in [9]. Indeed there is a physicochemical basis supporting the rationale: the production rates of ATP and intracellular nutrients, both serving as inputs to the master growth regulator TOR [16], should be reflected by the biomass synthesis rate, described by no other than the maximum specific growth rate.

Using the same rationale, it is then reasonable to think that the metabolic flux distribution of *S. cerevisiae* on lactate can be predicted from that on pyruvate, provided that we know the specific growth rates of the same strain on each carbon source. The ratio of maximum specific growth rate between the two conditions should correlate with the ratio of ATP production rate. The information can then be used to determine the likely amount of extramitochondrial (i.e., cytosolic) pyruvate, which can be used to synthesize various cytosolic metabolites.

The maximum specific growth rate of *S. cerevisiae* CEN.PK113-7D on pyruvate ($\mu_{max\ PYRUVATE\ (YM-24)}$) described in [9] was around 0.256 $h^{-1}$ (Figure 5R.14A), while that on lactate ($\mu_{max\ LACTATE\ (YM-25)}$) was around 0.414 $h^{-1}$ (Figure 5R.11A). As such, Equation S10 can be stated.

$$\frac{\mu_{max\ LACTATE\ (YM-25)}}{\mu_{max\ PYRUVATE\ (YM-24)}} = \frac{0.414\ h^{-1}}{0.256\ h^{-1}} = 1.617 \qquad \text{Equation S10}$$

Assuming that the ratio of maximum specific growth rates reflects the ratio of ATP production rate, then Equation S11 can be stated.

$$\frac{ATP\ production\ rate_{LACTATE}}{ATP\ production\ rate_{PYRUVATE}} = \frac{\mu_{max\ LACTATE\ (YM-25)}}{\mu_{max\ PYRUVATE\ (YM-24)}} = 1.617 \qquad \text{Equation S11}$$

According to the metabolic flux distribution described in [15], the ATP production flux per 100 molecules of pyruvate at quasi-steady-state can be calculated as shown in Table S26. Note that when grown on pyruvate, *S. cerevisiae* can be assumed to produce ATP solely through the substrate-level phosphorylation in the TCA cycle (i.e., the succinyl-CoA ligase reaction) and the oxidative phosphorylation of NADH and $FADH_2$ generated by the same cycle. Further, owing to the absence of proton-pumping NADH dehydrogenase in the species [17], the amount of ATP molecule that can be generated per molecule of NADH or $FADH_2$ can be assumed as 2.

**Table S26. Calculation of ATP molecules produced per 100 intracellular pyruvate molecules according to the metabolic flux distribution described by Fendt and Sauer (2010).**

Note that the metabolic flux distribution used as the basis of this calculation is reproduced in Figure 6 of this study.

| Reaction | | NADH or $FADH_2$ equivalent | ATP equivalent |
|---|---|---|---|
| Substrate | Product | | |
| $PYRUVATE_{mito}$ | $acetyl\text{-}CoA_{mito}$ | 18 | 36 |
| $Isocitrate_{mito}$ | $\alpha\text{-}ketoglutarate_{mito}$ | 17 | 34 |
| $\alpha\text{-}ketoglutarate_{mito}$ | $succinate_{mito}$ | 13 | 26 |
| $succinyl\text{-}CoA_{mito}$ | $succinate_{mito}$ | - | 13 |
| $succinate_{mito}$ | $fumarate_{mito}$ | 13 | 26 |
| $malate_{mito}$ | $oxaloacetate_{mito}$ | 13 | 26 |
| TOTAL ATP produced per 100 molecules of pyruvate | | | 161 |

Based on the calculation in Table S26, Equation S12 can be stated, where $k$ is the relevant multiplication constant.

$$\boldsymbol{ATP\ production\ rate_{PYRUVATE}} = \frac{moles\ of\ ATP\ produced}{100\ moles\ of\ PYRUVATE} \cdot k = \mathbf{161}\ \boldsymbol{k} \qquad \text{Equation S12}$$

Substituting Equation S12 to Equation S11, followed by rearrangement, results in Equation S13.

$$\begin{aligned} \boldsymbol{ATP\ production\ rate_{LACTATE}} &= 1.617 \times ATP\ production\ rate_{PYRUVATE} \\ &= 1.617 \times 161\ k = \mathbf{260.337}\ \boldsymbol{k} \end{aligned} \qquad \text{Equation S13}$$

When *S. cerevisiae* is grown on lactate, ATP will be produced through not only the TCA cycle but also the lactate:cytochrome *c* oxidoreductase (LCR) reaction, with the production rate being one molecule of ATP per molecule of lactate reacting. As such, Equations S14 and S15 can be stated.

$$\begin{aligned} &\boldsymbol{ATP\ production\ rate_{LACTATE}} \\ &\quad = \boldsymbol{ATP\ production\ rate_{LACTATE\ via\ TCA\ cycle}} + \boldsymbol{ATP\ production\ rate_{LACTATE\ via\ LCR}} \end{aligned} \qquad \text{Equation S14}$$

$$\boldsymbol{ATP\ production\ rate_{LACTATE\ via\ LCR}} = \frac{moles\ of\ ATP\ produced}{100\ moles\ of\ PYRUVATE} \cdot k = \mathbf{100}\ \boldsymbol{k} \qquad \text{Equation S15}$$

Susbtituting Equations S13 and S15 to Equation S14, followed by rearrangement, results in Equation S16.

$$\begin{aligned} &260.337\ k = ATP\ production\ rate_{LACTATE\ via\ TCA\ cycle} + 100\ k \\ &\Rightarrow \boldsymbol{ATP\ production\ rate_{LACTATE\ via\ TCA\ cycle}} = 260.337\ k - 100\ k = \mathbf{160.337}\ \boldsymbol{k} \end{aligned} \qquad \text{Equation S16}$$

Comparing Equation S16 with Equation S12 results in Equation S17.

$$\begin{aligned} &ATP\ production\ rate_{LACTATE-TCA\ cycle} = 160.337\ k \approx 161 \cdot k = ATP\ production\ rate_{PYRUVATE} \\ &\Rightarrow \boldsymbol{ATP\ production\ rate_{LACTATE\ via\ TCA\ cycle}} \approx \boldsymbol{ATP\ production\ rate_{PYRUVATE}} \end{aligned} \qquad \text{Equation S17}$$

Equation S16 largely describes the distribution of metabolic fluxes emanating from lactate-derived $PYRUVATE_{mito}$, but not lactate-derived $PYRUVATE_{cyto}$. Since the magnitude of the latter should reflect as biomass level, the ratio of total growth on lactate to that on pyruvate can provide an idea of how different they are. Using the values reported in Figures 5R.12A and 5R.15A in [9], the ratio can be stated as Equation S18.

$$\frac{\boldsymbol{Total\ growth_{LACTATE\ (YM-25)}}}{\boldsymbol{Total\ growth_{PYRUVATE\ (YM-24)}}} = \frac{6.61}{5.74} = \mathbf{1.15} \qquad \text{Equation S18}$$

Equation S18 suggests that the distribution of metabolic fluxes emanating from lactate-derived $PYRUVATE_{cyto}$ is likely very similar to that of pyruvate-grown culture. Taking into consideration both Equations S17 and S18, it is justified to think that the metabolic flux distribution of *S. cerevisiae* grown on lactate is very similar to what was described in [15] for *S. cerevisiae* grown on pyruvate, i.e., Figure 6 of this study.

##### Table S27. Abundance of proteins directly involved in cytosolic NADPH synthesis in *S. cerevisiae* grown on ten different carbon sources.

The raw data are obtained from [18]. The strain was BY4742. The minimal medium was based on YNB (with amino acids and ammonium sulfate) and contained 2% of the designated carbon source. Note that Idp2 is the only member of the ICDH functional category. Stdev: standard deviation.

| Functional category | Protein ID | glucose | maltose | oleate | fructose | sucrose | trehalose | lactate | acetate | pryuvate | glycerol | **Median** | **Stdev** | **Stdev/ median** |
|---|---|---|---|---|---|---|---|---|---|---|---|---|---|---|
| Oxidative branch of the pentose phosphate pathway (oxPPP) | **Zwf1** | 19769.600 | 24801.500 | 18536.700 | 18667.100 | 21568.200 | 21872.900 | 20120.200 | 15269.700 | 21419.100 | 16796.000 | **19944.9** | **2739.0794** | **0.14** |
| | **Gnd1** | 27623.300 | 23487.100 | 13006.100 | 24934.500 | 30023.400 | 22605.000 | 9559.040 | 11233.100 | 19378.800 | 24566.800 | **40012.45** | **13364.459** | **0.33** |
| | **Gnd2** | 9696.050 | 38594.300 | 11596.300 | 8206.760 | 10770.700 | 24523.500 | 26056.700 | 5969.200 | 12566.400 | 16650.400 | **23046.05** | **7100.7733** | **0.31** |
| Aldehyde dehydrogenase (ALDD) | **Ald6** | 40778.100 | 27238.000 | 42529.500 | 43626.700 | 34310.600 | 36863.100 | 48455.700 | 34940.100 | 76680.100 | 39246.800 | **12081.35** | **10231.693** | **0.85** |
| | **Ald3** | 2432.480 | 61679.900 | 11160.100 | 1943.820 | 2331.250 | 37673.100 | 39593.000 | 7412.440 | 10783.800 | 25489.500 | **2918.525** | **2132.7123** | **0.73** |
| | **Ald2** | 1002.94 | 7283.39 | 3274.5 | 998.259 | 1026.7 | 4595 | 5582.57 | 1935.75 | 2562.55 | 3681.2 | **36487** | **19233.24** | **0.53** |
| Isocitrate dehydrogenase (ICDH) | **Idp2** | 2804.300 | 43068.100 | 52335.600 | 2537.250 | 2277.210 | 38933.500 | 44209.300 | 40426.400 | 34040.500 | 28437.000 | **10971.95** | **20363.106** | **1.86** |

**Table S28. Abundance of proteins that are directly or indirectly involved in cytosolic NADPH synthesis in various glucose-grown *S. cerevisiae* cultures.**

The abundance data are obtained from Table S4 in [19]. The description of the dataset is obtained from Table 1 of the same reference. The information regarding the strain background is obtained from the primary research paper generating the original dataset. Note that the strain(s) used in the study may carry deliberate mutations to assist protein abundance quantification. Idp1 abundance should influence the amount of NADPH generated by Idp2 since both enzymes compete for the same substrate (Figure 6). Meanwhile, Pos5 abundance should allow for a less abundant Idp1, as the former accommodates an alternative route for mitochondrial NADPH synthesis (Figure 6).

| Dataset | Primary research reference | Strain background (WT or mutants) | Medium | Growth phase | Abundance | | | | | | | Abundance ratio | | | | |
|---|---|---|---|---|---|---|---|---|---|---|---|---|---|---|---|---|
| | | | | | Direct | | | | | Indirect | | Direct | | | | Indirect |
| | | | | | Zwf1 | Ald6 | Ald2 | Ald3 | Idp2 | Idp1 | Pos5 | Ald6/Zwf1 | Ald2/Zwf1 | Ald3/Zwf1 | Idp2/Zwf1 | Idp1/Pos5 |
| LAHT | [20] | CEN.PK113-7D | minimal | chemostat | 54955 | 893087 | 8557 | 2676 | 3742 | | 439 | 16.2512 | 0.1557 | 0.0487 | 0.0681 | |
| NAG | [21] | W303 MATα | YPD | mid-log | 31981 | 33205 | 41767 | 23512 | | 28359 | 1886 | 1.0383 | 1.3060 | 0.7352 | | 15.0 |
| LU | [22] | DBY8724 | YPD | mid-log | 6205 | 244734 | 19488 | 24478 | | 19801 | | 39.4414 | 3.1407 | 3.9449 | | |
| WEB | [23] | S288C | YPD | mid-log | 20926 | 47226 | | | 3173 | 16510 | 4403 | 2.2568 | | | 0.1516 | 3.7 |
| THAK | [24] | | minimal | mid-log | 24256 | 79096 | 1646 | 11403 | 5713 | 29592 | 1830 | 3.2609 | 0.0679 | 0.4701 | 0.2355 | 16.2 |
| LAW | [25] | BY4742 | minimal | chemostat | 319059 | 28162 | | 2688 | 2920 | 31570 | 3619 | 0.0883 | | 0.0084 | 0.0092 | 8.7 |
| PENG | [26] | | minimal | early log | 21681 | 61067 | 2116 | 8886 | 7926 | 63154 | 1376 | 2.8166 | 0.0976 | 0.4099 | 0.3656 | 45.9 |
| DEN | [27] | BY4741 | minimal | steady state | 9477 | 711791 | | 4162 | | 6226 | 3469 | 75.1072 | | 0.4392 | | 1.8 |
| CHO | [28] | | minimal | mid-log | 9683 | 240018 | | | | 16003 | | 24.7876 | | | | |
| BRE | [29] | | | mid-log | 10152 | 652548 | | 6241 | | 23590 | | 64.2778 | | 0.6148 | | |
| MAZ | [30] | | | mid-log | 8087 | | | | | 9623 | | | | | | |
| TKA | [31] | | | mid-log | 12465 | 604046 | 1716 | 1747 | | 27112 | | 48.4594 | 0.1377 | 0.1402 | | |
| KUL | [32] | | YPD | mid-log | 14092 | 116802 | 3412 | 4436 | 88 | 47477 | 2015 | 8.2885 | 0.2421 | 0.3148 | 0.0062 | 23.6 |
| LEE2 | [33] | | | mid-log | 28091 | 106796 | 6065 | 17161 | 4634 | 32733 | | 3.8018 | 0.2159 | 0.6109 | 0.1650 | |
| DAV | [34] | | | mid-log | 5991 | 150189 | 2070 | 172 | | 9617 | | 25.0691 | 0.3455 | 0.0287 | | |
| PIC | [35] | | | mid-log | 13872 | 36821 | 9602 | 5334 | 17956 | 16757 | 6516 | 2.6543 | 0.6922 | 0.3845 | 1.2944 | 2.6 |
| LEE | [36] | | | mid-log | 10960 | 251650 | | | | 24363 | | 22.9608 | | | | |
| NEW | [37] | | | mid-log | 11201 | 276461 | 1922 | 4237 | | 11380 | | 24.6818 | 0.1716 | 0.3783 | | |
| GHA | [38] | | | mid-log | 15000 | 135000 | | | 2620 | | 4650 | 9.0000 | | | 0.1747 | |
| DGD | [39] | Haploids and diploids:<br>• YAL6B MATa x Y15969 MATa<br>• BY4741 x BY4742 | minimal | mid-log | 19382 | 71642 | 7086 | 8740 | | 22650 | 4519 | 3.6963 | 0.3656 | 0.4509 | | 5.0 |
| | | Minimum | | | | | | | | | | 0.088 | 0.068 | 0.008 | 0.006 | 1.8 |
| | | Maximum | | | | | | | | | | 75.107 | 3.141 | 3.945 | 1.294 | 45.9 |
| | | Maximum/minimum | | | | | | | | | | 851 | 46 | 468 | 207 | 26 |
| | | Sample size (*n*) | | | | | | | | | | 19 | 12 | 15 | 9 | 9 |

**Table S29. Calculation of total biomass synthesis fluxes generated per 100 intracellular pyruvate molecules according to the metabolic flux distribution described by Fendt and Sauer (2010).**

Note that the numbers presented are also indicated in Figure 6. To register the number, the figure is scanned from left to right, top to bottom. Adapted with modification from Table 5D.3 in [9].

| Biomass precursor | Flux size |
|---|---|
| pentose-5-$P_{cyto}$ | 1 |
| erythrose-4-$P_{cyto}$ | 1 |
| $serine_{cyto}$ | 1 |
| $glycine_{cyto}$ | 1 |
| glucose-6-$P_{cyto}$ | 8 |
| triose-3-$P_{cyto}$ | 2 |
| $PYRUVATE_{mito}$ | 9 |
| $oxaloacetate_{cyto}$ | 5 |
| P-enol-$pyruvate_{cyto}$ | 2 |
| acetyl-$CoA_{mito}$ | 1 |
| $PYRUVATE_{cyto}$ | 1 |
| acetyl-$CoA_{cyto}$ | 6 |
| α-$ketoglutarate_{mito/cyto}$ | 4 |
| **TOTAL** | **42** |

#### Table S30. Endogenous enzymes involved in lactate metabolism in *S. cerevisiae*.

Summarized from [40,41]. "Mito": mitochondrial. "Cyto": cytosolic.

| Substrate | E.C. number | Coding gene | Popular enzyme name | Subcellular location | Abbreviation | References |
|---|---|---|---|---|---|---|
| D-Lactate | ? | ? | Anaerobic D-Lactate dehydrogenase | ? | Anaerobic D-LDH | [42,43] |
| | ? | DLD3? | Aerobic NAD-dependent D-Lactate dehydrogenase – cyto? | Cyto | Aerobic D-LDH? | [44,45] |
| | 1.1.1.28 | DLD2? | Aerobic NAD-dependent D-Lactate dehydrogenase – mito? | Mito | Aerobic D-LDH | [44–46] |
| | 1.1.2.4 | DLD1 | D-Lactate : cytochrome *c* oxidoreductase | Mito | D-LCR | [41,42,47–50] |
| L-Lactate | 1.1.1.27 | ? | NAD-dependent L-Lactate dehydrogenase | Mito | L-LDH | [46] |
| | 1.1.2.3 | CYB2 | L-Lactate : ferricytochrome *c* oxidoreductase | Mito | L-LCR; Cytochrome $b_2$ | [42,51–53] |

**Table S31. Abundance of proteins directly involved in the metabolism of cytosolic L-glycerol-3-P, DL-lactate, isocitrate, and glutamate in *S. cerevisiae* grown on glucose, lactate, or acetate.**

The raw data are obtained from [18]. The strain was BY4742. The minimal medium was based on YNB (with amino acids and ammonium sulfate) and contained 2% of the designated carbon source. The red font indicates flavoproteins.

| Functional category | Protein ID | glucose | lactate | acetate |
|---|---|---|---|---|
| Cytosolic L-glycerol-3-P metabolism | **Gut1** | 1164.110 | 18991.000 | 8395.260 |
| | **Gut2** | 6227.030 | 53304.300 | 54065.100 |
| | **Gpd1** | 6099.240 | 16779.400 | 23807.800 |
| | **Gpd2** | 8319.32 | 7078.61 | 5984.12 |
| Cytosolic DL-lactate metabolism | **Dld1** | 3402.13 | 27609 | 24576 |
| | **Cyb2** | 1163.63 | 26563.5 | 4323.23 |
| | **Dld2** | 5809.800 | 9557.040 | 18902.100 |
| | **Dld3** | 18457.800 | 2527.130 | 3114.960 |
| Cytosolic isocitrate metabolism | **Idp2** | 2804.300 | 44209.300 | 40426.400 |
| | **Icl1** | 4176.900 | 70501.300 | 166401.000 |
| Cytosolic L-glutamate biosynthesis | **Gdh1** | 25262.500 | 4686.480 | 2803.260 |
| | **Gdh3** | 7280.910 | 9842.400 | 2468.980 |
| | **Gln1** | 15079.600 | 6318.060 | 20284.400 |
| | **Glt1** | 65960.700 | 21324.500 | 7120.990 |

**Figure S17. Reactions and enzymes involved in the biosynthesis of cytosolic L-glutamate in *S. cerevisiae*.**

**A**, Gdh1/3-catalyzed reaction. **B**, Gln1-catalyzed reaction. **C**, Glt1-catalyzed reaction. Source of figures: *Saccharomyces* Genome Database (SGD, https://www.yeastgenome.org/).

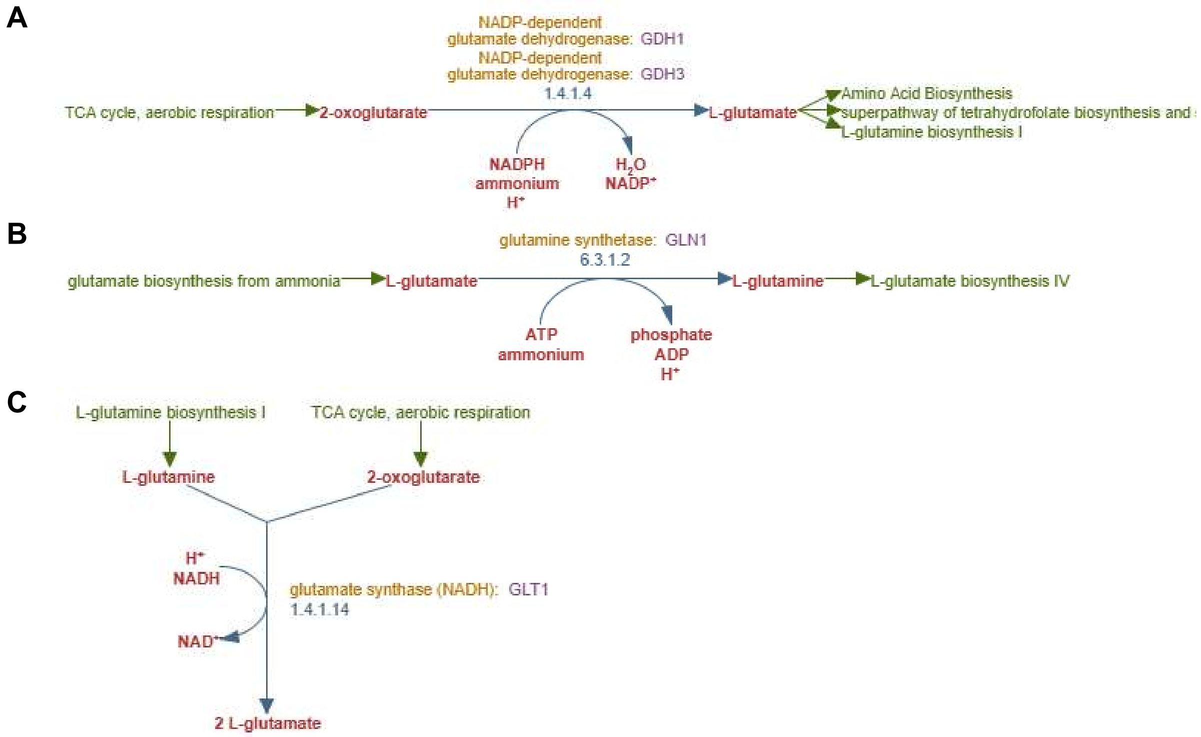

##### Figure S18. Reactions and enzymes involved in the glyoxylate cycle of *S. cerevisiae.*

Source of figure: *Saccharomyces* Genome Database (SGD, https://www.yeastgenome.org/).

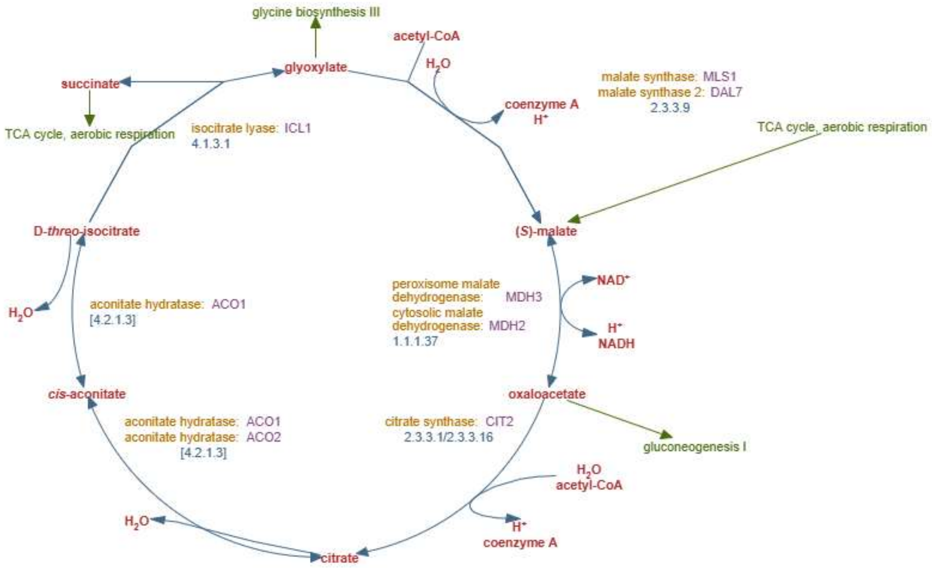

**Table S32. Kinetic parameters for isocitrate-processing enzymes in *S. cerevisiae*.**

| Enzyme | Enzyme | *S. cerevisiae* strain | Substrate | *Km* | Reference |
|---|---|---|---|---|---|
| Isocitrate lyase | **Icl1** | Fleischman's bakers' yeast | DL-isocitrate | 2.4 mM | 54 |
| | | | $\mathrm{L_s}$-isocitrate | 1.2 mM | |
| | | Unspecified diploid | DL-isocitrate | 1.3 mM | 55 |
| | | G-517 | *threo*-$\mathrm{D_s}$-isocitrate | 1.4 mM | 56 |
| NADP-isocitrate dehydrogenase | **Idp2**? Idp1 (mito)? | Unspecified diploid | DL-isocitrate | 0.04 mM | 55 |
| | Idp1 (mito) | S173-6B transformed with the relevant plasmid | D-isocitrate | 0.03 mM | 57 |
| | **Idp2** | | D-isocitrate | 0.22 mM | |
| | | | $NADP^+$ | 0.02 mM | |
| | | | α-ketoglutarate | 0.20 mM | |
| | | | NADPH | 0.04 mM | |

#### Table S33. Abundance of glyoxylate cycle enzymes in *S. cerevisiae* grown on glucose, lactate, and acetate.

Statistical significance (*p*-value) is calculated using the TTEST function (two-tailed distribution, two-sample equal variance (homoscedastic)) in Microsoft Excel.

| Protein ID | glucose | lactate | acetate | lactate/glucose | Apparent group # | *p*-value group 1 vs. group 2 |
|---|---|---|---|---|---|---|
| **Icl1** | 4176.900 | 70501.300 | 166401.000 | 16.9 | 1 | 0.002 |
| **Mls1** | 1062.940 | 28982.600 | 52768.900 | 27.3 | | |
| **Dal7** | 669.271 | 18324.000 | 35959.200 | 27.4 | | |
| **Mdh2** | 1777.710 | 25024.100 | 42071.600 | 14.1 | | |
| **Mdh3** | 5097.710 | 11472.000 | 14663.300 | 2.3 | 2 | |
| **Cit2** | 2818.370 | 16233.600 | 85489.100 | 5.8 | | |
| **Aco1** | 14846.300 | 59345.300 | 122999.000 | 4.0 | | |
| **Aco2** | 12662.200 | 9624.670 | 6659.830 | 0.8 | | |

#### Figure S19. Growth profiles of *S. cerevisiae* D 261 on D-lactate at varying concentrations.

Based on a graphical analysis of Fig. 10 in [40]. The turbidity of culture in 0 mM D-lactate was not reported but is logically assumed to be zero.

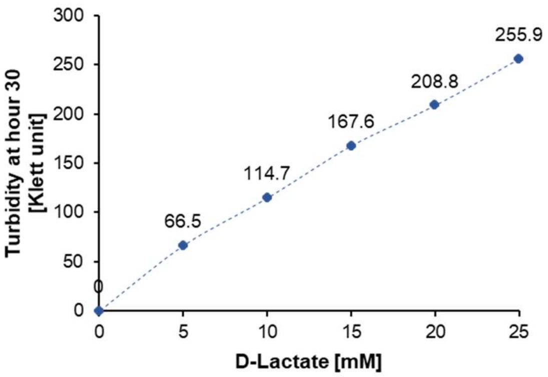

### Supplemental Information References

1. U.S. Department of Energy (2024). 2023 Billion-Ton Report: An Assessment of U.S. Renewable Carbon Resources https://doi.org/10.23720/BT2023/2316165.
2. Wu, H., Scheve, T., Dalke, R., Holtzapple, M., and Urgun-Demirtas, M. (2023). Scaling up carboxylic acid production from cheese whey and brewery wastewater via methane-arrested anaerobic digestion. Chem. Eng. J. *459*, 140080.
3. Skaggs, R.L., Coleman, A.M., Seiple, T.E., and Milbrandt, A.R. (2018). Waste-to-Energy biofuel production potential for selected feedstocks in the conterminous United States. Renew. Sustain. Energy Rev. *82*, 2640–2651.
4. Walkling, C.J., Zhang, D.D., and Harvey, B.G. (2024). Extended fuel properties of sustainable aviation fuel blends derived from linalool and isoprene. Fuel *356*, 129554.
5. Geiselman, G.M., Kirby, J., Landera, A., Otoupal, P., Papa, G., Barcelos, C., Sundstrom, E.R., Das, L., Magurudeniya, H.D., and Wehrs, M. (2020). Conversion of poplar biomass into high-energy density tricyclic sesquiterpene jet fuel blendstocks. Microb. Cell Fact. *19*, 1–16.
6. Monod, J. (1949). The growth of bacterial cultures. Annu. Rev. Microbiol. *3*, 371–394.
7. Van Dijken, J.P., Bauer, J., Brambilla, L., Duboc, P., Francois, J.M., Gancedo, C., Giuseppin, M.L.F., Heijnen, J.J., Hoare, M., and Lange, H.C. (2000). An interlaboratory comparison of physiological and genetic properties of four Saccharomyces cerevisiae strains. Enzyme Microb. Technol. *26*, 706–714.
8. Boubekeur, S., Camougrand, N., Bunoust, O., Rigoulet, M., and Guérin, B. (2001). Participation of acetaldehyde dehydrogenases in ethanol and pyruvate metabolism of the yeast Saccharomyces cerevisiae. Eur. J. Biochem. *268*, 5057–5065.
9. Nurani, W. (2023). Chapter 5. Impacts of glycerol cofeeding and precultivation on acetate on the growth of S. cerevisiae on various carbon sources. In Physiological augmentation strategies to support the production of non-native secondary metabolites in Saccharomyces cerevisiae. PhD thesis (Technical University of Denmark), pp. 139–189.
10. Babel, W. (2009). The auxiliary substrate concept: from simple considerations to heuristically valuable knowledge. Eng. Life Sci. *9*, 285–290.
11. Gientka, I., Gadaszewska, M., Błażejak, S., Kieliszek, M., Bzducha-Wróbel, A., Stasiak-Różańska, L., and Kot, A.M. (2017). Evaluation of lipid biosynthesis ability by Rhodotorula and Sporobolomyces strains in medium with glycerol. Eur. Food Res. Technol. *243*, 275–286.
12. Lubuta, P., Workman, M., Kerkhoven, E.J., and Workman, C.T. (2019). Investigating the influence of glycerol on the utilization of glucose in Yarrowia lipolytica using RNA-Seq-based transcriptomics. G3 Genes, Genomes, Genet. *9*, 4059–4071.
13. Lages, F., Silva-Graça, M., and Lucas, C. (1999). Active glycerol uptake is a mechanism underlying halotolerance in yeasts: a study of 42 species. Microbiology *145*, 2577–2585.
14. Workman, M., Holt, P., and Thykaer, J. (2013). Comparing cellular performance of Yarrowia lipolytica during growth on glucose and glycerol in submerged cultivations. Amb Express *3*, 58.
15. Fendt, S.-M., and Sauer, U. (2010). Transcriptional regulation of respiration in yeast metabolizing differently repressive carbon substrates. BMC Syst. Biol. *4*, 12.
16. Crespo, J.L., and Hall, M.N. (2002). Elucidating TOR signaling and rapamycin action: lessons from Saccharomyces cerevisiae. Microbiol. Mol. Biol. Rev. *66*, 579–591.
17. de Vries, S., and Marres, C.A.M. (1987). The mitochondrial respiratory chain of yeast. Structure and biosynthesis and the role in cellular metabolism. Biochim. Biophys. Acta (BBA)-Reviews Bioenerg. *895*, 205–239.
18. Paulo, J.A., O'Connell, J.D., Everley, R.A., O'Brien, J., Gygi, M.A., and Gygi, S.P. (2016). Quantitative mass spectrometry-based multiplexing compares the abundance of 5000 S. cerevisiae proteins across 10 carbon sources. J. Proteomics *148*, 85–93.
19. Ho, B., Baryshnikova, A., and Brown, G.W. (2018). Unification of protein abundance datasets yields a quantitative Saccharomyces cerevisiae proteome. Cell Syst. *6*, 192–205.
20. Lahtvee, P.-J., Sánchez, B.J., Smialowska, A., Kasvandik, S., Elsemman, I.E., Gatto, F., and Nielsen, J. (2017). Absolute quantification of protein and mRNA abundances demonstrate variability in gene-specific translation efficiency in yeast. Cell Syst. *4*, 495–504.
21. Nagaraj, N., Kulak, N.A., Cox, J., Neuhauser, N., Mayr, K., Hoerning, O., Vorm, O., and Mann, M. (2012). System-wide perturbation analysis with nearly complete coverage of the yeast proteome by single-shot ultra HPLC runs on a bench top Orbitrap. Mol. Cell. Proteomics *11*, M111-013722.
22. Lu, P., Vogel, C., Wang, R., Yao, X., and Marcotte, E.M. (2007). Absolute protein expression profiling estimates the relative contributions of transcriptional and translational regulation. Nat. Biotechnol. *25*, 117–124.
23. Webb, K.J., Xu, T., Park, S.K., and Yates III, J.R. (2013). Modified MuDPIT separation identified 4488 proteins in a system-wide analysis of quiescence in yeast. J. Proteome Res. *12*, 2177–2184.
24. Thakur, S.S., Geiger, T., Chatterjee, B., Bandilla, P., Fröhlich, F., Cox, J., and Mann, M. (2011). Deep and highly sensitive proteome coverage by LC-MS/MS without prefractionation. Mol. Cell. Proteomics *10*, M110-003699.
25. Lawless, C., Holman, S.W., Brownridge, P., Lanthaler, K., Harman, V.M., Watkins, R., Hammond, D.E.,

Miller, R.L., Sims, P.F.G., and Grant, C.M. (2016). Direct and absolute quantification of over 1800 yeast proteins via selected reaction monitoring. Mol. Cell. Proteomics *15*, 1309–1322.

26. Peng, M., Taouatas, N., Cappadona, S., Van Breukelen, B., Mohammed, S., Scholten, A., and Heck, A.J.R. (2012). Protease bias in absolute protein quantitation. Nat. Methods *9*, 524–525.
27. Dénervaud, N., Becker, J., Delgado-Gonzalo, R., Damay, P., Rajkumar, A.S., Unser, M., Shore, D., Naef, F., and Maerkl, S.J. (2013). A chemostat array enables the spatio-temporal analysis of the yeast proteome. Proc. Natl. Acad. Sci. *110*, 15842–15847.
28. Chong, Y.T., Koh, J.L.Y., Friesen, H., Duffy, S.K., Cox, M.J., Moses, A., Moffat, J., Boone, C., and Andrews, B.J. (2015). Yeast proteome dynamics from single cell imaging and automated analysis. Cell *161*, 1413–1424.
29. Breker, M., Gymrek, M., and Schuldiner, M. (2013). A novel single-cell screening platform reveals proteome plasticity during yeast stress responses. J. Cell Biol. *200*, 839–850.
30. Mazumder, A., Pesudo, L.Q., McRee, S., Bathe, M., and Samson, L.D. (2013). Genome-wide single-cell-level screen for protein abundance and localization changes in response to DNA damage in S. cerevisiae. Nucleic Acids Res. *41*, 9310–9324.
31. Tkach, J.M., Yimit, A., Lee, A.Y., Riffle, M., Costanzo, M., Jaschob, D., Hendry, J.A., Ou, J., Moffat, J., and Boone, C. (2012). Dissecting DNA damage response pathways by analysing protein localization and abundance changes during DNA replication stress. Nat. Cell Biol. *14*, 966–976.
32. Kulak, N.A., Pichler, G., Paron, I., Nagaraj, N., and Mann, M. (2014). Minimal, encapsulated proteomic-sample processing applied to copy-number estimation in eukaryotic cells. Nat. Methods *11*, 319–324.
33. Lee, M.V., Topper, S.E., Hubler, S.L., Hose, J., Wenger, C.D., Coon, J.J., and Gasch, A.P. (2011). A dynamic model of proteome changes reveals new roles for transcript alteration in yeast. Mol. Syst. Biol. *7*, 514.
34. Davidson, G.S., Joe, R.M., Roy, S., Meirelles, O., Allen, C.P., Wilson, M.R., Tapia, P.H., Manzanilla, E.E., Dodson, A.E., and Chakraborty, S. (2011). The proteomics of quiescent and nonquiescent cell differentiation in yeast stationary-phase cultures. Mol. Biol. Cell *22*, 988–998.
35. Picotti, P., Bodenmiller, B., Mueller, L.N., Domon, B., and Aebersold, R. (2009). Full dynamic range proteome analysis of S. cerevisiae by targeted proteomics. Cell *138*, 795–806.
36. Lee, M., Kim, B., Choi, H., Ryu, M., Kim, S., Kang, K., Cho, E., Youn, H., Huh, W., and Kim, S. (2007). Global protein expression profiling of budding yeast in response to DNA damage. Yeast *24*, 145–154.
37. Newman, J.R.S., Ghaemmaghami, S., Ihmels, J., Breslow, D.K., Noble, M., DeRisi, J.L., and Weissman, J.S. (2006). Single-cell proteomic analysis of S. cerevisiae reveals the architecture of biological noise. Nature *441*, 840–846.
38. Ghaemmaghami, S., Huh, W.-K., Bower, K., Howson, R.W., Belle, A., Dephoure, N., O'Shea, E.K., and Weissman, J.S. (2003). Global analysis of protein expression in yeast. Nature *425*, 737–741.
39. De Godoy, L.M.F., Olsen, J. V, Cox, J., Nielsen, M.L., Hubner, N.C., Fröhlich, F., Walther, T.C., and Mann, M. (2008). Comprehensive mass-spectrometry-based proteome quantification of haploid versus diploid yeast. Nature *455*, 1251–1254.
40. Pajot, P., and Claisse, M.L. (1974). Utilization by Yeast of d-Lactate and l-Lactate as Sources of Energy in the Presence of Antimycin A. FEBS J. *49*, 275–285.
41. Lodi, T., and Ferrero, I. (1993). Isolation of the DLD gene of Saccharomyces cerevisiae encoding the mitochondrial enzyme D-lactate ferricytochrome c oxidoreductase. Mol. Gen. Genet. MGG *238*, 315–324.
42. Labeyrie, F., and Slonimski, P.P. (1964). Mode daction des lacticodeshydrogenases liees aux systemes flavinique et cytochromique. Bull. Soc. Chim. Biol. (Paris). *46*, 1793-+.
43. Labeyrie, F., Slonimski, P.P., and Naslin, L. (1959). Sur la différence de stéréospécificité entre la déshydrogénase lactique extraite de la levure anaérobie et celle extraite de la levure aérobie. Biochim. Biophys. Acta *34*, 262–265.
44. Chelstowska, A., Liu, Z., Jia, Y., Amberg, D., and Butow, R.A. (1999). Signalling between mitochondria and the nucleus regulates the expression of a new d-lactate dehydrogenase activity in yeast. Yeast *15*, 1377–1391.
45. Becker-Kettern, J., Paczia, N., Conrotte, J.-F., Kay, D.P., Guignard, C., Jung, P.P., and Linster, C.L. (2016). Saccharomyces cerevisiae forms D-2-hydroxyglutarate and couples its degradation to D-lactate formation via a cytosolic transhydrogenase. J. Biol. Chem. *291*, 6036–6058.
46. Genga, A.M., Tassi, F., Lodi, T., and Ferrero, I. (1983). Mitochondrial NAD, L-lactate dehydrogenase and NAD, D-lactate dehydrogenase in the yeast Saccharomyces cerevisiae. Microbiologica *6*, 1–8.
47. Nygaard, A.P. (1960). Lactic dehydrogenase of yeast: III. A comparative study of the kinetic properties and the stability of two isolated forms of the enzyme. Biochim. Biophys. Acta *40*, 85–92.
48. Nygaard, A.P. (1961). Induction of d (-)-and l (+)-Lactic Cytochrome c Reductase in Yeast. J. Biol. Chem. *236*, 1585–1588.
49. Lodi, T., Alberti, A., Guiard, B., and Ferrero, I. (1999). Regulation of the Saccharomyces cerevisiae DLD1 gene encoding the mitochondrial protein D-lactate ferricytochrome c oxidoreductase by HAP1 and HAP2/3/4/5. Mol. Gen. Genet. MGG *262*, 623–632.
50. Rojo, E.E., Guiard, B., Neupert, W., and Stuart, R.A. (1998). Sorting of D-lactate dehydrogenase to the inner membrane of mitochondria: analysis of topogenic signal and energetic requirements. J. Biol. Chem. *273*, 8040–8047.
51. Bernheim, F. (1928). The specificity of the dehydrases: The separation of the citric acid dehydrase from liver and of the lactic acid dehydrase from yeast. Biochem. J. *22*, 1178.
